# Synergistic retinal UCHL1 dysregulation and synaptic vulnerability reflect Alzheimer’s disease severity

**DOI:** 10.64898/2026.02.03.703406

**Authors:** Altan Rentsendorj, Jean-Philippe Vit, Alexandre Hutton, Yosef Koronyo, Saba Shahin, Edward Robinson, Ernesto Barron, Anthony Rodriguez, Bhakta Prasad Gaire, Ousman Jallow, Julia Sheyn, Miyah Rene Davis, Alexander V. Ljubimov, Stuart L. Graham, Vivek K. Gupta, Mehdi Mirzaei, Debra Hawes, Keith L. Black, Lon S. Schneider, Alfredo A. Sadun, Jesse G. Meyer, Dieu-Trang Fuchs, Maya Koronyo-Hamaoui

## Abstract

Synaptic failure predicts cognitive decline in Alzheimer’s disease (AD), yet its impact and molecular drivers in the human retina remain unclear. Leveraging the retina as an accessible central nervous system (CNS) proxy, we integrated spatially resolved histopathology of retinal cross-sections with ultrastructural, proteomic, and biochemical profiling across independent postmortem cohorts spanning normal cognition, mild cognitive impairment due to AD (MCI), and AD dementia. We uncover early, progressive degeneration of excitatory glutamatergic synapses, evidenced by losses of presynaptic vesicular glutamate transporter 1 (VGLUT1) and synaptophysin, and postsynaptic density protein 95 (PSD95) and N-methyl-D-aspartate receptor subunit 2A (NMDAR2A), accompanied by disruption of synaptic ultrastructure. Retinal synaptic loss tightly associates with local accumulation of amyloid-β 42 (Aβ42) and immature tau species, heightened oxidative stress, and upregulation of the Aβ-binding death receptor p75 neurotrophin receptor (p75NTR). Notably, the synapse-enriched deubiquitinase ubiquitin C-terminal hydrolase L1 (UCHL1) is profoundly dysregulated, correlates with synaptic integrity and cognition, and emerges as the strongest retinal predictor of Braak stage and cognitive status in multivariable machine-learning models. Together, these findings position retinal Aβ/p75NTR-mediated UCHL1 imbalance as a proteostasis–synapse mechanistic hub and candidate biomarker reflecting AD severity.

## Introduction

Synaptic dysfunction is an early and defining process in the brains of Alzheimer’s disease (AD) patients and represents the most reliable predictor of cognitive decline^1–3^. Traditionally characterized by amyloid β-protein (Aβ) plaques and neurofibrillary tangles (NFTs) composed of hyperphosphorylated (p)tau protein^4–9^, AD is increasingly recognized as a disorder driven by early synaptic dysfunction, with deficits in glutamatergic signaling commonly preceding neuronal loss^2,10–17^. We previously found that overnight exposure to fibrillar, and moreover, oligomeric Aβ_42_ aggregates caused 50-70% loss of glutamatergic pre-and post-synaptic puncta in mouse primary cortical neuronal cultures^18^. However, despite decades of study, the mechanisms linking Aβ and tau accumulation to synaptic collapse and neurodegeneration remain unclear, underscoring the need for accessible biomarkers that faithfully reflect these pathological processes.

The retina, an embryological outgrowth of the brain, provides a uniquely accessible site into central nervous system (CNS) pathology. Increasing evidence demonstrates that AD hallmark pathologies—including Aβ deposits, tau inclusions, gliosis, and vascular changes—are detectable in the human retina and correlate with cerebral pathology and cognitive decline^19–39^. Advanced retinal imaging modalities, such as hyperspectral imaging and scanning laser ophthalmoscopy, have further detected Aβ and vascular dysfunction in living patients^27,40–51^. However, whether retinal synapses undergo degeneration that parallels brain neurodegeneration in AD, and which molecular mechanisms govern this process, remain largely unexplored.

One pathway strongly implicated in AD pathogenesis involves disruption of the ubiquitin–proteasome system (UPS), which contributes to Aβ and tau accumulation, oxidative stress, and synaptic loss^52–59^. Ubiquitin carboxyl-terminal hydrolase L1 (UCHL1), a neuron-enriched deubiquitinating enzyme and key component of the UPS, is essential for maintaining proteostasis, synaptic integrity, and neuronal survival^60–63^. In the brain, UCHL1 facilitates clearance of misfolded proteins and prevents aggregation of toxic Aβ42 and ptau^62,64–66^. Reduced UCHL1 expression in AD cortex and hippocampus correlates with synaptic loss and cognitive decline^56^. However, UCHL1 expression and its potential contribution to synaptic pathology have not been investigated in the AD retina. Another AD-relevant neurodegenerative pathway involves signaling via the p75 neurotrophin receptor (p75NTR). Initially characterized as a death receptor, p75NTR regulates axonal pruning, synaptic plasticity, and apoptosis in the CNS^67–71^. In brains from AD donors and rodent models, Aβ was shown to bind to p75NTR, driving neuritic degeneration, tau hyperphosphorylation, oxidative stress, and neuronal loss^72–77^. Furthermore, increased retinal p75NTR has been reported in experimental retinopathy models^78–80^; however, its relevance to AD retinal pathology and its potential relationship to synaptic failure remain unknown.

Here, we integrate spatially resolved histological quantification of supero- and infero-temporal (ST, IT) retinal cross-sections with ultrastructural and proteomic analyses to interrogate synaptic integrity, UPS function, and death receptor signaling across multiple independent human cohorts encompassing normal cognition (NC) controls, mild cognitive impairment (MCI due to AD), and AD dementia. We focus on excitatory glutamatergic synapses, quantifying presynaptic vesicular glutamate transporter 1 (VGLUT1) and synaptophysin (SYP), as well as postsynaptic postsynaptic density protein 95 (PSD95) and N-methyl-D-aspartate receptor subunit 2A (NMDAR2A). These proteins collectively orchestrate glutamate packaging and release, receptor-mediated excitatory transmission, and postsynaptic scaffolding, and their disruption is strongly implicated in cortical synaptic failure in AD^81–84^. Comprehensive proteomic, histological, biochemical, and transmission electron microscopy analyses reveal widespread degeneration of pre- and postsynaptic proteins, accompanied by ultrastructural collapse of synaptic architecture, early photoreceptor vulnerability, and marked dysregulation of UCHL1 in the retinas of MCI and AD patients. These changes strongly associate with retinal Aβ_42_ accumulation, immature tau species, oxidative stress, p75NTR signaling, and cognitive impairment, mirroring mechanisms described in the brain. Multivariable machine-learning models further identify UCHL1 as the strongest retinal predictor of Braak stage and cognitive status. Collectively, our findings establish the retina as a site of early synaptic and proteostatic collapse in AD and highlight retinal UCHL1 dysregulation as both a mechanistic hub and a candidate biomarker linking hallmark pathology to neuronal vulnerability.

## Results

To investigate synaptic integrity and associated neurodegeneration in postmortem retinas from patients with early and advanced-stage AD, we analyzed the spatiotemporal distribution and burden of synaptic proteins in superior- and inferior-temporal (ST/IT) retinal cross-sections from a cohort of 53 individuals comprising NC controls (n=21, mean age 82.6 ± 10.5 years, 11 females/10 males), MCI due-to-AD (n=11, mean age 89.3 ± 5.2 years, 7 females/4 males), and AD dementia (n=21, mean age 86.2 ± 8.9 years, 13 females/8 males). Age, sex, and post-mortem interval (PMI) did not differ significantly among the three diagnostic groups, and these variables were controlled for in each retinal biomarker subset, minimizing potential confounding effects. An additional cohort was used for mass spectrometry-based proteomics, Western Blot, and ELISA assays to assess dysregulation of synaptic and AD-related proteins in the temporal retina of AD patients and NC controls (n=16). Demographic, clinical, and neuropathological data for the cohorts used in histological and protein analyses is detailed in Figure 1A and Table 1 (individual human donors are listed in Table S1).

**Figure 1.**
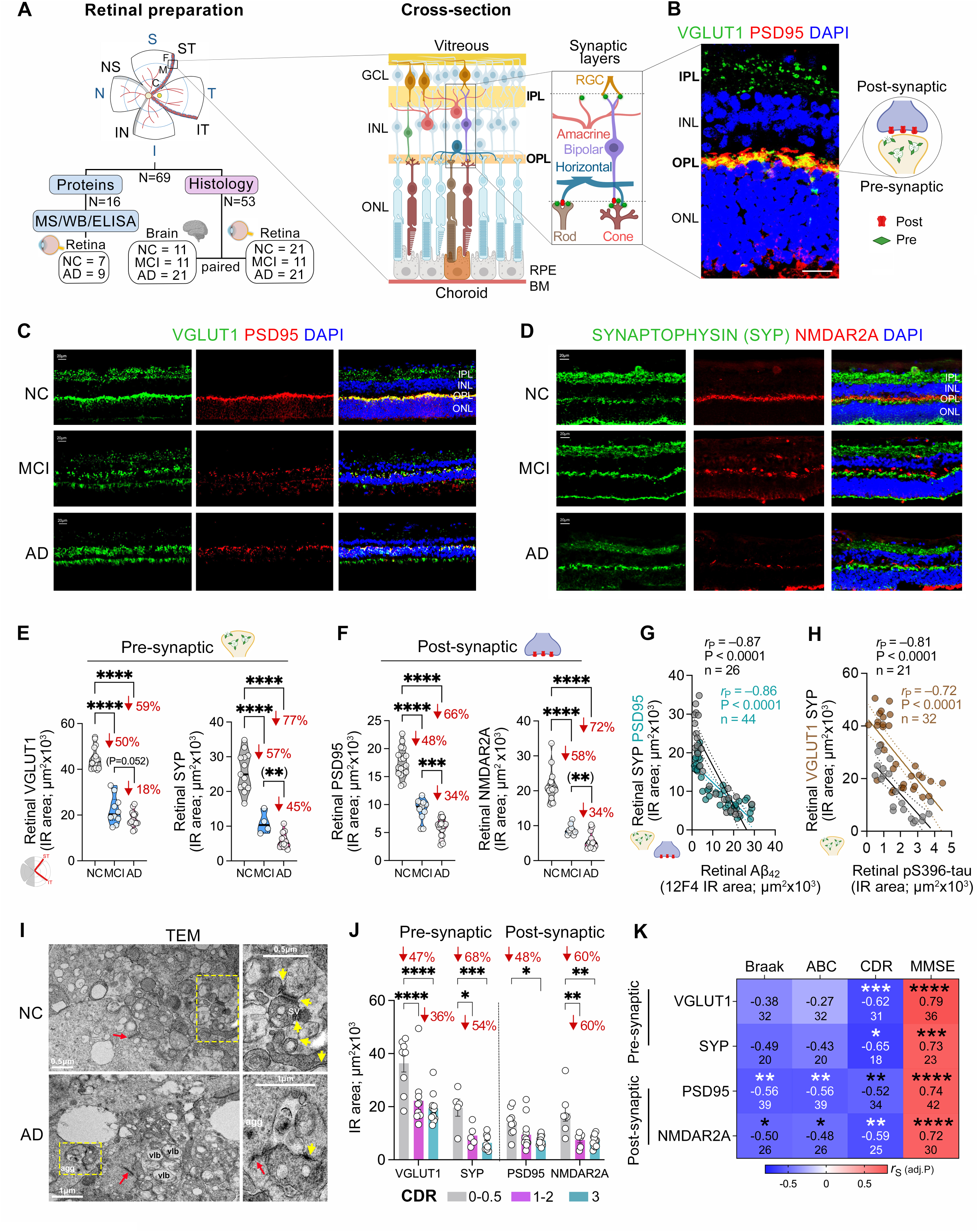
Synaptic loss in the retinas of MCI and AD patients and its relationship with core-AD pathologies and disease status. (**A**) Schematic of analyzed retinal cross-sections from individuals with normal cognition (NC), patients with mild cognitive impairment (MCI due to AD) or with Alzheimer’s disease (AD) in predefined geometrical regions within the superior- and inferior-temporal (ST/IT) retina (red strips), further subdivided into central (C), mid-periphery (M) and far-periphery (F) subregions. Retinal layers and basic synaptic architecture in the inner and outer plexiform layers (IPL and OPL), including pre- and post-synaptic terminals, are illustrated. The flow diagram outlines allocation of human donor eyes for histopathological and protein/ biochemical analyses, with subject numbers indicated. (**B**) Representative confocal image of retina immunolabeled for a pre- and post-synaptic marker, VGLUT1 (green) and PSD95 (red), showing strong synaptic labeling in the IPL and OPL. Nuclei are stained with DAPI (blue). Scale bar: 20 µm. (**C-D**) Representative micrographs/images of retinas from MCI, AD and NC donors, immunostained for pre-synaptic (VGLUT1 and synaptophysin SYP, green) and post-synaptic (PSD95 and NMDAR2A, red) markers. Nuclei are stained with DAPI (blue). Scale bars: 20 µm. (**E-F**) Violin plots showing quantitative immunoreactive (IR) area of retinal pre-synaptic (VGLUT1, SYP) and post-synaptic (PSD95, NMDAR2A) markers in MCI (n = 4-11) and AD (n = 12-20) patients, compared to age and sex-matched NC controls (n = 13-18). (**G-H**) Pearson’s correlation coefficient (*r*_P_) analysis showing a strong inverse relationship between (**G**) retinal PSD95, SYP and retinal Aβ_42_ (12F4) burden, and (**H**) retinal VGLUT1, SYP and retinal pS396-tau burden. (**I**) Transmission electron microscopy (TEM) images of OPL region from NC and AD retinas, labeled with pre- and post-synaptic markers, VGLUT1 and PSD95, respectively. Yellow arrowheads show synaptic clefts between pre- and post-synaptic terminals. Pre-synaptic terminals containing electron dense synaptic vesicles (SVs) are marked. Red arrows highlight membrane-docked pre-synaptic ribbons, which are preserved in NC retinas but are detached and floating in AD retinas. agg, electron dense aggregate; vlb, blebbing leakage. Scale bar: 0.5- 1 µm. (**J**) Quantitative analysis of retinal VGLUT1, SYP, PSD95 and NMDAR2A, stratified by clinical dementia rating (CDR) scores (CDR 0-0.5 = 5-9; CDR 1-2 = 5-12; CDR 3 = 8-13). (**K**) Heatmap showing Spearman’s correlation coefficients (*r*_S_) and significance (adjusted P values with Holm-Šídák multiple-comparison correction method for each retinal neuronal marker against 4 brain pathology and cognitive scores) between retinal synaptic markers and brain AD pathology (BRAAK, ABC scores) and cognitive performance/status (MMSE, CDR). Violin plots show individual subjects (circles) with lower, median and upper quartiles. Bar graph displays group means ± SEMs., *P < 0.05, ** P < 0.01, *** P < 0.001, **** P < 0.0001, by one-way ANOVA with Tukey’s post-hoc multiple comparison test. Percentage (%) changes are shown in red. IPL, inner plexiform layer; INL, inner nuclear layer; OPL, outer plexiform layer; ONL, outer nuclear layer. Schematic illustrations were created using Biorender.com.

**Table 1.**
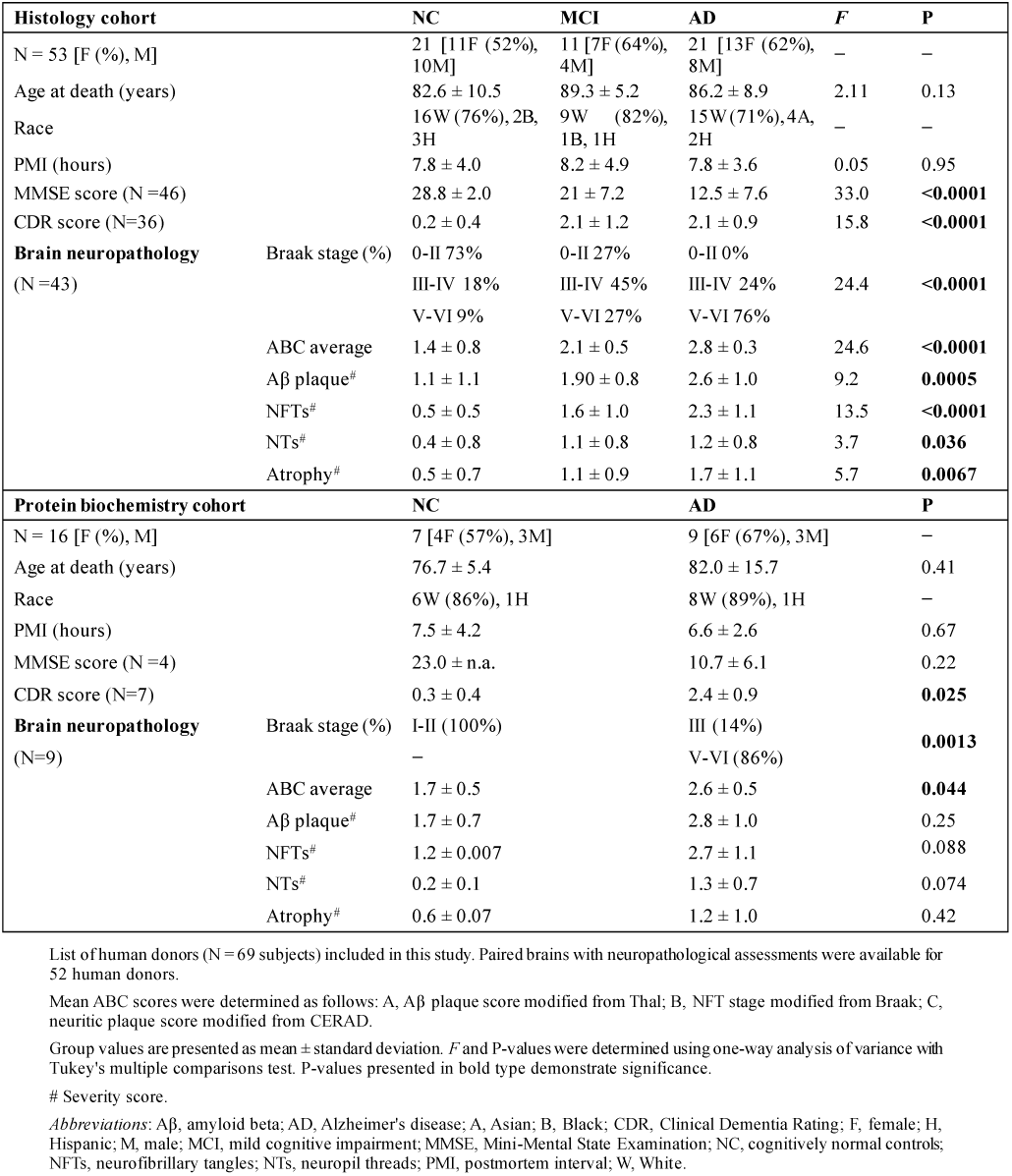
Demographic and neuropathological data on human brains and retinas in this study.

### Early and profound retinal synaptic degeneration strongly links to cognitive impairment in AD

To determine the impact of AD on retinal synaptic integrity, we initially conducted a histological analysis of glutamatergic pre- (VGLUT1) and post- (PSD95) synaptic proteins in retinal synapse-rich inner and outer plexiform layers (IPL/OPL), in a cohort of AD and MCI patients compared with age- and sex-matched NC controls (n=51; Figure 1A-B). We further analyzed the presynaptic vesicle glycoprotein synaptophysin (SYP) and the postsynaptic glutamatergic receptor (NMDAR2A) in a subset of these subjects (n=29-36). Synaptic marker expression was next correlated with disease status in respective brain regions (n=43). Immunohistochemical (IHC) analysis of these pre- and post-synaptic markers showed early and progressive synaptic protein loss in MCI and AD retinas (Figure 1C and 1D; extended data in Figure S1A and S1B), with distinct distribution patterns for pre-synapses (IPL and OPL) and post-synapses (OPL dominant). Quantitative IHC analysis revealed highly significant reductions in pre-synaptic markers (VGLUT1 and SYP) by 50-57% in MCI retinas and 59-77% in AD retinas compared to NC controls (P < 0.0001, Figure 1E). Post-synaptic marker levels (PSD95 and NMDAR2A) followed similar trajectories, with marked decreases of 48–58% and 66–72% in MCI and AD retinas, respectively (P < 0.0001, Figure 1F). A detailed geometrical analysis of retinal synaptic markers across central (C), mid-peripheral (M), and far-peripheral (F) subregions demonstrated synaptic losses that were not uniformly distributed across the MCI/AD retina (Figure S1C-F). Indeed, the mid- and far-peripheral retina exhibited the most pronounced degeneration in AD, particularly for synaptophysin (Figure S1D). This spatial pattern aligns with prior evidence of peripheral-dominant retinal amyloidosis and tauopathy distribution^28,37^, suggesting localized synaptic vulnerability to AD pathology.

To test this, we evaluated the relationship between synaptic integrity and Aβ/tau pathology in the same retinal regions (representative images in Figure S1G and S1H); all multivariable correlations reported hereafter are presented as adjusted *P* values (P*_adj._*). Pearson correlation coefficient (*r*_P_) analysis of retinal synaptic markers and retinal amyloidosis and tau isoforms (Table 2), revealed that all pre- and post-synaptic proteins, particularly synaptophysin and PSD95, were very closely associated with and highly vulnerable to Aβ_42_ levels in the same regions [*r* = –0.86-(–0.87), P*_adj._* < 0.0001; Figure 1G]. Synaptophysin was also tightly associated with Aβ oligomers in the retina (*r* = –0.83, P*_adj._* < 0.01; Table 2). With respect to retinal tauopathy, synaptophysin was very strongly correlated and susceptible to pathogenic tau species, particularly pS396-tau isoforms (*r* = –0.81, P*_adj._* < 0.0001; Figure 1H). In addition, retinal VGLUT1 levels were negatively correlated with Aβ_42_ and pS396-tau burden [*r* = –0.72-(–0.81), P*_adj._* < 0.0001; Table 2 and Figure 1H]. Most immature retinal tau species, including CitR_209_-tau, pS396-tau, and T22_ tau oligomers, exhibited strong associations with both pre- and postsynaptic markers (Table 2), whereas AT8_ p-tau showed significant correlations only with synaptophysin and PSD95 (*r* = –0.60-(–0.68), P*_adj._* < 0.05-0.01). Of note, mature forms of abnormal tau, including PHF-tau and MC1^+^ NFTs, did not correlate with any synaptic proteins in the retina (Table 2). These findings support a model in which retinal synaptic degeneration is driven by local Aβ and tau aggregation, mirroring well-characterized mechanisms in the AD brain. Interestingly, although retinal tauopathy colocalized with sites of synaptic loss in patients with MCI and AD, retinal Aβ_42_ was associated with the most detrimental impact on synaptic integrity.

**Table 2.**
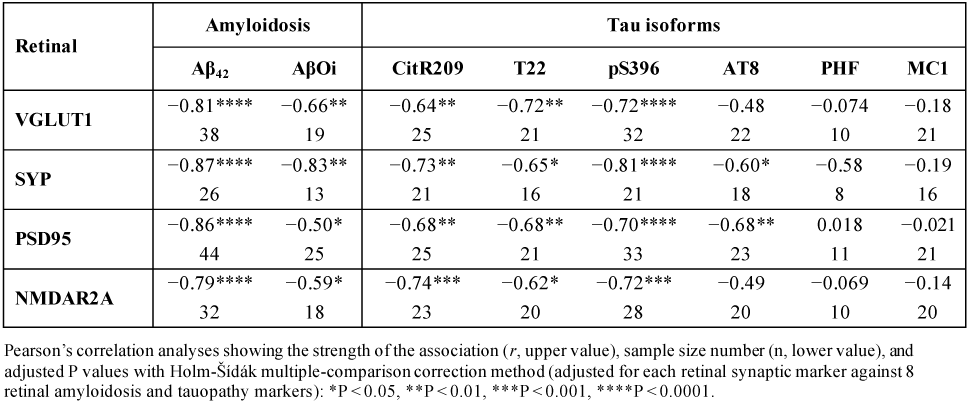
Correlations between retinal synaptic markers and retinal AD pathology.

Ultrastructural analysis of the IPL and OPL layers using transmission electron microscopy (TEM), following immunolabeling with pre-synaptic (VGLUT1) and post-synaptic (PSD95) markers, further confirmed synaptic morphological disruption and loss of synaptic integrity in the AD retina (Figures 1I). In NC control retinas, synapses displayed well-preserved architecture (yellow arrows), with intact clefts, membrane-anchored pre-synaptic ribbons (red arrows), and pre-synaptic terminals densely packed with electron-dense synaptic vesicles (SVs) aligned with clearly defined post-synaptic terminals (Figure 1I, upper panel). In contrast, AD retinas exhibited reduced synaptic density or strength, as well as severe ultrastructural synaptic abnormalities, including darkened and electron-dense aggregates (agg), fragmented and mis-localized free floating pre-synaptic ribbons within the cytoplasm, meaning, there was a loss of membrane-anchored ribbon structures, vesicular leakage, and cellular blebbing (vlb) (Figure 1I, lower panel). These morphological abnormalities are consistent with advanced synaptic degeneration and loss of structural integrity.

Next, we determined whether retinal synaptic loss reflected overall AD severity and stage, as indexed by brain pathology, including Braak stage and ABC scores, and cognitive status as assessed by the clinical dementia rating (CDR) and the mini-mental state examination (MMSE; Figure 1J and 1K; extended data in Figure S1I and S1J, and Tables S2-S5). Stratification by CDR categories revealed progressive reductions in synaptic markers, VGLUT1 by 36–47%, synaptophysin by 54–68%, PSD95 by 48%, and NMDAR2A by 60% in individuals with CDR 1–2 or CDR 3 as compared to individuals with CDR 0 (Figure 1J and Table S2). Stratification of our cohort per Braak stages, revealed similar patterns of significant reductions in retinal VGLUT1 (34–41%) and SYP (50–63%) in individuals with Braak stage III–VI compared to stage 0-II, whereas retinal PSD95 and NMDAR2A were significantly reduced by 52–54% only in individuals with stage V–VI but not III-IV (Figure S1J and Table S3). Individuals with MMSE scores of ≤26 demonstrated 52-67% reductions in retinal pre- and post-synaptic markers as compared to those with MMSE scores below 26 (Figure S1K and Table S4). Spearman’s rank pairwise correlation analyses revealed significant associations between these retinal synaptic markers and brain AD-related pathology, Aβ-plaque or NFT burden (Figure 1K and Table S5). While weak to moderate associations were found between retinal synaptic markers and brain Aβ-plaque or NFT neuropathological burden, a strong association was detected between retinal synaptophysin and brain NFT severity (*r* = –0.65, P < 0.01; Table S5). Notably, retinal post-synaptic markers (PSD95, NMDAR2A) compared with the pre-synaptic proteins, showed more significant correlations with disease stage (Braak, ABC), whereases the pre-synaptic markers (VGLUT1, SYP) exhibited slightly stronger associations with the cognitive deficit (CDR, MMSE; Figure 1K). Overall, our findings reveal robust, early, and spatially distinct patterns of synaptic density losses in the human AD retina which were tightly coupled to both retinal and brain pathology. The peripheral retina emerges as a focal point of synaptic vulnerability, likely reflecting early Aβ and tau accumulation. These results support the concept that the retina undergoes early synaptic degeneration prior to neuronal cell loss (neurodegeneration), parallelling disease processes in the brain.

### Proteomic evidence of retinal neurosynaptic deficits in AD with p75NTR upregulation

We next sought to delineate synaptic- and neurodegeneration-targeted molecular pathways in the AD retina by analyzing an independent human cohort using an integrated protein biochemistry workflow encompassing Western blotting (WB), secondary mass spectrometry (MS), and quantitative enzyme-linked immunosorbent assay (ELISA). This cohort comprised neuropathologically confirmed AD patients (n = 9; mean age = 82.0 ± 15.7 years; 6 females, 3 males) and age- and sex-matched individuals with normal cognition (NC; n = 7; mean age = 76.7 ± 5.4 years; 4 females, 3 males). Initial proteomic findings from this cohort were previously reported (PRIDE ID: PXD040225)^28^. Preranked gene set enrichment analysis (GSEA)^85^ of AD retinal MS data further confirmed inhibited synapse- and vision-linked pathways, including KEGG ‘adherens junction’ and ‘phototransduction’, alongside GO ‘response to light stimulus’ and ‘actin filament–based transport,’ while activated GO ‘negative regulation of cell cycle’—consistent with disrupted retinal connectivity, impaired neuronal homeostasis, and neurodegeneration in AD (Figure S2).

Further, Ingenuity Pathway Analysis (IPA) of differentially expressed proteins (DEPs: |fold change| > 1.2 and FDR-adjusted P values < 0.2) in AD versus NC retinas revealed perturbation of retinal homeostasis and function, with prominent photoreceptor degeneration, alongside synaptic dysfunction spanning synaptic depression, dendritic spine density/architecture, synaptic vesicle trafficking, and neurotransmission (Figure 2A). A substantial fraction of retinal proteins expressed in photoreceptors, including several with exclusive photoreceptor expression, was markedly dysregulated in AD (Figure S3A–B). Within photoreceptor outer segments, the site of phototransduction, opsins including rhodopsin (RHO), long-wave–sensitive opsin 1 (OPN1LW), and short-wave–sensitive opsin 1 (OPN1SW) were reduced by ∼47%–61% in AD retinas, accompanied by significant decreases in cyclic nucleotide–gated channel subunits (CNGA1, CNGB1), transducin γ subunit (GNGT1), and Na^+^/Ca^2+^–potassium channels (SLC24A1/2), key components of the visual transduction cascade. Photoreceptor inner-segment proteins implicated in excitability and phototransduction—including the potassium channels KCNB1, KCNV2, and KCNMA1—were similarly decreased, together with peripherin-2 (PRPH2), in the AD retina (Figure S3A–B). In parallel, pro-apoptotic mediators were increased in the AD retina, including Fas associated protein with death domain (FADD), pro-apoptotic caspase 3 (CASP3) and the Aβ-binding death receptor p75-neurotrophin nerve growth factor receptor (p75NTR, also known as NGFR; Figure S3C–D)^76,86,87^. Retinal p75NTR levels tightly and inversely correlated with all photoreceptor markers (Figure S3E; *P_adj._* < 0.05), consistent with pronounced photoreceptor degeneration in AD.

**Figure 2.**
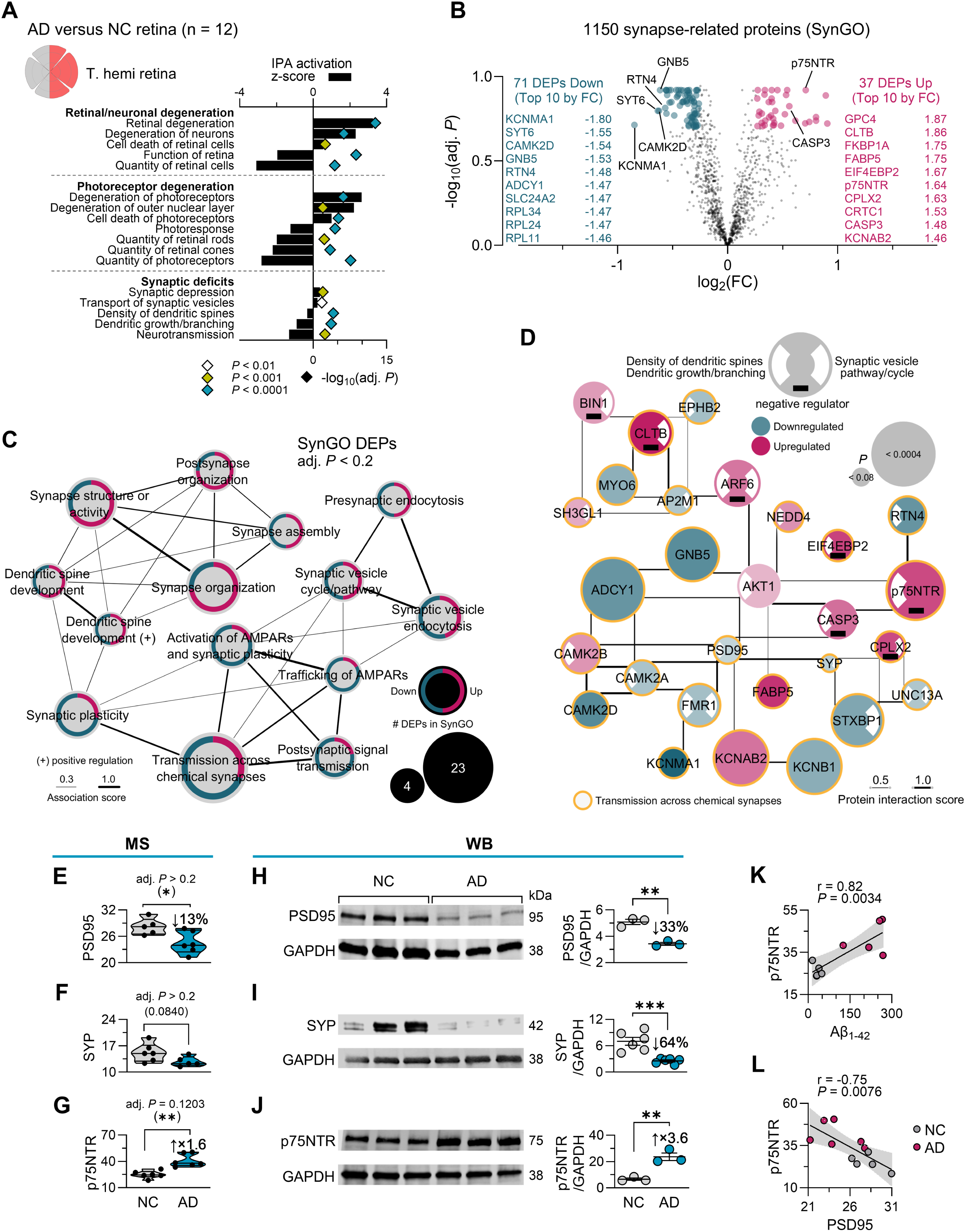
Dysregulated AD retinal proteome associated with synaptic and neuronal loss. (**A-F**) Pathway analysis of global proteomics via mass spectrometry (MS) in AD (n = 6) versus NC control (n = 6) retinas. For this analysis, DEPs were defined by |FC| > 1.2 and adj. *P* < 0.2. (**A**) Dysregulated biological pathways related to retinal/neuronal degeneration, photoreceptor degeneration and synaptic deficits, revealed by Ingenuity pathway analysis (IPA) of DEPs in the AD retina. Bar and symbol graphs represent activation z-scores, and Benjamini-Hochberg adjusted P-values, respectively. Range of P-values are indicated as color-coded symbols. (**B**) Volcano plot of dysregulated synapse-associated proteins in AD versus NC control retinas, extracted from SynGo release 1.2 knowledge base. The top 10 upregulated and downregulated DEPs are listed with their fold change (FC). (**C**) Pathway network (Metascape) from Gene Ontology (GO) analysis of synaptic-relevant DEPs revealed by SynGo. The size of the nodes represents the number of DEPs in SynGo, while the thickness of edges represents the overlapping DEPs (association score) between GO terms. (**D**) Interaction network analysis of DEPs related to synaptic plasticity, synaptic vesicle cycle and neurotransmission built from String v12.0 database. (**E-G**) Violin plots of the post-synaptic marker PSD95 (**E**), the pre-synaptic protein SYP (**F**) and the neuron-type-specific death receptor NGFR/p75NTR (**G**), quantified by MS, in AD versus control retinas. (**H-J**) Western blot analysis and normalized quantitation of retinal PSD95 (**H**), SYP (**I**) and p75NTR (**J**). (**K**) Pearson’s correlations between retinal Aβ_1-42_ levels, measured by ELISA, and retinal p75NTR, quantified by MS. (**L**) Pearson’s correlation between retinal PSD95 and retinal p75NTR. Violin plots show individual subjects (circles) with lower, median and upper quartiles. *P < 0.05, ** P < 0.01, *** P < 0.001, by unpaired Student t-test.

Alongside neurodegeneration, synaptic integrity was compromised, as evidenced by downregulation of synaptic proteins enriched in photoreceptor terminals, including synaptotagmin-6 (SYT6) and retinoschisin-1 (RS1) (Figure S3A). We analyzed synapse-associated DEPs in AD retinas using the SynGo 1.2 database (Figure 2B–D). Of the 1602 synapse-associated proteins included in the database, 1150 were detected in the human retina, with 37 significantly upregulated and 71 significantly downregulated in the AD retina (Figure 2B; list of synaptic-associated DEPs in Tables S6 and S7). These accounted for approximately 14% and 15% of upregulated and downregulated DEPs in the AD retina, respectively (Figure S3F). Gene Ontology (GO)-guided pathway analysis of the 108 significantly altered proteins, queried across GO Biological Process, Reactome, KEGG, and WikiPathways, converged on synaptic dysregulation in AD retinas, marked by reduced synaptic plasticity, chemical synaptic and postsynaptic signaling, and AMPAR activation/trafficking, alongside increased signatures of synaptic organization and structural remodeling (Figure 2C). Consistent with these changes, positive regulators of synaptic transmission were largely downregulated, including the Ca²_/calmodulin-regulated adenylyl cyclase 1 (ADCY1/AC1) linked to visual/sensory and photoreceptor biology, the core G-protein subunit β5 (GNB5) which tunes synaptic signaling and light responses, and the synaptic markers PSD95 and SYP (Figure 2D). Conversely, negative regulators of synaptic vesicle recycling—including synaptic vesicle endocytosis-related clathrin light chain B (CLTB), bridging integrator 1 (BIN1) and ADP ribosylation factor 6 (ARF6)—and dendritic spine formation—including the translation initiation repressor eukaryotic translation initiation factor 4E binding protein 2 (EIF4EBP2), CASP3 and p75NTR—were upregulated in the AD retina (Figure 2D).

Western blotting of retinal homogenates from a subset of AD patients and NC individuals corroborated the MS findings, confirming reduced expression of PSD95 (33% decrease; *P*<0.01) and synaptophysin (SYP; 64% decrease; P<0.001), alongside an increase in p75NTR (3.6-fold; *P*<0.01) in AD retinas (Figure 2E–J; full extended blots in Figure S4). Given that Aβ_42_ binds p75NTR and can promote neuritic degeneration, synaptic loss, and apoptosis, we next tested the p75NTR association with Aβ_1-42_ peptide levels (measured by a quantitative sandwich ELISA) and synaptic markers (Figure 2K–L). Retinal p75NTR expression very strongly correlated with retinal Aβ_1–42_ levels (Figure 2K; r=0.82, *P*=0.0034) and was inversely associated with the glutamatergic postsynaptic marker PSD95 (Figure 2L; r=−0.75, P=0.0076). Together, these relationships support the existence of an Aβ_42_–p75NTR axis contributing to synaptic loss and neurodegeneration in the AD retina.

Overall, these proteomic and biochemical analyses demonstrate that AD retinas undergo profound photoreceptor marker loss and synaptic dysfunction, potentially driven by Aβ_42_ accumulation and p75NTR-mediated apoptosis.

### Oxidative stress and p75NTR apoptotic signaling-mediated synaptic loss in the AD retina

Our protein biochemistry analyses show that synaptic protein deficits and activation of death receptor pathways have emerged as central pathological hallmarks of the AD retina. We next interrogated neurodegeneration-associated processes within anatomically defined retinal subregions and specific cellular layers in an expanded human cohort (Figure 3 and Table 3; extended data in Figure S5). To this end, we initially assessed neuronal integrity in retinas from MCI (n=9), AD (n=10), and age-matched NC controls (n=14) using neuronal-specific Nissl staining, which labels nuclei and granules (i.e., ribosomal RNA) in purple or blue (Figure 3A). Quantitative analysis revealed substantial neuronal loss across nuclear cell layers, with highly significant 32-33% losses of ONL neurons and 47–56% losses of INL neurons in MCI and AD retinas compared with NC controls (P < 0.01 – 0.001; Figure 3A). Retinal Nissl^+^ neuronal area was very strongly and directly correlated with retinal SYP (*r* = 0.82, P < 0.0001; Figure 3B) and strongly and negatively correlated with retinal Aβ_42_ burden (*r* = –0.65, P = 0.0001; Figure 3C), further supporting a link between Aβ_42_ accumulation and both neuronal and synaptic loss in the AD retina. There was also a moderate association between retinal Nissl and retinal CitR209-tau (*r* = 0.56, P = 0.05), but not with other tau isoforms (Table 3). To further characterize retinal neurodegeneration, we performed histomorphometric analysis of retinal thickness across six layers (ONL, OPL, INL, IPL, GC/NFL and NFL) in predefined C/M/F anatomical subregions (Figure 3D, mid periphery; extended data across all retinal layers and subregions are detailed in Figure S5A-I). The most noticeable thinning was detected in the mid-peripheral retina (Figure 3D), prominently in axonal- and synapse-rich layers (NFL, IPL, OPL), which were reduced by 50–63% in AD versus NC (P < 0.0001). The earliest and most pronounced loss in MCI retinas occurred in the GCL/NFL, diminished by 30–46% (P < 0.001 – 0.0001). These findings highlight region- and layer-specific vulnerability that mirrors synaptic degeneration and Aβ_42_ deposition, contributing to retinal neurodegeneration.

**Figure 3.**
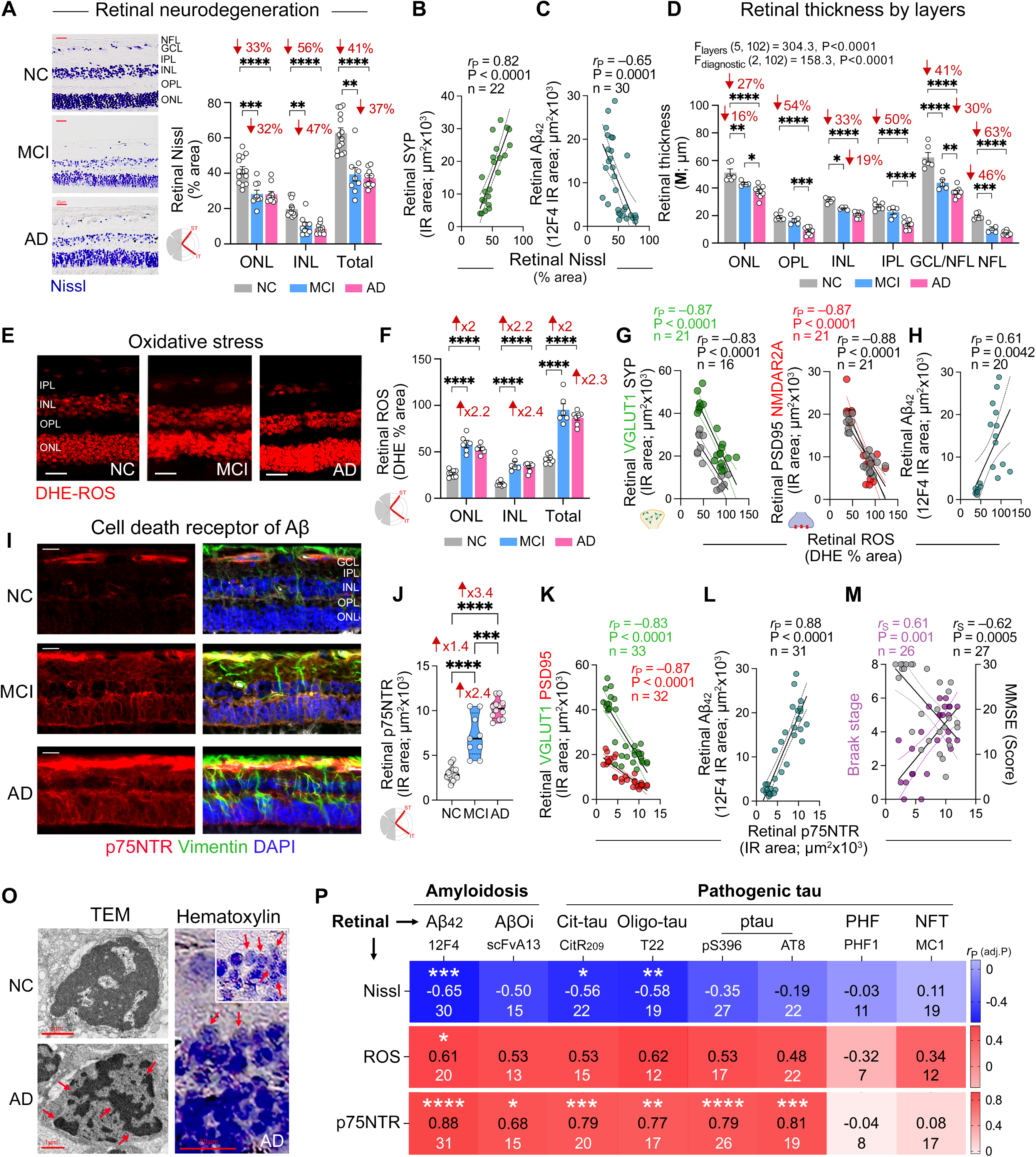
Retinal neurodegeneration and death receptor signaling in MCI and AD patient retinas. (**A**) Representative Nissl-stained retinal cross-sections from individuals with normal cognition (NC), MCI and AD dementia. Scale bars: 20 µm. Quantitative analysis of Nissl^+^ percent (%) area in the ONL, INL, and total retina (sum of GCL, INL, and ONL) of MCI (n = 9) and AD patients (n = 10) compared to NC controls (n = 14). (**B-C**) Pearson’s correlation coefficient (*r*_P_) analysis between Nissl^+^ % area and retinal (**C**) synaptophysin (SYP) and (**D**) Aβ_42_ burden. (**D**) Quantitative analysis of mid-peripheral retinal layer thickness in the superior- and inferior-temporal regions of the retina from MCI (n = 5), AD (n = 9) and NC (n = 6) individuals. (**E**) Representative images of dihydroethidium (DHE) staining showing reactive oxygen species (ROS) level in the retina from MCI, AD and NC individuals. Scale bars: 20 µm. (**F**) Quantitative analysis of retinal DHE-ROS levels in a subset of MCI (n = 6), AD patients (n = 7) and NC controls (n = 8). (**G-H**) Pearson’s correlation coefficient (*r*_P_) analysis between neuronal DHE^+^ % area and retinal (**G**) pre-synaptic markers VGLUT1 and SYP, post-synaptic markers PSD95 and NMDAR2A, and (**H**) Aβ_42_ burden. (**I**) Representative immunofluorescence images of retinal cross-sections from MCI, AD and NC individuals, labeled for p75NTR (nerve growth factor receptor; red) and vimentin (green); nuclei counterstained with DAPI (blue). Scale bars: 20 µm. (**J**) Violin plots display quantitative analysis of retinal p75NTR IR area in MCI (n=9) and AD (n=12) patients compared to matched NC controls (n = 13). (**K--L**) Pearson’s correlation coefficient (*r*_P_) analysis between retinal p75NTR versus (**K**) retinal VGLUT1 and PSD95, and (**L**) retinal Aβ_42_ burden, (**M**) Spearman’s rank correlation coefficient (*r*_S_) analysis between retinal p75NTR versus BRAAK stage and MMSE score. (**O**) TEM and H&E images showing neuronal nuclear morphology in NC and AD retinas. Shrunken, dense, and darkly stained pyknotic nuclei were observed in AD retinal cells (red arrows), indicative of cell death. (**P**) Heatmap showing Pearson’s correlations coefficient (*r*_P_) and significance (P values with Holm-Šídák multiple-comparison correction method adjusted for each retinal neuronal marker against 8 retinal synaptic, amyloidosis, and tauopathy marker) between retinal degeneration markers (Nissl, DHE-ROS, p75NTR) and retinal AD pathology, including amyloidosis (12F4^+^Aβ_42_ and scFvA13^+^ intraneuronal Aβ oligomers – AβOi) and tauopathy (pS396-tau, CitR_209_-tau, T22-tau oligomers, AT8-ptau, PHF-tau, MC1-NFTs). Violin plots show individual subjects (circles) with lower, median and upper quartiles. Bar graph displays group means ± SEMs. *P < 0.05, ** P < 0.01, *** P < 0.001, **** P < 0.0001, by one-way or two-way ANOVA followed by Tukey’s post-hoc multiple comparison test. Percentage (%) changes are shown in red. NFL, retinal nerve fiber layer; GCL, ganglion cell layer; IPL, inner plexiform layer; INL, inner nuclear layer; OPL, outer plexiform layer; ONL, outer nuclear layer.

**Table 3.**
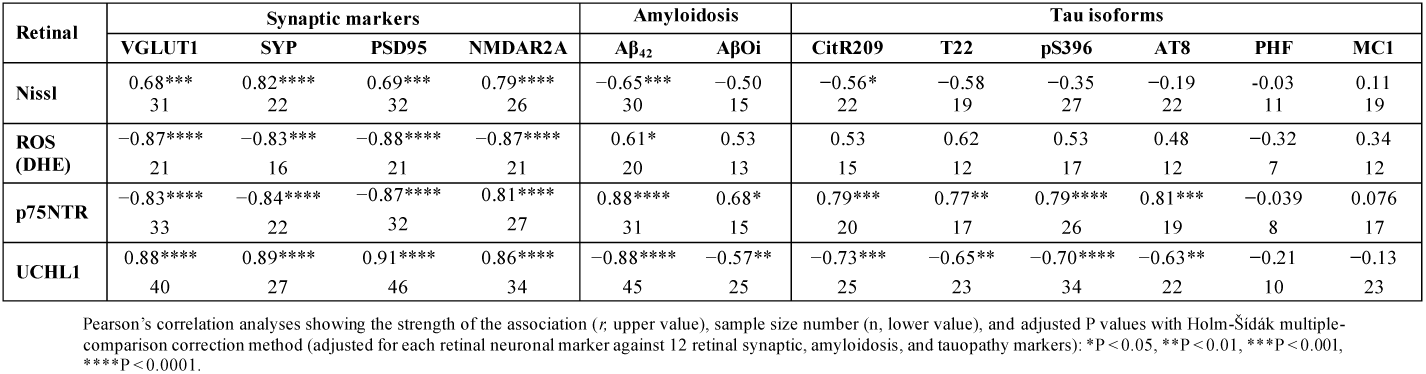
Correlations between retinal neurodegeneration-related and UCHL1 markers with retinal AD pathology.

Pathological Aβ and tau species have been shown to drive oxidative stress^88–90^. Given the retina’s high metabolic demand, vulnerability to oxidative damage, and progressive neurodegeneration in AD, we next quantified retinal labeling related to reactive oxygen species (ROS) in the MCI and AD retina (Figure 3E-F). Dihydroethidium (DHE) labeling^91^ revealed a 2.0–2.4-fold increase in oxidized DHE fluorescence (percent area), consistent with elevated superoxide/ROS levels, in the retinal nuclear layers of MCI and AD patients compared with NC controls (P < 0.0001). Retinal DHE levels exhibited very strong inverse correlations with retinal synaptic markers (Figure 3G, Table 3), most notably PSD95 (*r* = –0.88, P*_adj._* < 0.0001) and NMDAR2A (*r* = –0.87, P*_adj._* < 0.0001), alongside strong positive correlations with 12F4^+^ Aβ_42_ area (*r* = 0.61, P = 0.0042; Figure 3H) but not with pathogenic tau species. These findings implicate oxidative stress as a downstream mediator of Aβ_42_ accumulation driving synaptic and neuronal loss in the AD retina.

Previously, p75NTR has been implicated in regulating oxidative stress, apoptosis, neuronal survival, and synaptic plasticity, and in mediating Aβ-driven neurite degeneration, tau hyperphosphorylation, and neuronal loss in AD models^72–75^, and has emerged as a top upregulated synapse-associated protein in AD retina in our MS and WB analyses (Figure 2D, 2I, 2L). Consistent with these findings, IHC analysis revealed robust increases in p75NTR expression in MCI and AD retinas (2.4–3.4-fold vs NC; *P* < 0.0001), with signal localizing predominantly to vimentin^+^ Müller glia (Figure 3I-J). Moreover, retinal p75NTR expression exhibited very strong inverse correlations with all retinal synaptic markers (Table 3), including the pre- and post-synaptic markers VGLUT1 and PSD95 (*r* = –0.83 and –0.87, respectively, P < 0.0001; Figure 3K), and a very tight direct correlation with retinal Aβ_42_ burden (*r* = 0.88, P < 0.0001, Figure 3L), supporting its role as a mediator of Aβ-induced synaptic loss in the retina, mirroring mechanisms observed in the AD brain^74,92–95^. Interestingly, strong associations between retinal p75NTR and immature but not mature tau isoforms were also observed (*r* = 0.77–0.81, P*_adj._* = 0.0012–0.00001, Table 3), proposing the involvement of tauopathy in p75NTR-mediated synaptic degeneration in the AD retina. We next assessed the clinical relevance of retinal p75NTR by evaluating its potential associations with brain AD pathology and cognitive status (Table 4). Retinal p75NTR expression moderate-to-strongly correlated with brain atrophy, Braak stage, and ABC scores (*r* = 0.53–0.61, P*_adj._* = 0.024–0.0061; Figure 3M and Table 4) and inversely associated with MMSE scores (*r* = –0.62, P = 0.0005; Figure 3M), indicating that retinal p75NTR upregulation reflects both disease severity and cognitive impairment.

**Table 4.**
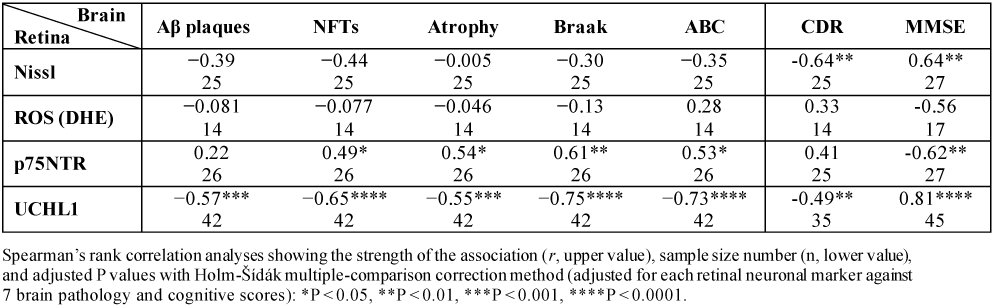
Correlations between retinal UCHL1 and neurodegeneration-related markers with brain pathology and cognition.

TEM and hematoxylin and eosin (H&E) staining further showed the ultrastructural features of neuronal death, revealing shrunken, condensed, and darkly stained pyknotic nuclei, hallmarks of oxidative stress-induced apoptosis or necroptosis, in AD compared with NC retinas (Figure 3O, red arrows). A heatmap summary analysis (Figure 3P) highlights strong inter-relationships among retinal Aβ/tau pathologies and neurodegeneration, with retinal p75NTR emerging as a robustly correlated marker. These correlations reinforce p75NTR as a central integrator of Aβ- and tau-driven toxicity and a potential mediator of synaptic degeneration in the retina, paralleling AD mechanisms in the brain^74,92–95^.

### Retinal UCHL1 dysregulation in AD is a strong predictor of Braak stage and cognitive status

Ubiquitin carboxyl-terminal hydrolase L1 (UCHL1), a neuron-enriched deubiquitinating enzyme and key component of the UPS, is essential for proteostasis, oxidative stress responses, synaptic integrity, and neuronal survival (Figure 4A)^60,61,96^. In the brain, UCHL1 facilitates clearance of misfolded proteins and limits the aggregation of neurotoxic species such as Aβ_42_ and p-Tau^97^, whereas reduced UCHL1 expression has been linked to Aβ and pTau–mediated synaptic dysfunction and cognitive decline in AD^56,62^. We therefore evaluated UCHL1 expression and its potential association with synaptic loss and neurodegeneration in postmortem retinas from a cohort of AD (n=21, mean age 86.2 ± 8.9 years, 13 females/8 males) and MCI (n=11, mean age 89.3 ± 5.2 years, 7 females/4 males) patients, compared with NC controls (n=18, mean age 82.6 ± 10.3 years, 8 females/10 males) (Figure 4B-D; extended data in Fig. S6A-D).

**Figure 4.**
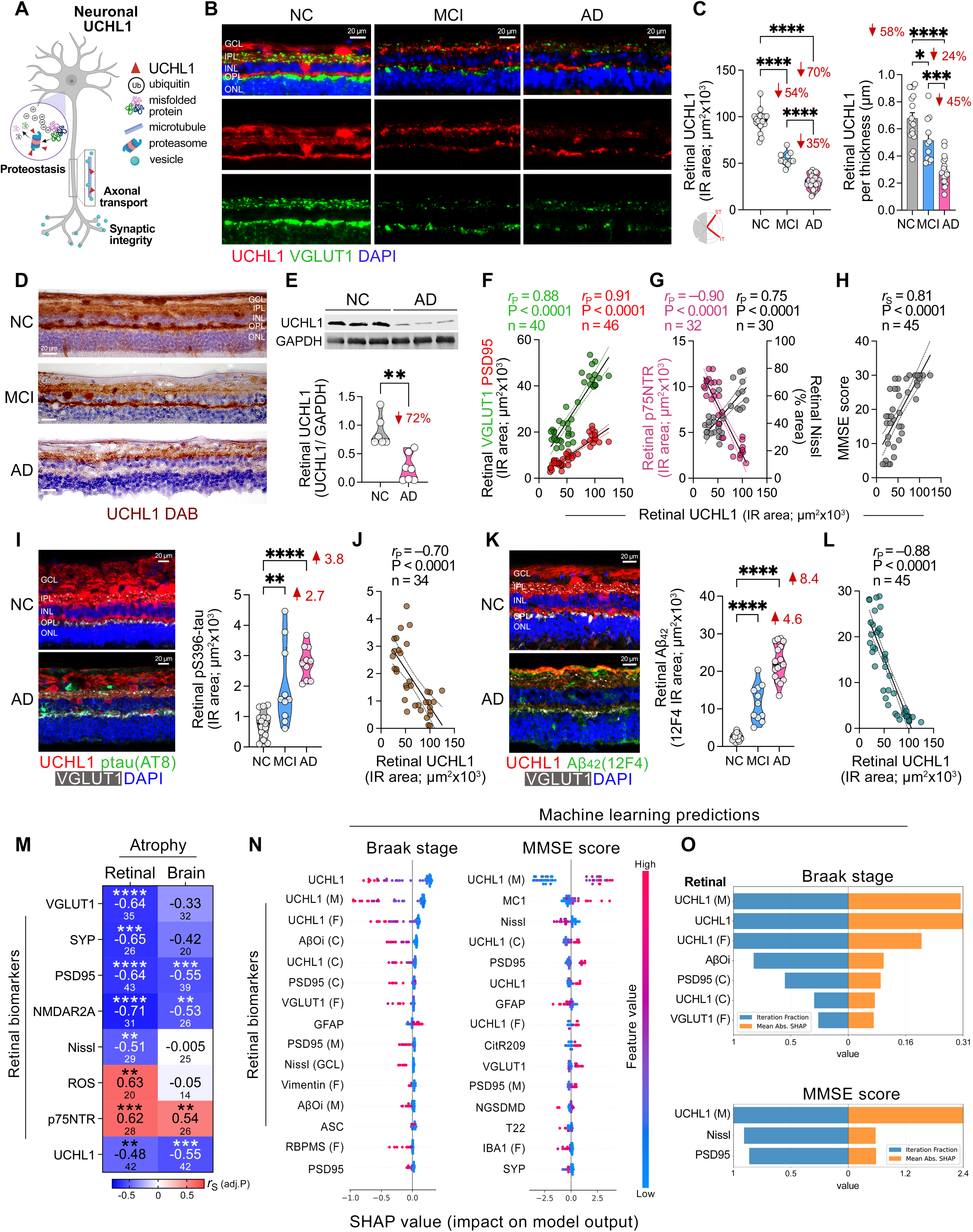
Retinal UCHL1 in relation to synapses and proteinopathies. (**A**) Schematic illustration of UCHL1 functions in neurons. (**B**) Representative fluorescent microscopic images of retinal cross sections immunolabelled for UCHL1 (red) in combination with pre-synaptic marker – VGLUT1 (green). Nuclei are labelled with DAPI (blue). Scale bars: 20 µm. (**C**) Violin plots display quantitative analysis of retinal UCHL1 IR area in patients with AD (n = 21) and MCI (n = 11) as compared with NC controls (n = 18). (**D**) Representative retinal cross-section images from MCI and AD patients compared to NC controls immunolabeled with UCHL1 (brown) using peroxidase-based DAB and hematoxylin counterstaining. Scale bars: 20 µm. (**E**) Representative Western blot showing the levels of UCHL1 in retinal lysates from AD patients versus NC (upper panel). The densitometric analyses of UCHL1, normalized to GAPDH, is shown (lower panel) (n = 7 AD, 6 NC). (**F-H**) Pearson’s or Spearman’s correlation analyses between retinal UCHL1 IR area and the following: (**F**) retinal pre-synaptic marker - VGLUT1 (green) or post-synaptic marker - PSD95 (red), (**G**) retinal p75NTR and Nissl, and (**H**) the MMSE-cognitive scores. (**I**) Representative fluorescent images of retinal cross-sections, immunostained for UCHL1 (red), AT8^+^-ptau (green), and VGLUT1 (white). Nuclei are stained with DAPI (blue). Scale bars: 20 µm. The violin plot presents quantitative analysis of retinal pS396 IR area across subjects in this cohort with AD (n = 10), MCI (n= 10) and NC (n =15). (**J**) Pearson’s correlation analysis between retinal UCHL1 and retinal pS396-tau IR areas. (**K**) Representative fluorescent images of retinal cross-sections immunostained for UCHL1 (red), 12F4^+^ Aβ_42_ (green), and VGLUT1 (white). Nuclei are stained with DAPI (blue). Scale bars: 20 µm. The violin plot presents quantitative analysis of retinal 12F4^+^Aβ_42_ IR area across subjects in this cohort with AD (n = 17), MCI (n= 11) and NC (n =19). (**L**) Pearson’s correlation analysis between retinal UCHL1 IR area and retinal Aβ_42_ burden. (**M**) Heatmap showing Spearman’s rank correlation coefficient (*r_S_*) and significance (adjusted P values with Holm-Šídák multiple-comparison correction method for each retinal neuronal marker against multiple retinal and brain atrophy markers) between retinal (left) and brain (right) atrophy to retinal synaptic, neurodegeneration and UCHL1 markers. (**N-O**) Identification of features impacting model predictions for (N) Braak staging and (O) MMSE score. The left bars indicate the fraction of randomly-instantiated models where feature was in the top 5 features ranked by mean absolute SHAP value; the right bars indicate the mean absolute SHAP value. Violin plots show individual subjects (circles) with lower, median and upper quartiles. *P < 0.05, ** P < 0.01, *** P < 0.001, **** P < 0.0001, by unpaired Student t-test or one-way ANOVA followed by Tukey’s post-hoc multiple comparison test. Percentage (%) changes are shown in red. GCL, ganglion cell layer; IPL, inner plexiform layer; INL, inner nuclear layer; OPL, outer plexiform layer; ONL, outer nuclear layer. Schematic illustration was created using Biorender.com.

In NC retinas, double immunofluorescence labeling of UCHL1 with the presynaptic marker VGLUT1 and peroxidase-based immunolabeling of UCHL1 with hematoxylin counterstaining showed intense UCHL1 expression across multiple retinal cell types, including horizontal, bipolar, amacrine, and ganglion cells. Robust expression was also evident in synapse-rich and axonal layers (OPL, IPL, NFL), whereas only sparse signal was detected in the nuclear ONL and INL (Figure 4B, 4D). Within the OPL and IPL, UCHL1 was found in close spatial proximity to VGLUT1 but exhibited limited direct co-localization, suggesting a complementary rather than overlapping synaptic distribution. In contrast, MCI and AD retinas exhibited significant decrease in UCHL1 expression in these cell types and layers, paralleling a concomitant decrease in VGLUT1 expression, indicating a tight association between UCHL1 loss and synaptic degeneration (Figure 4B). Quantitative analysis revealed significant 54–70% reductions in UCHL1 immunoreactive area in retinas from MCI and AD patients compared to NC controls (Figure 4C, left graph, P < 0.0001; Extended UCHL1 analyses of percent immunoreactive area and regional mapping, confirming 47–70% reductions across the ST/IT retina and C/M/F subregions in MCI and AD in Figure S6B-C). To determine whether reduced retinal UCHL1 simply reflects retinal thinning or neurodegeneration (significant in MCI and AD), we normalized UCHL1 to retinal thickness. The reduction persisted: UCHL1 remained significantly decreased by 24% in MCI and 45% in AD retinas (Fig. 4C, right graph, P < 0.05-0.0001). In agreement with these results, Western blot analysis in additional human cohort demonstrated a 72% decrease in UCHL1 expression normalized to GAPDH in 7 AD retinas relative to 6 NC controls (Figure 4E; a representative full WBs in Figure S6E). Proteomics showed broad UPS dysregulation in AD retinas: GSEA indicated suppression of the GO ‘protein SUMOylation’ program alongside activation of the WikiPathways ‘Parkin–ubiquitin–proteasomal system pathway’, consistent with stress-driven proteostasis and UCHL1-linked post-translational modification (Figure S7A-B). In line with prior work implicating UPS dysfunction in AD^64^, MS further revealed increased particulate/aggregated and oxidized UCHL1 and UCHL3 (Figure S7C-D), whereas our IHC and WB analyses showed reduced soluble UCHL1, together indicating a marked retinal UCHL1 imbalance in AD, paralleling reports in the AD brain^98–102^.

To assess the potential clinical significance of retinal UCHL1 in AD, we assessed its relationships with various retinal markers, brain AD-pathology, and cognitive parameters using correlation analyses (Figure 4F-H and Tables 3-4). Retinal UCHL1 exhibited robust correlations with all synaptic markers, particularly the pre-synaptic VGLUT1 and the post-synaptic PSD95 expression (*r* = 0.88-0.91, P*_adj._* < 0.0001; Figure 4F and Table 3). We also identified very strong to strong inverse associations between retinal UCHL1 and the cell death receptor p75NTR (*r* = –0.90, P < 0.0001), retinal neurodegeneration Nissl^+^ % area (*r* = 0.75, P < 0.0001; Figure 4G), and retinal DHE/ROS levels (*r* = –0.77, P < 0.0001; not shown). Moreover, retinal UCHL1 displayed a strong positive correlation with MMSE scores (*r* = 0.81, P*_adj._*< 0.0001; Figure 4H and Table 4), suggesting a direct link between retinal UCHL1 loss and cognitive decline. Notably, marked reductions in UCHL1 in MCI and AD retinas coincided with elevated pathogenic tau and Aβ species (Figure 4I–L, Table 3). With the exception of retinal mature PHF-tau and MC1^+^ tau, UCHL1 showed moderate to very strong inverse correlations with various abnormal tau and Aβ species, most prominently pS396-tau (*r* = –0.70, P*_adj._* < 0.0001), CitR209-tau (*r* = –0.73, P*_adj._* < 0.001), and Aβ_42_ burden (*r* = –0.88, P*_adj._* < 0.0001; Figure 4J, 4L, Table 3).

Multivariable correlations between retinal UCHL1 expression (IHC) and brain AD pathology, disease stage, and cognitive score are summarized in Table 4. Notably, UCHL1 expression inversely correlated with retinal and brain atrophy (Figure 4M), mirroring the pattern observed for synaptic markers and Nissl^+^ neuronal area. Conversely, retinal DAE-ROS levels and p75NTR expression were positively associated with retinal and brain atrophy, suggesting that oxidative stress and death receptor signaling contribute to widespread neurodegeneration observed in both the retina and brain in AD (Figure 4M). Heatmap analysis and Metascape network mapping demonstrated coordinated dysregulation of deubiquitination/UPS proteins and pathways in AD retinas (Figures S7E–F), with interconnected disruption of UPS, autophagy, axonal transport, and axonogenesis, positioning retinal UCHL1 as a central hub linking proteostasis failure to p75NTR-mediated neurodegeneration.

Finally, we trained Random Forests^103^ to predict Braak stage and MMSE score. Learning curves (Figure S7G-H) indicate that the model predicting Braak stage likely had sufficient data, but that the one for MMSE retained significant differences across folds. Both Random Forest models were trained with 80 estimators and L1 loss and used an identical retinal biomarker feature set. The mean R^2^ score across 5-fold cross-validation was 0.370 for predicting Braak stage, and 0.137 for predicting MMSE. Among all dysregulated retinal proteins, UCHL1 and PSD95 emerged as the top predictors of both Braak stage and MMSE score (Figure 4N), retinal AβOi (center) was predictive of Braak stage but not MMSE, and retinal Nissl signal and MC1 (tau tangles) were predictive of MMSE but not Braak. To mitigate sensitivity to initialization, we refit each model across 1,000 random seeds and quantified feature robustness by the consistency of their SHAP-derived importance across runs (Figure 4O). The impact of each feature on model prediction varied considerably, but UCHL1 was both consistently identified and had the greatest impact. These results highlight the potential of using non-invasive retinal synaptic biomarkers for monitoring AD progression and cognitive decline.

## Discussion

The present study reveals that retinal Aβ/p75NTR-mediated synaptic degeneration is coupled with profound dysregulation of UCHL1, which emerges as an early and defining pathological feature of AD. Across independent postmortem cohorts, including donors with MCI due to AD, we identified marked reductions in excitatory presynaptic and postsynaptic proteins, coupled with aberrant p75NTR signaling and UCHL1 imbalance, accompanied by ultrastructural collapse of synaptic architecture. These alterations were tightly associated with local Aβ_42_ and pathogenic tau species, previously reported to be elevated in AD patients^21,27–30,35–37,40,41^ and extended to measures of disease stage, brain pathology, and cognitive impairment, with retinal UCHL1 levels showing particularly strong associations. Collectively, our findings suggest that the retina undergoes a Aβ/p75NTR-mediated synaptic decline that parallels—and may faithfully reflect—the earliest cerebral synaptic pathology in AD.

A notable finding was the non-uniform spatial distribution of synaptic degeneration, with mid- and far-peripheral regions exhibiting the most pronounced deficits in MCI and AD patients compared with NC controls. The observed 50-77% reduction in pre-synaptic markers and 48-72% in post-synaptic markers highlights significant synaptic compromise even at the earliest symptomatic stages of the AD progression (MCI). This pattern of synaptic degeneration was consistent with current and prior evidence of peripheral-predominant Aβ_42_ and tau deposition in the retina^22,27,28,30,37,104^ and suggests that localized microenvironments may confer heightened vulnerability to proteinopathies. Consistent with our previous study, primary cortical neurons exhibited a 50–70% loss of glutamatergic pre- and postsynaptic densities following exposure to fibrillar or oligomeric Aβ_42_ aggregates^18^. Beyond Aβ, here we found that retinal immature and oligomeric tau isoforms, rather than mature PHF- or MC1-positive tau aggregates, were tightly correlated with synaptic loss, echoing accumulating evidence from the brain that soluble, non-fibrillar tau species exert disproportionate synaptotoxic effects^105–107^.

Retinal proteome analysis highlighted a dual vulnerability in the AD retina: retinal synaptic failure and photoreceptor degeneration. Photoreceptor degeneration was determined by broad downregulation of photoreceptor proteins, including opsins and other critical elements of the phototransduction cascade. Analysis of synapse-associated proteins indicated dysregulation of critical molecular pathways related to the synaptic vesicle cycle, neuronal synaptic plasticity, and dendritic spine development. Decreased levels of positive regulators of synaptic docking—including SYP, UNC13A, and STXBP1—and increase of negative regulators of synaptic endocytosis—including BIN1, ARF6, and CLTB—indicate the disruption of neurotransmitter release and synaptic vesicle recycling, respectively^108–114^. Other key positive regulators of synaptic transmission—such as ADCY1, EPHB2, GNB5 and PSD95^111,115,116^—were largely downregulated. Together with GSEA suggesting dysregulated synaptic related pathways, these alterations suggest impaired vesicle cycling and reduced excitatory transmission, converging on mechanisms of functional disconnection previously established in hippocampal and cortical circuits^117–121^.

The robust upregulation of p75NTR emerged as a particularly striking feature of the AD retina. A death receptor, p75NTR plays key roles in axon pruning, synaptic plasticity, and apoptosis in the CNS ^73,122^. In AD models, Aβ binding to p75NTR induces neuritic degeneration, tau hyperphosphorylation, oxidative stress, and neuronal loss^72,74,75^. Elevated p75NTR levels correlated closely with reduced synaptic proteins (VGlut1, PSD95), increased Aβ_42_ burden, immature tau isoforms, and elevated oxidative stress, suggesting its role as a mediator of convergent neurotoxic pathways. Given the ability of Aβ to bind p75NTR and trigger apoptotic signaling^76,86,87^, our data support the possibility that p75NTR integrates multiple pathogenic inputs, Aβ, tau, and oxidative stress, culminating in synaptic degeneration. These findings align with prior work in AD models demonstrating that p75NTR contributes to neuritic degeneration, tau hyperphosphorylation, and neuronal loss in the AD retina^74,75,123^.

A central and novel observation is the dysregulation of UCHL1, a neuron-enriched deubiquitinating enzyme critical for proteostasis and synaptic integrity^61,124^. UCHL1 expression was markedly reduced in MCI and AD retinas, particularly in synapse-rich layers, while mass spectrometry revealed accumulation of particulate and oxidized forms. UCHL1 is susceptible to ROS and oxidized UCHL1, which is prone to aggregation, accumulates in AD brains^98,99^, whereas soluble UCHL1 levels and activity are reduced^101^. The antibody used in this study primarily recognizes the cytosolic, soluble form of UCHL1, which has been reported to be reduced in brain lysates from patients with AD and Parkinson’s disease^60,125^. This reduction may explain the diminished UCHL1 signal observed in the AD retina through fluorescence and non-fluorescence histological analyses and further validation by biochemical analysis should demonstrate the soluble form of UCHL1. In contrast, mass spectrometry likely detects the oxidized form. This imbalance echoes brain studies linking UCHL1 dysfunction to impaired clearance of Aβ and tau aggregates^126,127^. In our cohorts, retinal UCHL1 strongly correlated with synaptic markers, p75NTR, ROS levels, pathogenic tau isoforms, Aβ_42_ burden, and cognitive performance. Network analyses further implicated retinal UCHL1 in interlinked processes of autophagy, axonal transport, and synaptic recycling, suggesting that loss of soluble UCHL1 may represent a nodal point at which multiple pathogenic cascades converge. Indeed, our data suggests that UCHL1 sits at the crossroads of different disease mechanisms (proteostasis, oxidative stress, tau and Aβ toxicity, synaptic dysfunction) and its dysregulation could connect these otherwise separate processes into a common retinal and brain neurodegenerative pathway.

Correlative analyses of both retinal and brain pathologies revealed that both synaptic markers and UCHL1 track with Braak stages, brain atrophy, and cognitive impairment. Importantly, AI-based multivariable modeling identified retinal UCHL1 as the top predictor of disease stage and MMSE score, underscoring its translational potential as a non-invasive biomarker. While these findings cannot establish causality, they highlight the retina as a uniquely accessible site in which to interrogate synaptic and proteostatic dysfunction in AD.

Several limitations warrant consideration. Postmortem analyses cannot capture longitudinal trajectories or establish causal mechanisms, and differences in tissue preservation may affect protein integrity. Cohort sizes, while substantial for retinal neuropathological studies, remain modest for multivariable modeling. Finally, although retinal findings closely parallel those observed in the brain, the extent to which retinal pathology reflects systemic versus locally driven processes remains unresolved. Future work integrating in vivo imaging of retinal UCHL1 or synaptic integrity with functional and mechanistically targeted studies will be essential to validate these findings and assess their diagnostic utility.

In summary, our results position the retina as a critical site of early synaptic degeneration, proteostatic imbalance, and death receptor signaling in AD. The convergence of Aβ_42_, tau, oxidative stress, p75NTR, and UCHL1 dysregulation highlights interconnected mechanisms that likely contribute to both retinal and cerebral neurodegeneration. By revealing retinal UCHL1 as a strong predictor of disease stage and cognitive impairment, these findings provide a foundation for the development of novel retinal biomarkers. Such advances may deepen our understanding of AD pathophysiology and provide new means for clinically monitoring the progression of the disease against various conditions including purported new treatments.

## Lead contact

Further information and requests for resources and reagents should be directed to and will be fulfilled by the lead contact, Maya Koronyo-Hamaoui

## Material availability

This study did not generate new unique reagents.

## Data and code availability

Mass Spectrometry data are publicly available via the PRIDE database under the ProteomeXchange accession #PXD040225. All data are available in the main text, tables, and figures or supplementary materials (Figure S1-S7 and Tables S1-S7). All other material or data requests should be addressed to the corresponding author. The Jupyter notebook containing the analysis with random forests has been uploaded to https://github.com/xomicsdatascience/ubiquitin_ad. Data will become publicly available as of the date of publication.

## Acknowledgments

This work was supported by the National Institutes of Health (NIH)/the National Institute on Aging (NIA) through the following grants: R01R01AG056478, R01AG055865, and AG056478-04S1 (MKH), the National Institute for General Medical Sciences: R35GM142502 (JGM), The Hertz Innovation Fund and the Gordon, Goldrich, Wilstein, and Saban Private Foundations (MKH). M.R.D. and E.R. were supported by The Ray Charles Foundation. We thank Elijiah Maxfield for editing the manuscript. We thank Drs. Giovanni Meli and Antonino Cattaneo for providing the Aβ oligomers (scFvA13), Rakez Kayed, Maj-Linda B. Selenica, Daniel C. Lee and late Peter Davies for sharing the T22, CitR209, and PHF-1 antibodies used in our previously published work. We thank Dr. Carol A. Miller for providing a portion of the human tissues and neuropathological reports and Cedars-Sinai Biobank and Pathology Core for assistance with retinal tissue FFPE processing. Illustrations Figure 1A and Figure 4A were created with BioRender. This article is dedicated to the memory of Dr. Salomon Moni Hamaoui and Lillian Jones Black, both of whom passed away from Alzheimer’s disease.

## Author Contributions

**Conceptualization**, A.R., M.K.H.; **Methodology**, A.R., J-P.V., A.H., Y.K., S.S., E.R., E.B., An.R., B.P.G., O.J., J.S., D.H., L.S.S, J.G.M, D-T.F., M.K.H.; **Formal analysis and data curation**, A.R., J-P.V., A.H., Y.K., S.S., E.R., E.B., An.R., O.J., J.S., D.H, L.S.S, M.M., J.G.M, D-T.F., M.K.H.; **writing – original draft**, A.R., J-P.V., B.P.G., D-T.F., M.K.H.; **writing – review & editing**: all authors; **visualization**, A.R., J-P.V., A.H., Y.K., S.S., J.G.M., D-T. F., M.K.H.; **supervision**, M.K.H.; **funding acquisition**: M.K.H., K.L.B

## STAR★METHODS

### Key resources table

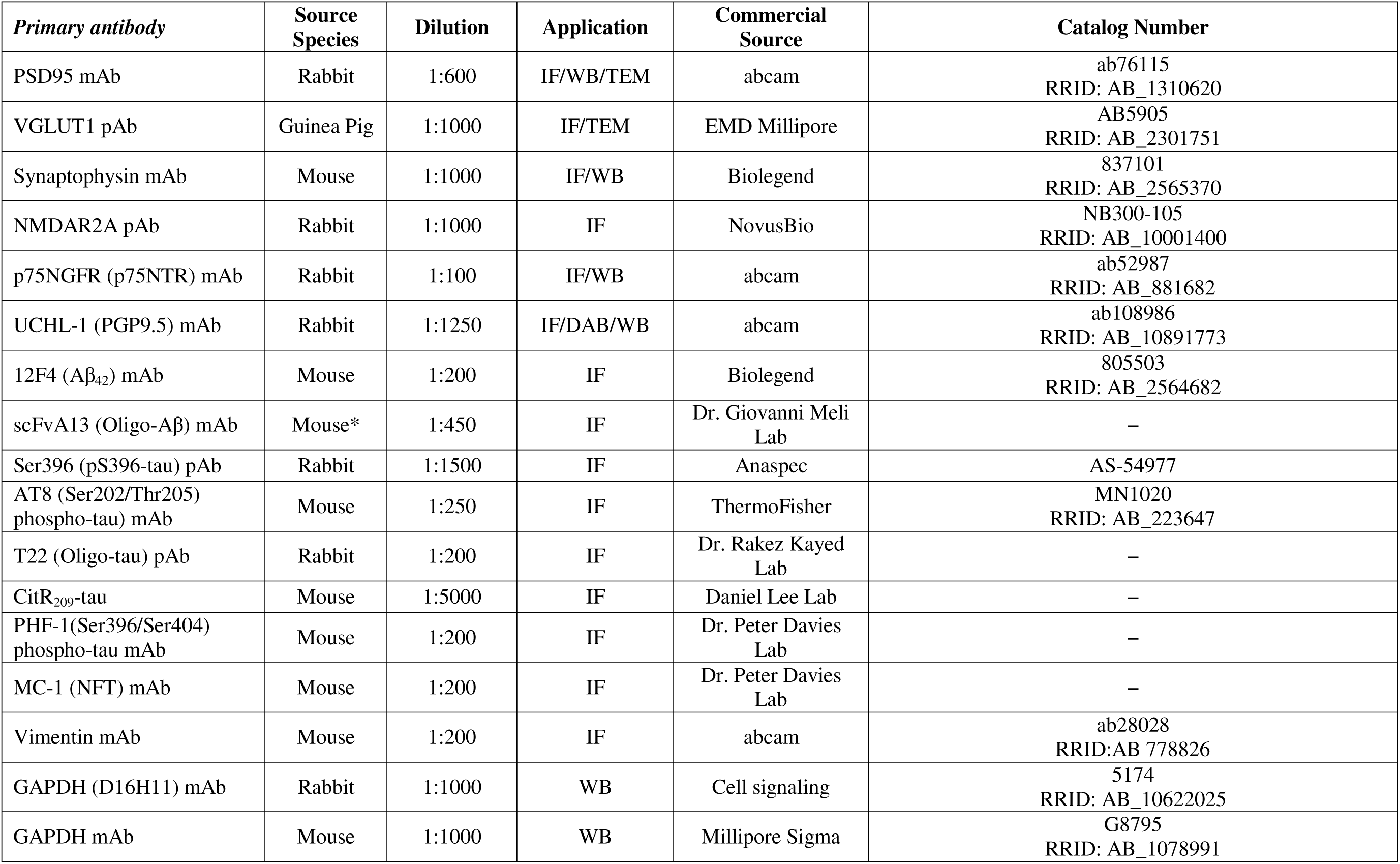
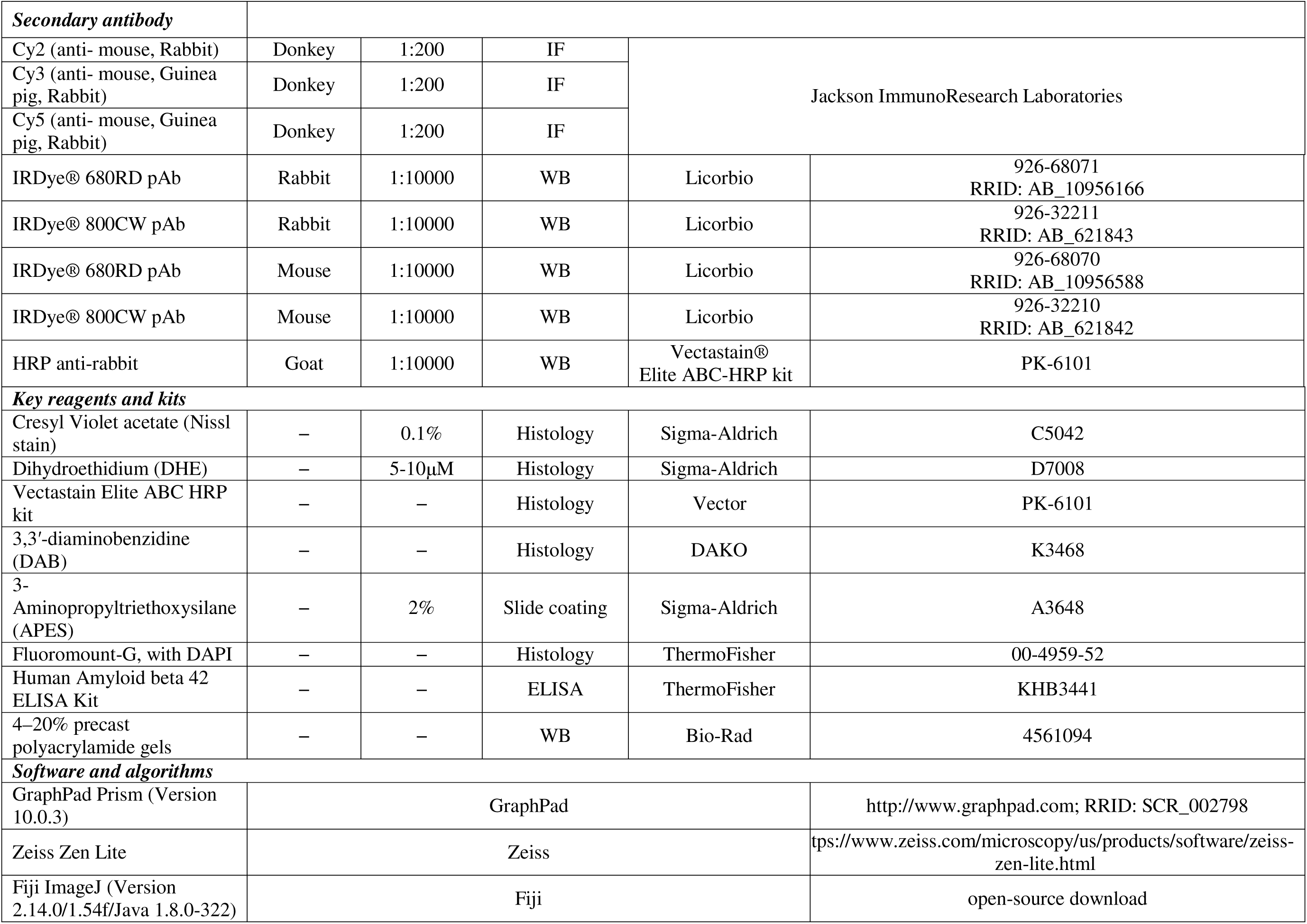
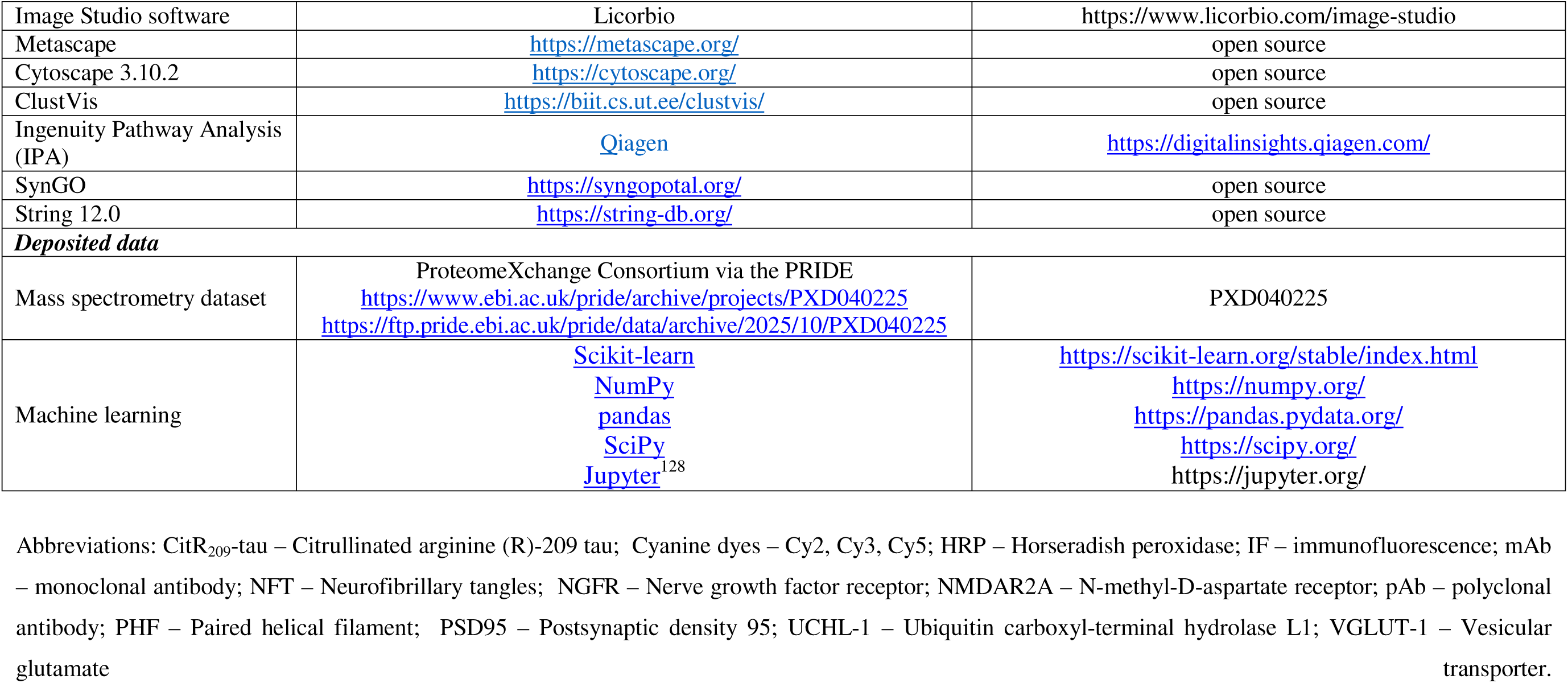

| *Primary antibody* | **Source Species** | **Dilution** | **Application** | **Commercial Source** | **Catalog Number** |
| --- | --- | --- | --- | --- | --- |
| PSD95 mAb | Rabbit | 1:600 | IF/WB/TEM | abcam | ab76115<br>RRID: AB_1310620 |
| VGLUT1 pAb | Guinea Pig | 1:1000 | IF/TEM | EMD Millipore | AB5905<br>RRID: AB_2301751 |
| Synaptophysin mAb | Mouse | 1:1000 | IF/WB | Biologend | 837101<br>RRID: AB_2565370 |
| NMDAR2A pAb | Rabbit | 1:1000 | IF | NovusBio | NB300-105<br>RRID: AB_10001400 |
| p75NGFR (p75NTR) mAb | Rabbit | 1:100 | IF/WB | abcam | ab52987<br>RRID: AB_881682 |
| UCHL-1 (PGP9.5) mAb | Rabbit | 1:1250 | IF/DAB/WB | abcam | ab108986<br>RRID: AB_10891773 |
| 12F4 (Aβ<sub>42</sub>) mAb | Mouse | 1:200 | IF | Biologend | 805503<br>RRID: AB_2564682 |
| scFvA13 (Oligo-Aβ) mAb | Mouse* | 1:450 | IF | Dr. Giovanni Meli Lab | – |
| Ser396 (pS396-tau) pAb | Rabbit | 1:1500 | IF | Anaspec | AS-54977 |
| AT8 (Ser202/Thr205) phospho-tau) mAb | Mouse | 1:250 | IF | ThermoFisher | MN1020<br>RRID: AB_223647 |
| T22 (Oligo-tau) pAb | Rabbit | 1:200 | IF | Dr. Rakez Kayed Lab | – |
| CitR<sub>209</sub>-tau | Mouse | 1:5000 | IF | Daniel Lee Lab | – |
| PHF-1(Ser396/Ser404) phospho-tau mAb | Mouse | 1:200 | IF | Dr. Peter Davies Lab | – |
| MC-1 (NFT) mAb | Mouse | 1:200 | IF | Dr. Peter Davies Lab | – |
| Vimentin mAb | Mouse | 1:200 | IF | abcam | ab28028<br>RRID:AB_778826 |
| GAPDH (D16H11) mAb | Rabbit | 1:1000 | WB | Cell signaling | 5174<br>RRID: AB_10622025 |
| GAPDH mAb | Mouse | 1:1000 | WB | Millipore Sigma | G8795<br>RRID: AB_1078991 |
| **Secondary antibody** |  |  |  |  |  |
| Cy2 (anti- mouse, Rabbit) | Donkey | 1:200 | IF | Jackson ImmunoResearch Laboratories |  |
| Cy3 (anti- mouse, Guinea pig, Rabbit) | Donkey | 1:200 | IF |  |  |
| Cy5 (anti- mouse, Guinea pig, Rabbit) | Donkey | 1:200 | IF |  |  |
| IRDye® 680RD pAb | Rabbit | 1:10000 | WB | Licorbio | 926-68071<br>RRID: AB_10956166 |
| IRDye® 800CW pAb | Rabbit | 1:10000 | WB | Licorbio | 926-32211<br>RRID: AB_621843 |
| IRDye® 680RD pAb | Mouse | 1:10000 | WB | Licorbio | 926-68070<br>RRID: AB_10956588 |
| IRDye® 800CW pAb | Mouse | 1:10000 | WB | Licorbio | 926-32210<br>RRID: AB_621842 |
| HRP anti-rabbit | Goat | 1:10000 | WB | Vectastain®<br>Elite ABC-HRP kit | PK-6101 |
| **Key reagents and kits** |  |  |  |  |  |
| Cresyl Violet acetate (Nissl stain) | – | 0.1% | Histology | Sigma-Aldrich | C5042 |
| Dihydroethidium (DHE) | – | 5-10µM | Histology | Sigma-Aldrich | D7008 |
| Vectastain Elite ABC HRP kit | – | – | Histology | Vector | PK-6101 |
| 3,3'-diaminobenzidine (DAB) | – | – | Histology | DAKO | K3468 |
| 3-Aminopropyltriethoxysilane (APES) | – | 2% | Slide coating | Sigma-Aldrich | A3648 |
| Fluoromount-G, with DAPI | – | – | Histology | ThermoFisher | 00-4959-52 |
| Human Amyloid beta 42 ELISA Kit | – | – | ELISA | ThermoFisher | KHB3441 |
| 4–20% precast polyacrylamide gels | – | – | WB | Bio-Rad | 4561094 |
| **Software and algorithms** |  |  |  |  |  |
| GraphPad Prism (Version 10.0.3) | GraphPad |  |  | http://www.graphpad.com; RRID: SCR_002798 |  |
| Zeiss Zen Lite | Zeiss |  |  | https://www.zeiss.com/microscopy/us/products/software/zeiss-zen-lite.html |  |
| Fiji ImageJ (Version 2.14.0/1.54f/Java 1.8.0-322) | Fiji |  |  | open-source download |  |

|  |  |  |
| --- | --- | --- |
| Image Studio software | Licorbio | https://www.licorbio.com/image-studio |
| Metascape | https://metascape.org/ | open source |
| Cytoscape 3.10.2 | https://cytoscape.org/ | open source |
| ClustVis | https://biit.cs.ut.ee/clustvis/ | open source |
| Ingenuity Pathway Analysis (IPA) | Qiagen | https://digitalinsights.qiagen.com/ |
| SynGO | https://syngopotal.org/ | open source |
| String 12.0 | https://string-db.org/ | open source |
| **Deposited data** |  |  |
| Mass spectrometry dataset | ProteomeXchange Consortium via the PRIDE<br>https://www.ebi.ac.uk/pride/archive/projects/PXD040225<br>https://ftp.pride.ebi.ac.uk/pride/data/archive/2025/10/PXD040225 | PXD040225 |
| Machine learning | [Scikit-learn](https://scikit-learn.org/stable/index.html)<br>[NumPy](https://numpy.org/)<br>[pandas](https://pandas.pydata.org/)<br>[SciPy](https://scipy.org/)<br>[Jupyter](https://jupyter.org/)<sup>128</sup> | https://scikit-learn.org/stable/index.html<br>https://numpy.org/<br>https://pandas.pydata.org/<br>https://scipy.org/<br>https://jupyter.org/ |

Abbreviations: CitR<sub>209</sub>-tau – Citrullinated arginine (R)-209 tau; Cyanine dyes – Cy2, Cy3, Cy5; HRP – Horseradish peroxidase; IF – immunofluorescence; mAb – monoclonal antibody; NFT – Neurofibrillary tangles; NGFR – Nerve growth factor receptor; NMDAR2A – N-methyl-D-aspartate receptor; pAb – polyclonal antibody; PHF – Paired helical filament; PSD95 – Postsynaptic density 95; UCHL-1 – Ubiquitin carboxyl-terminal hydrolase L1; VGLUT-1 – Vesicular glutamate transporter.

## EXPERIMENTAL MODEL AND STUDY PARTICIPANT DETAILS

### Postmortem Human eye and brain samples

Postmortem human eye globes and brain tissues were obtained from the Alzheimer’s Disease Research Center (ADRC) Neuropathology Core in the Department of Pathology at the University of Southern California (USC, Los Angeles, CA; IRB protocol HS-042071). In addition, eye globes were obtained from the National Disease Research Interchange (NDRI, Philadelphia, PA; IRB exempt protocol EX-1055). For a subset of patients and controls, brain specimens were also obtained from the ADRC Neuropathology Core at the University of California, Irvine (UCI [IRB protocol HS#2014–1526]). USC-ADRC, NDRI, and UCI ADRC maintain human tissue collection protocols that are approved by their respective governing committees and are subject to oversight by the National Institutes of Health and institutional guidelines. All the histological procedures were conducted at Cedars-Sinai Medical Center under IRB protocols (Pro00053412 and Pro00019393, Pro00055802). For histological examinations, 53 retinas were collected from deceased donors with confirmed AD (n = 21) or MCI due to AD (n = 11), and from age- and sex-matched deceased donors with NC (n = 21). For analysis of retinal proteins using mass spectrometry (MS), Enzyme-Linked Immunosorbent Assay (ELISA) and Western blot (WB), fresh-frozen retinas were collected from another cohort of deceased donors (n = 16) with clinically and neuropathologically confirmed AD patients (n = 9) and matched NC controls (n = 7). Comprehensive cohort details are presented in Table 1, and Table S1. The human cohort used in this study exhibited no significant differences in age, sex, or post-mortem interval (PMI) hours. Patients’ confidentiality was maintained by de-identifying all tissue samples, ensuring that donors could not be identified.

### Clinical and neuropathological assessments

The detailed clinical and neuropathological assessment procedures are described in our recent publication^28,37^. In summary, clinical and neuropathological reports detailing patients’ neurological examinations and neuropsychological and cognitive assessments were generously provided by the ADRC system using through the Unified Data Set^129^. The NDRI provided patient information, including sex, ethnicity, age at death, cause of death, medical history indicating AD, the presence or absence of dementia, and any accompanying medical conditions. Most cognitive assessments were conducted annually, typically within one year prior to death. In this study, we utilized cognitive scores obtained closest to the patient’s time of death. Three global indicators of cognitive status were used for clinical assessment: the Clinical Dementia Rating (CDR scores: 0 = normal; 0.5 = very mild impairment; 1 = mild dementia; 2 = moderate dementia; or 3 = severe dementia)^130^, Montreal Cognitive Assessment (MOCA scores: ≥26 = cognitively normal or <26 = cognitively impaired^131,132^, and the Mini-Mental State Examination (MMSE scores: normal cognition = 24 - 30; MCI = 20 - 23; moderate dementia = 10 - 19; or severe dementia ≤ 9)^133^.

The assessment of cerebral Aβ burden comprised analysis of diffuse and neuritic plaques (including both immature and mature forms), along with amyloid angiopathy, neurofibrillary tangles (NFTs), neuritic threads (NTs), granulovacuolar degeneration, Lewy bodies, Hirano bodies, Pick bodies, ballooned neurons, neuronal loss, microvascular changes, and gliosis. These evaluations were conducted across multiple brain regions, notably in the hippocampus, the entorhinal cortex, the superior frontal gyrus in the frontal lobe, the superior temporal gyrus in the temporal lobe, the superior parietal lobule in the parietal lobe, the primary visual cortex, and the visual association area in the occipital lobe. All brain samples were uniformly collected by a neuropathologist.

Formalin-fixed, paraffin-embedded brain sections were used to determine the severity of amyloid plaques and NFTs in the brain using anti Aβ monoclonal antibody (mAb) clone 4G8, anti phospho-tau mAb clone AT8, Thioflavin-S (ThioS), and Gallyas silver staining. Two neuropathologists independently rated the burden of Aβ, NFTs, and NTs on a scale of 0, 1, 3, and 5 [(0 = none, 1 = sparse (0-5), 3 = moderate (6-20), 5 = abundant/frequent (21-30 or above), N/A = not applicable)], with final scores calculated as the average of the two readings. The final diagnosis included AD neuropathologic change. The Aβ plaque scoring system, was adapted from Tal et al., (A0 = no Aβ or amyloid plaques, A1 = Thal phase 1 or 2, A2 = Thal phase 3, and A3 = Thal phase 4 or 5)^134^. NFT staging was adjusted from Braak for silver-based histochemistry or p-tau immunohistochemistry (B0 = No NFTs, B1 = Braak stage I or II, B2 = Braak stage III or IV, B3 = Braak stage V or VI)^135^. The neuritic plaque score was adapted from CERAD (C0 = no neuritic plaques, C1 = CERAD sparse, C2 = CERAD moderate, C3 = CERAD frequent)^136^. Additional evaluations included neuronal loss, gliosis, granulovacuolar degeneration, Hirano bodies, Lewy bodies, Pick bodies, and ballooned neurons, using hematoxylin and eosin staining, with scores of 0 for absent and 1 for present. Amyloid angiopathy was classified into 4 grades: Grade I indicates amyloid around normal/atrophic smooth muscle cells of vessels; Grade II shows media replaced by amyloid without blood leakage; Grade III involves extensive amyloid deposition with vessel wall fragmentation and perivascular leakage; Grade IV includes extensive amyloid deposition with fibrinoid necrosis, microaneurysms, mural thrombi, lumen inflammation, and perivascular neuritis.

## METHOD DETAILS

### Collection and processing of eyes tissues

Donor eyes were obtained within an average of 7.9 hours from the time of death. For histological and TEM analyses, eyes were punctured once at the limbus, fixed in either 10% neutral buffered formalin or 4% paraformaldehyde (PFA), and stored at 4°C. Freshly collected eyes for biochemical analyses were preserved in Optisol-GS media (Bausch & Lomb, 50006-OPT) and stored at 4°C for less than 36 hours, after which they were snap frozen and stored at −80°C. These procedures were uniformly performed across all tissue sources and collection sites.

### Preparation of retinal tissues and cross-sections

Eyes that were either freshly preserved in Optisol-GS or fixed were dissected on ice. After removal of the anterior chamber to obtain eyecups, the vitreous humor was carefully cleared. Retinas were then meticulously isolated, separated from the choroid, and prepared as flatmounts following established procedures^27,28,37^. Flatmount strips (∼2mm wide) extending from the ora serrata to the optic disc were dissected to generate four strips: superior-temporal (ST), inferior-temporal (IT), inferior-nasal (IN), and superior-nasal (SN). The ST and IT strips analyzed in this study, generated from fixed retinas, were embedded in paraffin, rotated 90° horizontally, and re-embedded in paraffin blocks. These retinal strips, encompassing central, mid, and far retinal subregions, were sectioned at 8-10 µm and mounted on microscope slides treated with 3-Aminopropyltriethoxysilane (APES, Sigma A3648). Temporal hemiretinas dissected from fresh tissue was stored at −80 °C for downstream protein analysis (MS, ELISA, WB). Together, these preparation methods provided consistent and comprehensive access to retinal quadrants, layers, and pathological subregions.

### Immunohistochemical analysis (IHC)

Before the IHC procedure, paraffin-embedded retinal cross-section slides were deparaffinized in 100% xylene twice (10 min each), rehydrated through decreasing concentrations of ethanol (100% to 70%), and washed with distilled water followed by PBS. Antigen retrieval was performed by incubating retinal cross-sections in target retrieval solution (pH 6.1; S1699, DAKO) at 99 °C for 1 h, followed by PBS washes. In some staining cases, sections were additionally treated with 70% formic acid (ACROS) for 10 min at room temperature (RT).

#### Peroxidase-based IHC

labelling was performed using the Vectastain Elite ABC HRP kit (Vector, PK-6101, Peroxidase rabbit IgG) according to the manufacturer’s instructions. In summary, after incubation with 3% H_2_O_2_ for 20 min, sections were washed in PBS and incubated with blocking serum containing 0.25% Triton X-100 (Sigma, T8787) for 45 min at RT. Primary antibodies (key resources table) were diluted in PBS containing blocking serum and applied overnight at 4 °C. The following day, sections were washed three times in PBS, incubated with secondary antibody for 30 min at 37°C, washed again three times in PBS and incubated with ABC reagent for 30 min at RT. After PBS washes, signal was visualized with 3,3′-diaminobenzidine (DAB) substrate (DAKO K3468). Hematoxylin counterstaining was performed, and slides were mounted with Paramount aqueous mounting medium (DAKO, S3025). Control sections were processed without the primary antibodies to assess nonspecific labeling.

#### Fluorescence-based immunostaining

sections were treated with blocking solution (DAKO X0909) containing 0.25% Triton X-100 (Sigma, T8787) for 1 h at RT, followed by overnight incubation at 4°C with primary antibodies diluted in the same blocking solution. The following day, after PBS washes, sections were incubated with fluorophore-conjugated secondary antibodies (key resources table) for 1 hour at RT. After PBS washes, sections were mounted using Fluoromount-G, with DAPI (Invitrogen, #00-4959-52). Control sections processed without the primary antibodies were used to assess nonspecific fluorescence.

### Nissl staining

Neuronal cytoplasm was specifically stained with the Nissl staining technique. Deparaffinized and rehydrated sections were stained in 0.1% Cresyl Violet acetate (Sigma #C5042) for 5 min, rapidly rinsed in tap water, and briefly dipped in 70% ethanol. Sections were then dehydrated through 2 changes of absolute ethanol for 3 min each, followed by immersion in xylene twice for 2 min, and mounted using mounting medium xylene (Fisher scientific company, L.L.C. #245–691). An average of 12 images per section, covering the retinal neurons from the optic disc to the ora serrata, were captured at a 20x objective and analyzed to quantify the percent neuronal area in the ONL and INL.

### Dihydroethidium (DHE) labeling

Tissues were fluorescently labeled with DHE (Sigma-Aldrich #D7008) for 30 min at 37°C (10 μM final concentration in PBS) prior to mounting.

### Microscopy and Quantitative immunohistochemistry

Fluorescence and bright field images were acquired using a Carl Zeiss Axio Imager Z1 fluorescence microscope with ZEN 2.6 blue edition software (Carl Zeiss MicroImaging, Inc.) and equipped with ApoTome, AxioCam MRm, and AxioCam HRc cameras. Multi-channel image acquisition was used to create images with several channels. Images were repeatedly captured at 20 × (resolution of 1388 × 1040 pixels, 447.63 µm × 335.30 µm /per image) or 40 × (resolution of 1388 × 1040 pixels, 223.82 µm × 167.70 µm /per image) at the same focal planes with the same exposure time for each marker and human donor. We randomly acquired 3 images from the central, 4 from the mid-, and 3 from the far-retinal subregions, for analytical purposes. Images were exported to Fiji ImageJ (version 2.14.0) to analyze parameters of interest. Acquired images were converted to grayscale and standardized to baseline by using a histogram-based threshold in Fiji ImageJ. This baseline-derived threshold was then applied uniformly to the corresponding single channel for all subjects across diagnostic groups. Images were subsequently subjected to ImageJ2/Fiji particle analysis for each biomarker to determine total and % immunoreactive area. Thickness measurements in µm (from the inner limiting membrane to the outer limiting membrane) were manually assessed using ZEN 2.6 Blue Edition software. Throughout the analysis process, the researchers were blinded to each patient’s diagnosis.

### Transmission Electron Microscopy (TEM)

Tissue samples were initially fixed in 4% paraformaldehyde (PFA) in phosphate-buffered saline (PBS) for 20 min at room temperature (RT), followed by three washes in PBS (10 min each). For intracellular antigen labeling, samples were permeabilized and blocked in PBS containing 1% bovine serum albumin (BSA) and 0.1% Triton X-100 for 15 min at RT, then rinsed in PBS. Primary antibodies against PSD95 and VGLUT1 were diluted in blocking solution (1% BSA in PBS) and applied overnight at 4°C in a humidified chamber. Samples were then washed three times with PBS (5 min each). Secondary detection was performed using species-specific, gold-conjugated secondary antibodies (anti-rabbit, 6 nm; anti–guinea pig, 12 nm), diluted in PBS containing 1% BSA. Samples were incubated for 1 hour at RT, followed by three additional PBS washes (5 min each). Subsequently, samples were post-fixed in half-strength Karnovsky’s fixative (2% paraformaldehyde, 2.5% glutaraldehyde in 0.1 M cacodylate buffer, pH 7.2) for 15 min. Following post-fixation, the samples were rinsed in 0.1 M cacodylate buffer (pH 7.2), then post-fixed in 2% osmium tetroxide for 1 hour, rinsed again in cacodylate buffer, and subsequently in sodium acetate. En bloc staining was performed using 1% aqueous uranyl acetate for 1 hour, followed by a wash in sodium acetate. Samples were then dehydrated through a graded series of ethanol (50%, 70%, 85%, 95%, and 100%), followed by incubation in 1:1 and 1:2 mixtures of ethanol and propylene oxide (PO), and finally in pure PO. Tissues were infiltrated with PO/Eponate resin mixtures and embedded in 100% Eponate resin. Polymerization was carried out at 60°C for 48 hours. Ultrathin sections (60 nm) were obtained using an ultramicrotome and collected on copper grids. Sections were imaged using a JEOL JEM-2100 LaB6 transmission electron microscope operating at 120 kV (JEOL USA). Digital images were captured with an Orius SC1000B CCD camera (Gatan).

### Proteome analysis by mass spectrometry (MS)

In this study, we performed secondary analyses on previously published retinal mass spectrometry data ^28^, to identify DEPs associated with synaptic function and integrity. In brief, this analysis included the following steps: (1) preparation of retinal and brain homogenates; (2) tandem mass tag labeling; (3) nanoflow liquid chromatography electrospray ionization tandem mass spectrometry; and (4) database searching, peptide quantification, and statistical analysis. Given the exploratory nature and limited sample size of this human retinal dataset, we adopted a less stringent threshold, defining DEPs as proteins with an FDR-adjusted P < 0.20 and |FC| > 1.2 (or, in selected sub-analyses, an unadjusted P < 0.05 and |FC| > 1.2, as specified for each analysis) for downstream interpretation.

#### Functional Network and Computational Analysis

prediction of activation or inhibition of pathways related to retinal degeneration, synaptic function and integrity, and neurotransmission were analyzed with Ingenuity Pathway Analysis (IPA, Qiagen; https://digitalinsights.qiagen.com). Activation z-scores, P-values and Benjamini-Hochberg adjusted P-values are reported. Volcano plots representing expression changes [log_2_(FC)] and significance levels [-log_10_(P)] in AD versus NC was created using Prism 10.3.1 (GraphPad) and included human protein related to synapse. The list of synaptic-associated proteins was extracted from SunGO release 1.2 knowledge base. Gene Ontology (GO) analysis of synapse-related differentially expressed proteins (DEPs) was performed in Metascape (https://metascape.org/; cutoffs: overlap ≥ 3, *P*-value < 0.01, enrichment ≥ 1.5) and included the GO Biological Processes, Reactome, Kyoto Encyclopedia of Genes and Genomes (KEGG) and WikiPathways databases. Pathway networks were created in Metascape and subsequently loaded and modified in Cytoscape 3.10.2 (https://cytoscape.org/). Protein interaction networks were generated in String v12.0 and modified in Cytoscape. Heatmaps corresponding to the protein expression level of select proteins in the retina of the 6 NC individuals and the 6 AD patients, standardized by unit variance scaling, were generated in ClustVis (https://biit.cs.ut.ee/clustvis/). The mass spectrometry proteomics data have been deposited to the ProteomeXchange Consortium via the PRIDE^137^ partner repository under the dataset identifier PXD040225 (https://www.ebi.ac.uk/pride/archive/projects/PXD040225; https://ftp.pride.ebi.ac.uk/pride/data/archive/2025/10/PXD040225).

#### Generation of Gene-level Ranking Scores and Gene Set Enrichment Analysis

Differential protein abundance statistics were processed to generate a continuous, directionally signed ranking metric suitable for preranked gene set enrichment analysis (GSEA)^85^. For each quantified protein, the log fold-change direction (positive or negative) and the raw P-value from the comparison of interest were used to compute a score defined as: rank_score=sign(logFC)×[−log10(P)]. This metric integrates the magnitude and direction of differential abundance with the statistical strength of the evidence, producing a continuous distribution of values that preserves ordering across proteins. Proteins lacking a gene symbol or P-value were removed. When multiple entries mapped to the same gene symbol, scores were averaged to produce a single representative value per gene. The resulting gene-level scores were sorted in descending order to generate the ranked list required for preranked GSEA.

Pathway enrichment analysis was performed using the *gseapy* Python package (https://github.com/zqfang/gseapy), which implements the Broad Institute’s GSEA algorithm. The prerank module was run with 1,000 permutations, a minimum gene set size of 15, and a maximum gene set size of 500. Gene sets were obtained from the designated human pathway collections (e.g., KEGG_2021_Human, GO Biological Process, or WikiPathways, depending on the analysis). Normalized enrichment scores (NES), nominal P-values, and false discovery rate (FDR) values were extracted from the gseapy output. Selected pathways are shown. Enrichment plots and ranked-metric distributions were generated directly using gseapy’s plotting utilities.

### Biochemical determination of Aβ_1–42_ levels by sandwich ELISA in human retina

In this study, we reused retinal Aβ_1–42_ data generated in our previous study^28^ to perform pairwise correlation analyses with newly identified differentially expressed proteins. Fresh-frozen human retinal strips from the temporal hemisphere were homogenized as previously described^27,28^. The amount of Aβ_1–42_ in retinal homogenates was determined using an anti-human Aβ_1–42_ end-specific sandwich ELISA kit (Thermo Fisher, KHB3441) and normalized to the total protein concentration (Thermo Fisher Scientific).

#### Western blot analysis of human retina

Fresh-frozen postmortem temporal hemiretina tissues from human donors were homogenized in radioimmunoprecipitation assay (RIPA) buffer (0.5M Tris-HCl, pH 7.4, 1.5M NaCl, 2.5% deoxycholic acid, 10% NP-40, 10mM EDTA, Millipore; 20–188) supplemented with protease and phosphatase inhibitors. Protein concentration was determined using a bicinchoninic acid protein assay kit (Pierce^TM^). Lysates were cleared with brief centrifugation for 10 min at 8000g, normalized, and boiled at 95°C after addition of 6X SDS loading dye. Equal amounts of protein (30 µg per sample) were separated on 4–20% precast polyacrylamide gels (Bio-Rad, catalog #4561094) and transferred to polyvinylidene difluoride membranes. The membranes were then blocked with 2.5% bovine serum albumin in 1x TBS-T (Tris-buffered saline with 0.1% Tween-20) for 1 h at RT, followed by overnight incubation at 4 °C with the anti-UCHL1, anti-PSD95, anti-synaptophysin, and anti-p75NTR primary antibody (key resources table). After washing with 1x TBS-T, the membrane was incubated with fluorescently labeled secondary antibodies, and bands were detected using the LI-COR Odyssey imaging system. Band intensities were quantified using Image Studio software (LI-COR), and relative protein expression levels were calculated by normalizing target protein signals to GAPDH. In some cases, membranes were re-probed with primary antibodies after stripping with 1X stripping buffer (NewBlot^TM^ Nitro stripping buffer, Licorbio #928-40030).

#### Machine learning prediction

The dataset was split into training (80%, 38/14/44 NC/MCI/AD) and test sets (20%, 10/4/10 NC/MCI/AD), stratified by clinical diagnosis and patient sex. Two random forest regression models were trained to predict Braak stage and MMSE using 80 estimators and L1 loss, with both models using the same set of retinal biomarkers for their predictions. Due to random forest models being sensitive to their initial conditions, we instantiated the models using 1,000 different random seeds to identify which features were robustly predictive of their target based on their SHAP values^138^. At each iteration, the top features sorted by mean absolute SHAP were tabulated. After all iterations, these features were gathered and the fraction of iterations where they were present was computed (1.0 signifies the feature was in all iterations; 0.5 in half). Learning curves were produced using 10 steps between 10% and 100% of data; cross-validation scores were obtained using 5-fold cross-validation and taking the mean across folds.

## QUANTIFICATION AND STATISTICAL ANALYSIS

All statistical analyses were conducted using GraphPad Prism 10.0.3 (GraphPad Software). Data are presented as mean ± standard error of the mean (SEM) for bar graphs, or as lower quartile, median, and upper quartile values for violin plots. Comparisons among three or more groups were performed using one-way or two-way analysis of variance (ANOVA), followed by Tukey’s post hoc test for multiple comparisons. Comparisons between two independent groups were assessed using two-tailed unpaired Student’s t-tests. Associations between continuous variables were evaluated using Pearson’s correlation coefficient (*r_p_*) for normally distributed data or Spearman’s rank correlation coefficient (*r_s_*) for non-parametric data. Pairwise correlation *r_p_ or r_s_* and P values were reported to indicate the direction, strength, and significance of the linear relationship. For multivariable correlation analyses, P values were further adjusted for multiple comparisons using the Holm–Šídák method, and P*_adj._* are reported. A P value of <0.05 was considered statistically significant. Where applicable, fold changes (FC) and corresponding 95% confidence intervals (CI) were calculated to quantify the magnitude and precision of observed effects.

## References

1. Arendt, T. (2009). Synaptic degeneration in Alzheimer’s disease. Acta Neuropathol 118, 167–179. 10.1007/s00401-009-0536-x.

2. Selkoe, D.J. (2002). Alzheimer’s disease is a synaptic failure. Science 298, 789–791. 10.1126/science.1074069.

3. Spires-Jones, T.L., and Hyman, B.T. (2014). The intersection of amyloid beta and tau at synapses in Alzheimer’s disease. Neuron 82, 756–771. 10.1016/j.neuron.2014.05.004.

4. de Paula, V.J.R., Guimaraes, F.M., Diniz, B.S., and Forlenza, O.V. (2009). Neurobiological pathways to Alzheimer’s disease: Amyloid-beta, TAU protein or both? Dement Neuropsychol 3, 188–194. 10.1590/S1980-57642009DN30300003.

5. Grundke-Iqbal, I., Iqbal, K., Tung, Y.C., Quinlan, M., Wisniewski, H.M., and Binder, L.I. (1986). Abnormal phosphorylation of the microtubule-associated protein tau (tau) in Alzheimer cytoskeletal pathology. Proc Natl Acad Sci U S A 83, 4913–4917. 10.1073/pnas.83.13.4913.

6. Hampel, H., Hardy, J., Blennow, K., Chen, C., Perry, G., Kim, S.H., Villemagne, V.L., Aisen, P., Vendruscolo, M., Iwatsubo, T., et al. (2021). The Amyloid-beta Pathway in Alzheimer’s Disease. Mol Psychiatry 26, 5481–5503. 10.1038/s41380-021-01249-0.

7. Kosik, K.S., Joachim, C.L., and Selkoe, D.J. (1986). Microtubule-associated protein tau (tau) is a major antigenic component of paired helical filaments in Alzheimer disease. Proc Natl Acad Sci U S A 83, 4044–4048. 10.1073/pnas.83.11.4044.

8. Masters, C.L., Simms, G., Weinman, N.A., Multhaup, G., McDonald, B.L., and Beyreuther, K. (1985). Amyloid plaque core protein in Alzheimer disease and Down syndrome. Proc Natl Acad Sci U S A 82, 4245–4249. 10.1073/pnas.82.12.4245.

9. Yarns, B.C., Holiday, K.A., Carlson, D.M., Cosgrove, C.K., and Melrose, R.J. (2022). Pathophysiology of Alzheimer’s Disease. Psychiatr Clin North Am 45, 663–676. 10.1016/j.psc.2022.07.003.

10. Anggono, V., Tsai, L.H., and Gotz, J. (2016). Glutamate Receptors in Alzheimer’s Disease: Mechanisms and Therapies. Neural Plast 2016, 8256196. 10.1155/2016/8256196.

11. Benarroch, E.E. (2018). Glutamatergic synaptic plasticity and dysfunction in Alzheimer disease: Emerging mechanisms. Neurology 91, 125–132. 10.1212/WNL.0000000000005807.

12. Colom-Cadena, M., Spires-Jones, T., Zetterberg, H., Blennow, K., Caggiano, A., DeKosky, S.T., Fillit, H., Harrison, J.E., Schneider, L.S., Scheltens, P., et al. (2020). The clinical promise of biomarkers of synapse damage or loss in Alzheimer’s disease. Alzheimers Res Ther 12, 21. 10.1186/s13195-020-00588-4.

13. Griffiths, J., and Grant, S.G.N. (2023). Synapse pathology in Alzheimer’s disease. Semin Cell Dev Biol 139, 13–23. 10.1016/j.semcdb.2022.05.028.

14. Kumar, S., Ramos, E., Hidalgo, A., Rodarte, D., Sharma, B., Torres, M.M., Devara, D., and Gadad, S.S. (2025). Integrated multi-omics analyses of synaptosomes revealed synapse-associated novel targets in Alzheimer’s disease. Mol Psychiatry. 10.1038/s41380-025-03095-w.

15. Masliah, E., Alford, M., DeTeresa, R., Mallory, M., and Hansen, L. (1996). Deficient glutamate transport is associated with neurodegeneration in Alzheimer’s disease. Ann Neurol 40, 759–766. 10.1002/ana.410400512.

16. Meftah, S., and Gan, J. (2023). Alzheimer’s disease as a synaptopathy: Evidence for dysfunction of synapses during disease progression. Front Synaptic Neurosci 15, 1129036. 10.3389/fnsyn.2023.1129036.

17. Zott, B., and Konnerth, A. (2023). Impairments of glutamatergic synaptic transmission in Alzheimer’s disease. Semin Cell Dev Biol 139, 24–34. 10.1016/j.semcdb.2022.03.013.

18. Li, S., Hayden, E.Y., Garcia, V.J., Fuchs, D.T., Sheyn, J., Daley, D.A., Rentsendorj, A., Torbati, T., Black, K.L., Rutishauser, U., et al. (2020). Activated Bone Marrow-Derived Macrophages Eradicate Alzheimer’s-Related Abeta(42) Oligomers and Protect Synapses. Front Immunol 11, 49. 10.3389/fimmu.2020.00049.

19. Cao, K.J., Kim, J.H., Kroeger, H., Gaffney, P.M., Lin, J.H., Sigurdson, C.J., and Yang, J. (2021). ARCAM-1 Facilitates Fluorescence Detection of Amyloid-Containing Deposits in the Retina. Transl Vis Sci Technol 10, 5. 10.1167/tvst.10.7.5.

20. Cao, Q., Yang, S., Wang, X., Sun, H., Chen, W., Wang, Y., Gao, J., Wu, Y., Yang, Q., Chen, X., et al. (2024). Transport of beta-amyloid from brain to eye causes retinal degeneration in Alzheimer’s disease. J Exp Med 221. 10.1084/jem.20240386.

21. Davis, M.R., Robinson, E., Koronyo, Y., Salobrar-Garcia, E., Rentsendorj, A., Gaire, B.P., Mirzaei, N., Kayed, R., Sadun, A.A., Ljubimov, A.V., et al. (2025). Retinal ganglion cell vulnerability to pathogenic tau in Alzheimer’s disease. Acta Neuropathol Commun 13, 31. 10.1186/s40478-025-01935-y.

22. den Haan, J., Morrema, T.H.J., Verbraak, F.D., de Boer, J.F., Scheltens, P., Rozemuller, A.J., Bergen, A.A.B., Bouwman, F.H., and Hoozemans, J.J. (2018). Amyloid-beta and phosphorylated tau in post-mortem Alzheimer’s disease retinas. Acta Neuropathol Commun 6, 147. 10.1186/s40478-018-0650-x.

23. Gaire, B.P., Koronyo, Y., Fuchs, D.T., Shi, H., Rentsendorj, A., Danziger, R., Vit, J.P., Mirzaei, N., Doustar, J., Sheyn, J., et al. (2024). Alzheimer’s disease pathophysiology in the Retina. Prog Retin Eye Res 101, 101273. 10.1016/j.preteyeres.2024.101273.

24. Grimaldi, A., Pediconi, N., Oieni, F., Pizzarelli, R., Rosito, M., Giubettini, M., Santini, T., Limatola, C., Ruocco, G., Ragozzino, D., and Di Angelantonio, S. (2019). Neuroinflammatory Processes, A1 Astrocyte Activation and Protein Aggregation in the Retina of Alzheimer’s Disease Patients, Possible Biomarkers for Early Diagnosis. Front Neurosci 13, 925. 10.3389/fnins.2019.00925.

25. Hart de Ruyter, F.J., Evers, M., Morrema, T.H.J., Dijkstra, A.A., den Haan, J., Twisk, J.W.R., de Boer, J.F., Scheltens, P., Bouwman, F.H., Verbraak, F.D., et al. (2024). Neuropathological hallmarks in the post-mortem retina of neurodegenerative diseases. Acta Neuropathol 148, 24. 10.1007/s00401-024-02769-z.

26. Hart de Ruyter, F.J., Morrema, T.H.J., den Haan, J., Netherlands Brain, B., Twisk, J.W.R., de Boer, J.F., Scheltens, P., Boon, B.D.C., Thal, D.R., Rozemuller, A.J., et al. (2023). Phosphorylated tau in the retina correlates with tau pathology in the brain in Alzheimer’s disease and primary tauopathies. Acta Neuropathol 145, 197–218. 10.1007/s00401-022-02525-1.

27. Koronyo, Y., Biggs, D., Barron, E., Boyer, D.S., Pearlman, J.A., Au, W.J., Kile, S.J., Blanco, A., Fuchs, D.T., Ashfaq, A., et al. (2017). Retinal amyloid pathology and proof-of-concept imaging trial in Alzheimer’s disease. JCI Insight 2. 10.1172/jci.insight.93621.

28. Koronyo, Y., Rentsendorj, A., Mirzaei, N., Regis, G.C., Sheyn, J., Shi, H., Barron, E., Cook-Wiens, G., Rodriguez, A.R., Medeiros, R., et al. (2023). Retinal pathological features and proteome signatures of Alzheimer’s disease. Acta Neuropathol 145, 409–438. 10.1007/s00401-023-02548-2.

29. Koronyo-Hamaoui, M., Koronyo, Y., Ljubimov, A.V., Miller, C.A., Ko, M.K., Black, K.L., Schwartz, M., and Farkas, D.L. (2011). Identification of amyloid plaques in retinas from Alzheimer’s patients and noninvasive in vivo optical imaging of retinal plaques in a mouse model. Neuroimage 54 *Suppl 1*, S204–217. 10.1016/j.neuroimage.2010.06.020.

30. La Morgia, C., Ross-Cisneros, F.N., Koronyo, Y., Hannibal, J., Gallassi, R., Cantalupo, G., Sambati, L., Pan, B.X., Tozer, K.R., Barboni, P., et al. (2016). Melanopsin retinal ganglion cell loss in Alzheimer disease. Ann Neurol 79, 90–109. 10.1002/ana.24548.

31. Lee, S., Jiang, K., McIlmoyle, B., To, E., Xu, Q.A., Hirsch-Reinshagen, V., Mackenzie, I.R., Hsiung, G.R., Eadie, B.D., Sarunic, M.V., et al. (2020). Amyloid Beta Immunoreactivity in the Retinal Ganglion Cell Layer of the Alzheimer’s Eye. Front Neurosci 14, 758. 10.3389/fnins.2020.00758.

32. Nunez-Diaz, C., Andersson, E., Schultz, N., Poceviciute, D., Hansson, O., Netherlands Brain, B., Nilsson, K.P.R., and Wennstrom, M. (2024). The fluorescent ligand bTVBT2 reveals increased p-tau uptake by retinal microglia in Alzheimer’s disease patients and App(NL-F/NL-F) mice. Alzheimers Res Ther 16, 4. 10.1186/s13195-023-01375-7.

33. Santiago, J., Poceviciute, D., Vogel, J., Netherlands Brain, B., Brinkmalm, G., and Wennstrom, M. (2025). Retinal tau phosphorylation in Alzheimer’s disease: A mass spectrometry study. Neurobiol Dis 215, 107057. 10.1016/j.nbd.2025.107057.

34. Schultz, N., Byman, E., Netherlands Brain, B., and Wennstrom, M. (2020). Levels of Retinal Amyloid-beta Correlate with Levels of Retinal IAPP and Hippocampal Amyloid-beta in Neuropathologically Evaluated Individuals. J Alzheimers Dis 73, 1201–1209. 10.3233/JAD-190868.

35. Shi, H., Koronyo, Y., Fuchs, D.T., Sheyn, J., Jallow, O., Mandalia, K., Graham, S.L., Gupta, V.K., Mirzaei, M., Kramerov, A.A., et al. (2023). Retinal arterial Abeta(40) deposition is linked with tight junction loss and cerebral amyloid angiopathy in MCI and AD patients. Alzheimers Dement 19, 5185–5197. 10.1002/alz.13086.

36. Shi, H., Koronyo, Y., Rentsendorj, A., Regis, G.C., Sheyn, J., Fuchs, D.T., Kramerov, A.A., Ljubimov, A.V., Dumitrascu, O.M., Rodriguez, A.R., et al. (2020). Identification of early pericyte loss and vascular amyloidosis in Alzheimer’s disease retina. Acta Neuropathol 139, 813–836. 10.1007/s00401-020-02134-w.

37. Shi, H., Mirzaei, N., Koronyo, Y., Davis, M.R., Robinson, E., Braun, G.M., Jallow, O., Rentsendorj, A., Ramanujan, V.K., Fert-Bober, J., et al. (2024). Identification of retinal oligomeric, citrullinated, and other tau isoforms in early and advanced AD and relations to disease status. Acta Neuropathol 148, 3. 10.1007/s00401-024-02760-8.

38. Wijesinghe, P., Hosseini, A., Campbell, M., Tejpal, S., Haynes, J., Xi, J., Mackenzie, I.R., Hirsch-Reinshagen, V., Hsiung, G.R., Spiller, B.W., et al. (2025). Decoding amyloid beta clearance systems at inner blood-retina barrier using three-dimensional ex vivo retinal imaging in Alzheimer’s disease. Alzheimers Dement 21, e70592. 10.1002/alz.70592.

39. Xu, Q.A., Boerkoel, P., Hirsch-Reinshagen, V., Mackenzie, I.R., Hsiung, G.R., Charm, G., To, E.F., Liu, A.Q., Schwab, K., Jiang, K., et al. (2022). Muller cell degeneration and microglial dysfunction in the Alzheimer’s retina. Acta Neuropathol Commun 10, 145. 10.1186/s40478-022-01448-y.

40. Du, X., Koronyo, Y., Mirzaei, N., Yang, C., Fuchs, D.T., Black, K.L., Koronyo-Hamaoui, M., and Gao, L. (2022). Label-free hyperspectral imaging and deep-learning prediction of retinal amyloid beta-protein and phosphorylated tau. PNAS Nexus 1, pgac164. 10.1093/pnasnexus/pgac164.

41. Dumitrascu, O.M., Doustar, J., Fuchs, D.T., Koronyo, Y., Sherman, D.S., Miller, M.S., Johnson, K.O., Carare, R.O., Verdooner, S.R., Lyden, P.D., et al. (2024). Retinal peri-arteriolar versus peri-venular amyloidosis, hippocampal atrophy, and cognitive impairment: exploratory trial. Acta Neuropathol Commun 12, 109. 10.1186/s40478-024-01810-2.

42. Dumitrascu, O.M., Lyden, P.D., Torbati, T., Sheyn, J., Sherzai, A., Sherzai, D., Sherman, D.S., Rosenberry, R., Cheng, S., Johnson, K.O., et al. (2020). Sectoral segmentation of retinal amyloid imaging in subjects with cognitive decline. Alzheimers Dement (Amst) 12, e12109. 10.1002/dad2.12109.

43. Dumitrascu, O.M., Rosenberry, R., Sherman, D.S., Khansari, M.M., Sheyn, J., Torbati, T., Sherzai, A., Sherzai, D., Johnson, K.O., Czeszynski, A.D., et al. (2021). Retinal Venular Tortuosity Jointly with Retinal Amyloid Burden Correlates with Verbal Memory Loss: A Pilot Study. Cells 10. 10.3390/cells10112926.

44. Gonzales, M., Fuchs, D.T., Doustar, J., Koronyo, Y., Sherman, D.S., Lyden, P.D., Sicotte, N.L., Black, K.L., Kremen, S., Dumitrascu, O.M., and Koronyo-Hamaoui, M. (2025). Integrated analysis of in vivo retinal perivascular amyloid imaging and cognition. Alzheimers Dement 21, e70366. 10.1002/alz.70366.

45. Hadoux, X., Hui, F., Lim, J.K.H., Masters, C.L., Pebay, A., Chevalier, S., Ha, J., Loi, S., Fowler, C.J., Rowe, C., et al. (2019). Non-invasive in vivo hyperspectral imaging of the retina for potential biomarker use in Alzheimer’s disease. Nat Commun 10, 4227. 10.1038/s41467-019-12242-1.

46. Lemmens, S., Van Craenendonck, T., Van Eijgen, J., De Groef, L., Bruffaerts, R., de Jesus, D.A., Charle, W., Jayapala, M., Sunaric-Megevand, G., Standaert, A., et al. (2020). Combination of snapshot hyperspectral retinal imaging and optical coherence tomography to identify Alzheimer’s disease patients. Alzheimers Res Ther 12, 144. 10.1186/s13195-020-00715-1.

47. More, S.S., Beach, J.M., McClelland, C., Mokhtarzadeh, A., and Vince, R. (2019). In Vivo Assessment of Retinal Biomarkers by Hyperspectral Imaging: Early Detection of Alzheimer’s Disease. ACS Chem Neurosci 10, 4492–4501. 10.1021/acschemneuro.9b00331.

48. Ngolab, J., Donohue, M., Belsha, A., Salazar, J., Cohen, P., Jaiswal, S., Tan, V., Gessert, D., Korouri, S., Aggarwal, N.T., et al. (2021). Feasibility study for detection of retinal amyloid in clinical trials: The Anti-Amyloid Treatment in Asymptomatic Alzheimer’s Disease (A4) trial. Alzheimers Dement (Amst) 13, e12199. 10.1002/dad2.12199.

49. Poudel, P., Frost, S.M., Eslick, S., Sohrabi, H.R., Taddei, K., Martins, R.N., and Hone, E. (2024). Hyperspectral Retinal Imaging as a Non-Invasive Marker to Determine Brain Amyloid Status. J Alzheimers Dis 100, S131–S152. 10.3233/JAD-240631.

50. Sharafi, S.M., Sylvestre, J.P., Chevrefils, C., Soucy, J.P., Beaulieu, S., Pascoal, T.A., Arbour, J.D., Rheaume, M.A., Robillard, A., Chayer, C., et al. (2019). Vascular retinal biomarkers improves the detection of the likely cerebral amyloid status from hyperspectral retinal images. Alzheimers Dement (N Y) 5, 610–617. 10.1016/j.trci.2019.09.006.

51. Tadokoro, K., Yamashita, T., Kimura, S., Nomura, E., Ohta, Y., Omote, Y., Takemoto, M., Hishikawa, N., Morihara, R., Morizane, Y., and Abe, K. (2021). Retinal Amyloid Imaging for Screening Alzheimer’s Disease. J Alzheimers Dis 83, 927–934. 10.3233/JAD-210327.

52. Dantuma, N.P., and Bott, L.C. (2014). The ubiquitin-proteasome system in neurodegenerative diseases: precipitating factor, yet part of the solution. Front Mol Neurosci 7, 70. 10.3389/fnmol.2014.00070.

53. Gong, B., Radulovic, M., Figueiredo-Pereira, M.E., and Cardozo, C. (2016). The Ubiquitin-Proteasome System: Potential Therapeutic Targets for Alzheimer’s Disease and Spinal Cord Injury. Front Mol Neurosci 9, 4. 10.3389/fnmol.2016.00004.

54. Kwon, Y.T., and Ciechanover, A. (2017). The Ubiquitin Code in the Ubiquitin-Proteasome System and Autophagy. Trends Biochem Sci 42, 873–886. 10.1016/j.tibs.2017.09.002.

55. Maniv, I., Sarji, M., Bdarneh, A., Feldman, A., Ankawa, R., Koren, E., Magid-Gold, I., Reis, N., Soteriou, D., Salomon-Zimri, S., et al. (2023). Altered ubiquitin signaling induces Alzheimer’s disease-like hallmarks in a three-dimensional human neural cell culture model. Nat Commun 14, 5922. 10.1038/s41467-023-41545-7.

56. Mi, Z., and Graham, S.H. (2023). Role of UCHL1 in the pathogenesis of neurodegenerative diseases and brain injury. Ageing Res Rev 86, 101856. 10.1016/j.arr.2023.101856.

57. Oddo, S. (2008). The ubiquitin-proteasome system in Alzheimer’s disease. J Cell Mol Med 12, 363–373. 10.1111/j.1582-4934.2008.00276.x.

58. Tseng, B.P., Green, K.N., Chan, J.L., Blurton-Jones, M., and LaFerla, F.M. (2008). Abeta inhibits the proteasome and enhances amyloid and tau accumulation. Neurobiol Aging 29, 1607–1618. 10.1016/j.neurobiolaging.2007.04.014.

59. Weng, F.L., and He, L. (2021). Disrupted ubiquitin proteasome system underlying tau accumulation in Alzheimer’s disease. Neurobiol Aging 99, 79–85. 10.1016/j.neurobiolaging.2020.11.015.

60. Bilguvar, K., Tyagi, N.K., Ozkara, C., Tuysuz, B., Bakircioglu, M., Choi, M., Delil, S., Caglayan, A.O., Baranoski, J.F., Erturk, O., et al. (2013). Recessive loss of function of the neuronal ubiquitin hydrolase UCHL1 leads to early-onset progressive neurodegeneration. Proc Natl Acad Sci U S A 110, 3489–3494. 10.1073/pnas.1222732110.

61. Cartier, A.E., Djakovic, S.N., Salehi, A., Wilson, S.M., Masliah, E., and Patrick, G.N. (2009). Regulation of synaptic structure by ubiquitin C-terminal hydrolase L1. J Neurosci 29, 7857–7868. 10.1523/JNEUROSCI.1817-09.2009.

62. Gong, B., Cao, Z., Zheng, P., Vitolo, O.V., Liu, S., Staniszewski, A., Moolman, D., Zhang, H., Shelanski, M., and Arancio, O. (2006). Ubiquitin hydrolase Uch-L1 rescues beta-amyloid-induced decreases in synaptic function and contextual memory. Cell 126, 775–788. 10.1016/j.cell.2006.06.046.

63. Saigoh, K., Wang, Y.L., Suh, J.G., Yamanishi, T., Sakai, Y., Kiyosawa, H., Harada, T., Ichihara, N., Wakana, S., Kikuchi, T., and Wada, K. (1999). Intragenic deletion in the gene encoding ubiquitin carboxy-terminal hydrolase in gad mice. Nat Genet 23, 47–51. 10.1038/12647.

64. Qin, B., Chen, X., Wang, F., and Wang, Y. (2024). DUBs in Alzheimer’s disease: mechanisms and therapeutic implications. Cell Death Discov 10, 475. 10.1038/s41420-024-02237-3.

65. Xie, M., Han, Y., Yu, Q., Wang, X., Wang, S., and Liao, X. (2016). UCH-L1 Inhibition Decreases the Microtubule-Binding Function of Tau Protein. J Alzheimers Dis 49, 353–363. 10.3233/JAD-150032.

66. Zhang, M., Cai, F., Zhang, S., Zhang, S., and Song, W. (2014). Overexpression of ubiquitin carboxyl-terminal hydrolase L1 (UCHL1) delays Alzheimer’s progression in vivo. Sci Rep 4, 7298. 10.1038/srep07298.

67. Bamji, S.X., Majdan, M., Pozniak, C.D., Belliveau, D.J., Aloyz, R., Kohn, J., Causing, C.G., and Miller, F.D. (1998). The p75 neurotrophin receptor mediates neuronal apoptosis and is essential for naturally occurring sympathetic neuron death. J Cell Biol 140, 911–923. 10.1083/jcb.140.4.911.

68. Demuth, H., Hosseini, S., Dusedeau, H.P., Dunay, I.R., Korte, M., and Zagrebelsky, M. (2023). Deletion of p75(NTR) rescues the synaptic but not the inflammatory status in the brain of a mouse model for Alzheimer’s disease. Front Mol Neurosci 16, 1163087. 10.3389/fnmol.2023.1163087.

69. Kraemer, B.R., Snow, J.P., Vollbrecht, P., Pathak, A., Valentine, W.M., Deutch, A.Y., and Carter, B.D. (2014). A role for the p75 neurotrophin receptor in axonal degeneration and apoptosis induced by oxidative stress. J Biol Chem 289, 21205–21216. 10.1074/jbc.M114.563403.

70. Meeker, R.B., and Williams, K.S. (2015). The p75 neurotrophin receptor: at the crossroad of neural repair and death. Neural Regen Res 10, 721–725. 10.4103/1673-5374.156967.

71. Singh, K.K., Park, K.J., Hong, E.J., Kramer, B.M., Greenberg, M.E., Kaplan, D.R., and Miller, F.D. (2008). Developmental axon pruning mediated by BDNF-p75NTR-dependent axon degeneration. Nat Neurosci 11, 649–658. 10.1038/nn.2114.

72. Coulson, E.J., May, L.M., Sykes, A.M., and Hamlin, A.S. (2009). The role of the p75 neurotrophin receptor in cholinergic dysfunction in Alzheimer’s disease. Neuroscientist 15, 317–323. 10.1177/1073858408331376.

73. Knowles, J.K., Rajadas, J., Nguyen, T.V., Yang, T., LeMieux, M.C., Vander Griend, L., Ishikawa, C., Massa, S.M., Wyss-Coray, T., and Longo, F.M. (2009). The p75 neurotrophin receptor promotes amyloid-beta(1-42)-induced neuritic dystrophy in vitro and in vivo. J Neurosci 29, 10627–10637. 10.1523/JNEUROSCI.0620-09.2009.

74. Shen, L.L., Li, W.W., Xu, Y.L., Gao, S.H., Xu, M.Y., Bu, X.L., Liu, Y.H., Wang, J., Zhu, J., Zeng, F., et al. (2019). Neurotrophin receptor p75 mediates amyloid beta-induced tau pathology. Neurobiol Dis 132, 104567. 10.1016/j.nbd.2019.104567.

75. Sotthibundhu, A., Sykes, A.M., Fox, B., Underwood, C.K., Thangnipon, W., and Coulson, E.J. (2008). Beta-amyloid(1-42) induces neuronal death through the p75 neurotrophin receptor. J Neurosci 28, 3941–3946. 10.1523/JNEUROSCI.0350-08.2008.

76. Yaar, M., Zhai, S., Pilch, P.F., Doyle, S.M., Eisenhauer, P.B., Fine, R.E., and Gilchrest, B.A. (1997). Binding of beta-amyloid to the p75 neurotrophin receptor induces apoptosis. A possible mechanism for Alzheimer’s disease. J Clin Invest 100, 2333–2340. 10.1172/JCI119772.

77. Yao, X.Q., Jiao, S.S., Saadipour, K., Zeng, F., Wang, Q.H., Zhu, C., Shen, L.L., Zeng, G.H., Liang, C.R., Wang, J., et al. (2015). p75NTR ectodomain is a physiological neuroprotective molecule against amyloid-beta toxicity in the brain of Alzheimer’s disease. Mol Psychiatry 20, 1301–1310. 10.1038/mp.2015.49.

78. Balzamino, B.O., Biamonte, F., Esposito, G., Marino, R., Fanelli, F., Keller, F., and Micera, A. (2014). Characterization of NGF, trkA (NGFR), and p75 (NTR) in Retina of Mice Lacking Reelin Glycoprotein. Int J Cell Biol 2014, 725928. 10.1155/2014/725928.

79. Subirada, P.V., Tovo, A., Vaglienti, M.V., Luna Pinto, J.D., Saragovi, H.U., Sanchez, M.C., Anastasia, A., and Barcelona, P.F. (2023). Etiological Roles of p75(NTR) in a Mouse Model of Wet Age-Related Macular Degeneration. Cells 12. 10.3390/cells12020297.

80. Wei, Y., Wang, N., Lu, Q., Zhang, N., Zheng, D., and Li, J. (2007). Enhanced protein expressions of sortilin and p75NTR in retina of rat following elevated intraocular pressure-induced retinal ischemia. Neurosci Lett 429, 169–174. 10.1016/j.neulet.2007.10.012.

81. Kashani, A., Lepicard, E., Poirel, O., Videau, C., David, J.P., Fallet-Bianco, C., Simon, A., Delacourte, A., Giros, B., Epelbaum, J., et al. (2008). Loss of VGLUT1 and VGLUT2 in the prefrontal cortex is correlated with cognitive decline in Alzheimer disease. Neurobiol Aging 29, 1619–1630. 10.1016/j.neurobiolaging.2007.04.010.

82. Poirel, O., Mella, S., Videau, C., Ramet, L., Davoli, M.A., Herzog, E., Katsel, P., Mechawar, N., Haroutunian, V., Epelbaum, J., et al. (2018). Moderate decline in select synaptic markers in the prefrontal cortex (BA9) of patients with Alzheimer’s disease at various cognitive stages. Sci Rep 8, 938. 10.1038/s41598-018-19154-y.

83. Proctor, D.T., Coulson, E.J., and Dodd, P.R. (2010). Reduction in post-synaptic scaffolding PSD-95 and SAP-102 protein levels in the Alzheimer inferior temporal cortex is correlated with disease pathology. J Alzheimers Dis 21, 795–811. 10.3233/JAD-2010-100090.

84. Sze, C.I., Troncoso, J.C., Kawas, C., Mouton, P., Price, D.L., and Martin, L.J. (1997). Loss of the presynaptic vesicle protein synaptophysin in hippocampus correlates with cognitive decline in Alzheimer disease. J Neuropathol Exp Neurol 56, 933–944. 10.1097/00005072-199708000-00011.

85. Subramanian, A., Tamayo, P., Mootha, V.K., Mukherjee, S., Ebert, B.L., Gillette, M.A., Paulovich, A., Pomeroy, S.L., Golub, T.R., Lander, E.S., and Mesirov, J.P. (2005). Gene set enrichment analysis: a knowledge-based approach for interpreting genome-wide expression profiles. Proc Natl Acad Sci U S A 102, 15545–15550. 10.1073/pnas.0506580102.

86. Costantini, C., Rossi, F., Formaggio, E., Bernardoni, R., Cecconi, D., and Della-Bianca, V. (2005). Characterization of the signaling pathway downstream p75 neurotrophin receptor involved in beta-amyloid peptide-dependent cell death. J Mol Neurosci 25, 141–156. 10.1385/JMN:25:2:141.

87. Hashimoto, Y., Kaneko, Y., Tsukamoto, E., Frankowski, H., Kouyama, K., Kita, Y., Niikura, T., Aiso, S., Bredesen, D.E., Matsuoka, M., and Nishimoto, I. (2004). Molecular characterization of neurohybrid cell death induced by Alzheimer’s amyloid-beta peptides via p75NTR/PLAIDD. J Neurochem 90, 549–558. 10.1111/j.1471-4159.2004.02513.x.

88. Bohm, E.W., Buonfiglio, F., Voigt, A.M., Bachmann, P., Safi, T., Pfeiffer, N., and Gericke, A. (2023). Oxidative stress in the eye and its role in the pathophysiology of ocular diseases. Redox Biol 68, 102967. 10.1016/j.redox.2023.102967.

89. Country, M.W. (2017). Retinal metabolism: A comparative look at energetics in the retina. Brain Res 1672, 50–57. 10.1016/j.brainres.2017.07.025.

90. Viegas, F.O., and Neuhauss, S.C.F. (2021). A Metabolic Landscape for Maintaining Retina Integrity and Function. Front Mol Neurosci 14, 656000. 10.3389/fnmol.2021.656000.

91. Fan, N., Silverman, S.M., Liu, Y., Wang, X., Kim, B.J., Tang, L., Clark, A.F., Liu, X., and Pang, I.H. (2017). Rapid repeatable in vivo detection of retinal reactive oxygen species. Exp Eye Res 161, 71–81. 10.1016/j.exer.2017.06.004.

92. Tzioras, M., McGeachan, R.I., Durrant, C.S., and Spires-Jones, T.L. (2023). Synaptic degeneration in Alzheimer disease. Nat Rev Neurol 19, 19–38. 10.1038/s41582-022-00749-z.

93. Zeng, F., Lu, J.J., Zhou, X.F., and Wang, Y.J. (2011). Roles of p75NTR in the pathogenesis of Alzheimer’s disease: a novel therapeutic target. Biochem Pharmacol 82, 1500–1509. 10.1016/j.bcp.2011.06.040.

94. Zhang, J., Zhang, Y., Wang, J., Xia, Y., Zhang, J., and Chen, L. (2024). Recent advances in Alzheimer’s disease: Mechanisms, clinical trials and new drug development strategies. Signal Transduct Target Ther 9, 211. 10.1038/s41392-024-01911-3.

95. Zhang, Y., Chen, H., Li, R., Sterling, K., and Song, W. (2023). Amyloid beta-based therapy for Alzheimer’s disease: challenges, successes and future. Signal Transduct Target Ther 8, 248. 10.1038/s41392-023-01484-7.

96. Nakamura, T., Oh, C.K., Liao, L., Zhang, X., Lopez, K.M., Gibbs, D., Deal, A.K., Scott, H.R., Spencer, B., Masliah, E., et al. (2021). Noncanonical transnitrosylation network contributes to synapse loss in Alzheimer’s disease. Science 371. 10.1126/science.aaw0843.

97. Tramutola, A., Di Domenico, F., Barone, E., Perluigi, M., and Butterfield, D.A. (2016). It Is All about (U)biquitin: Role of Altered Ubiquitin-Proteasome System and UCHL1 in Alzheimer Disease. Oxid Med Cell Longev 2016, 2756068. 10.1155/2016/2756068.

98. Castegna, A., Aksenov, M., Aksenova, M., Thongboonkerd, V., Klein, J.B., Pierce, W.M., Booze, R., Markesbery, W.R., and Butterfield, D.A. (2002). Proteomic identification of oxidatively modified proteins in Alzheimer’s disease brain. Part I: creatine kinase BB, glutamine synthase, and ubiquitin carboxy-terminal hydrolase L-1. Free Radic Biol Med 33, 562–571. 10.1016/s0891-5849(02)00914-0.

99. Choi, J., Levey, A.I., Weintraub, S.T., Rees, H.D., Gearing, M., Chin, L.S., and Li, L. (2004). Oxidative modifications and down-regulation of ubiquitin carboxyl-terminal hydrolase L1 associated with idiopathic Parkinson’s and Alzheimer’s diseases. J Biol Chem 279, 13256–13264. 10.1074/jbc.M314124200.

100. Donovan, L.E., Higginbotham, L., Dammer, E.B., Gearing, M., Rees, H.D., Xia, Q., Duong, D.M., Seyfried, N.T., Lah, J.J., and Levey, A.I. (2012). Analysis of a membrane-enriched proteome from postmortem human brain tissue in Alzheimer’s disease. Proteomics Clin Appl 6, 201–211. 10.1002/prca.201100068.

101. Guglielmotto, M., Monteleone, D., Vasciaveo, V., Repetto, I.E., Manassero, G., Tabaton, M., and Tamagno, E. (2017). The Decrease of Uch-L1 Activity Is a Common Mechanism Responsible for Abeta 42 Accumulation in Alzheimer’s and Vascular Disease. Front Aging Neurosci 9, 320. 10.3389/fnagi.2017.00320.

102. Kepchia, D., Huang, L., Dargusch, R., Rissman, R.A., Shokhirev, M.N., Fischer, W., and Schubert, D. (2020). Diverse proteins aggregate in mild cognitive impairment and Alzheimer’s disease brain. Alzheimers Res Ther 12, 75. 10.1186/s13195-020-00641-2.

103. Breiman, L. (2001). Random Forests. Machine Learning 45, 5–32. 10.1023/A:1010933404324.

104. Walkiewicz, G., Ronisz, A., Van Ginderdeuren, R., Lemmens, S., Bouwman, F.H., Hoozemans, J.J.M., Morrema, T.H.J., Rozemuller, A.J., Hart de Ruyter, F.J., De Groef, L., et al. (2024). Primary retinal tauopathy: A tauopathy with a distinct molecular pattern. Alzheimers Dement 20, 330–340. 10.1002/alz.13424.

105. Guerrero-Munoz, M.J., Gerson, J., and Castillo-Carranza, D.L. (2015). Tau Oligomers: The Toxic Player at Synapses in Alzheimer’s Disease. Front Cell Neurosci 9, 464. 10.3389/fncel.2015.00464.

106. Lee, A., Kondapalli, C., Virga, D.M., Lewis, T.L., Jr., Koo, S.Y., Ashok, A., Mairet-Coello, G., Herzig, S., Foretz, M., Viollet, B., et al. (2022). Abeta42 oligomers trigger synaptic loss through CAMKK2-AMPK-dependent effectors coordinating mitochondrial fission and mitophagy. Nat Commun 13, 4444. 10.1038/s41467-022-32130-5.

107. Selkoe, D.J. (2008). Soluble oligomers of the amyloid beta-protein impair synaptic plasticity and behavior. Behav Brain Res 192, 106–113. 10.1016/j.bbr.2008.02.016.

108. De Rossi, P., Nomura, T., Andrew, R.J., Masse, N.Y., Sampathkumar, V., Musial, T.F., Sudwarts, A., Recupero, A.J., Le Metayer, T., Hansen, M.T., et al. (2020). Neuronal BIN1 Regulates Presynaptic Neurotransmitter Release and Memory Consolidation. Cell Rep 30, 3520–3535 e3527. 10.1016/j.celrep.2020.02.026.

109. Kwon, S.E., and Chapman, E.R. (2011). Synaptophysin regulates the kinetics of synaptic vesicle endocytosis in central neurons. Neuron 70, 847–854. 10.1016/j.neuron.2011.04.001.

110. Lammertse, H.C.A., Moro, A., Saarloos, I., Toonen, R.F., and Verhage, M. (2022). Reduced dynamin-1 levels in neurons lacking MUNC18-1. J Cell Sci 135. 10.1242/jcs.260132.

111. Madison, J.M., Nurrish, S., and Kaplan, J.M. (2005). UNC-13 interaction with syntaxin is required for synaptic transmission. Curr Biol 15, 2236–2242. 10.1016/j.cub.2005.10.049.

112. Redlingshofer, L., McLeod, F., Chen, Y., Camus, M.D., Burden, J.J., Palomer, E., Briant, K., Dannhauser, P.N., Salinas, P.C., and Brodsky, F.M. (2020). Clathrin light chain diversity regulates membrane deformation in vitro and synaptic vesicle formation in vivo. Proc Natl Acad Sci U S A 117, 23527–23538. 10.1073/pnas.2003662117.

113. Sundborger, A., Soderblom, C., Vorontsova, O., Evergren, E., Hinshaw, J.E., and Shupliakov, O. (2011). An endophilin-dynamin complex promotes budding of clathrin-coated vesicles during synaptic vesicle recycling. J Cell Sci 124, 133–143. 10.1242/jcs.072686.

114. Tagliatti, E., Fadda, M., Falace, A., Benfenati, F., and Fassio, A. (2016). Arf6 regulates the cycling and the readily releasable pool of synaptic vesicles at hippocampal synapse. Elife 5. 10.7554/eLife.10116.

115. Essmann, C.L., Martinez, E., Geiger, J.C., Zimmer, M., Traut, M.H., Stein, V., Klein, R., and Acker-Palmer, A. (2008). Serine phosphorylation of ephrinB2 regulates trafficking of synaptic AMPA receptors. Nat Neurosci 11, 1035–1043. 10.1038/nn.2171.

116. Yang, Y., Geng, Y., Jiang, D., Ning, L., Kim, H.J., Jeon, N.L., Lau, A., Chen, L., and Lin, M.Z. (2019). Kinase pathway inhibition restores PSD95 induction in neurons lacking fragile X mental retardation protein. Proc Natl Acad Sci U S A 116, 12007–12012. 10.1073/pnas.1812056116.

117. Palop, J.J., Chin, J., Roberson, E.D., Wang, J., Thwin, M.T., Bien-Ly, N., Yoo, J., Ho, K.O., Yu, G.Q., Kreitzer, A., et al. (2007). Aberrant excitatory neuronal activity and compensatory remodeling of inhibitory hippocampal circuits in mouse models of Alzheimer’s disease. Neuron 55, 697–711. 10.1016/j.neuron.2007.07.025.

118. Perdigao, C., Barata, M.A., Araujo, M.N., Mirfakhar, F.S., Castanheira, J., and Guimas Almeida, C. (2020). Intracellular Trafficking Mechanisms of Synaptic Dysfunction in Alzheimer’s Disease. Front Cell Neurosci 14, 72. 10.3389/fncel.2020.00072.

119. Scaduto, P., Lauterborn, J.C., Cox, C.D., Fracassi, A., Zeppillo, T., Gutierrez, B.A., Keene, C.D., Crane, P.K., Mukherjee, S., Russell, W.K., et al. (2023). Functional excitatory to inhibitory synaptic imbalance and loss of cognitive performance in people with Alzheimer’s disease neuropathologic change. Acta Neuropathol 145, 303–324. 10.1007/s00401-022-02526-0.

120. Scheff, S.W., and Price, D.A. (2006). Alzheimer’s disease-related alterations in synaptic density: neocortex and hippocampus. J Alzheimers Dis 9, 101–115. 10.3233/jad-2006-9s312.

121. Wu, X.M., Shi, C.N., Liu, K., Hu, X.Y., He, Q.L., Yao, H., Fan, D., Ma, D.Q., Yang, J.J., Shen, J.C., and Ji, M.H. (2025). Decreased excitatory and increased inhibitory transmission in the hippocampal CA1 drive neuroinflammation-induced cognitive impairments in mice. Brain Behav Immun 128, 416–428. 10.1016/j.bbi.2025.04.027.

122. Dechant, G., and Barde, Y.A. (2002). The neurotrophin receptor p75(NTR): novel functions and implications for diseases of the nervous system. Nat Neurosci 5, 1131–1136. 10.1038/nn1102-1131.

123. Manucat-Tan, N.B., Shen, L.L., Bobrovskaya, L., Al-Hawwas, M., Zhou, F.H., Wang, Y.J., and Zhou, X.F. (2019). Knockout of p75 neurotrophin receptor attenuates the hyperphosphorylation of Tau in pR5 mouse model. Aging (Albany NY) 11, 6762–6791. 10.18632/aging.102202.

124. Ristic, G., Tsou, W.L., and Todi, S.V. (2014). An optimal ubiquitin-proteasome pathway in the nervous system: the role of deubiquitinating enzymes. Front Mol Neurosci 7, 72. 10.3389/fnmol.2014.00072.

125. Bishop, P., Rubin, P., Thomson, A.R., Rocca, D., and Henley, J.M. (2014). The ubiquitin C-terminal hydrolase L1 (UCH-L1) C terminus plays a key role in protein stability, but its farnesylation is not required for membrane association in primary neurons. J Biol Chem 289, 36140–36149. 10.1074/jbc.M114.557124.

126. Yu, Q., Zhang, H., Li, Y., Liu, C., Wang, S., and Liao, X. (2018). UCH-L1 Inhibition Suppresses tau Aggresome Formation during Proteasomal Impairment. Mol Neurobiol 55, 3812–3821. 10.1007/s12035-017-0558-7.

127. Zhang, M., Deng, Y., Luo, Y., Zhang, S., Zou, H., Cai, F., Wada, K., and Song, W. (2012). Control of BACE1 degradation and APP processing by ubiquitin carboxyl-terminal hydrolase L1. J Neurochem 120, 1129–1138. 10.1111/j.1471-4159.2011.07644.x.

128. Granger, B.E., and Perez, F. (2021). Jupyter: Thinking and Storytelling With Code and Data. Computing in Science & Engineering 23, 7–14. 10.1109/mcse.2021.3059263.

129. Besser, L., Kukull, W., Knopman, D.S., Chui, H., Galasko, D., Weintraub, S., Jicha, G., Carlsson, C., Burns, J., Quinn, J., et al. (2018). Version 3 of the National Alzheimer’s Coordinating Center’s Uniform Data Set. Alzheimer Dis Assoc Disord 32, 351–358. 10.1097/WAD.0000000000000279.

130. Morris, J.C. (1993). The Clinical Dementia Rating (CDR): current version and scoring rules. Neurology 43, 2412–2414. 10.1212/wnl.43.11.2412-a.

131. Nasreddine, Z.S., Phillips, N.A., Bedirian, V., Charbonneau, S., Whitehead, V., Collin, I., Cummings, J.L., and Chertkow, H. (2005). The Montreal Cognitive Assessment, MoCA: a brief screening tool for mild cognitive impairment. J Am Geriatr Soc 53, 695–699. 10.1111/j.1532-5415.2005.53221.x.

132. Ratcliffe, L.N., McDonald, T., Robinson, B., Sass, J.R., Loring, D.W., and Hewitt, K.C. (2023). Classification statistics of the Montreal Cognitive Assessment (MoCA): Are we interpreting the MoCA correctly? Clin Neuropsychol 37, 562–576. 10.1080/13854046.2022.2086487.

133. Folstein, M.F., Folstein, S.E., and McHugh, P.R. (1975). “Mini-mental state”. A practical method for grading the cognitive state of patients for the clinician. J Psychiatr Res 12, 189–198. 10.1016/0022-3956(75)90026-6.

134. Thal, D.R., Rub, U., Orantes, M., and Braak, H. (2002). Phases of A beta-deposition in the human brain and its relevance for the development of AD. Neurology 58, 1791–1800. 10.1212/wnl.58.12.1791.

135. Braak, H., Alafuzoff, I., Arzberger, T., Kretzschmar, H., and Del Tredici, K. (2006). Staging of Alzheimer disease-associated neurofibrillary pathology using paraffin sections and immunocytochemistry. Acta Neuropathol 112, 389–404. 10.1007/s00401-006-0127-z.

136. Mirra, S.S., Heyman, A., McKeel, D., Sumi, S.M., Crain, B.J., Brownlee, L.M., Vogel, F.S., Hughes, J.P., van Belle, G., and Berg, L. (1991). The Consortium to Establish a Registry for Alzheimer’s Disease (CERAD). Part II. Standardization of the neuropathologic assessment of Alzheimer’s disease. Neurology 41, 479–486. 10.1212/wnl.41.4.479.

137. Perez-Riverol, Y., Bai, J., Bandla, C., Garcia-Seisdedos, D., Hewapathirana, S., Kamatchinathan, S., Kundu, D.J., Prakash, A., Frericks-Zipper, A., Eisenacher, M., et al. (2022). The PRIDE database resources in 2022: a hub for mass spectrometry-based proteomics evidences. Nucleic Acids Res 50, D543–D552. 10.1093/nar/gkab1038.

138. Chen, H., Lundberg, S.M., and Lee, S.-I. (2022). Explaining a series of models by propagating Shapley values. Nature Communications 13, 4512. 10.1038/s41467-022-31384-3.

