## Supplementary data for "Synergistic retinal UCHL1 dysregulation and synaptic vulnerability reflect Alzheimer’s disease severity"

**Table S1**. List of human donors in this study.
**Table S2.** Stratification of retinal synaptic markers by CDR.
**Table S3.** Stratification of retinal synaptic markers by Braak stage.
**Table S4.** Stratification of retinal synaptic markers by MMSE score.
**Table S5.** Correlation between retinal synaptic markers and brain pathology.
**Table S6.** Synapse-associated DEPs upregulated in AD versus NC.
**Table S7.** Synapse-associated DEPs downregulated in AD versus NC.

**Figure S1.** Pre- and post-synaptic density in retinal sub-regions from MCI and AD patients and relations to retinal amyloidosis and tauopathy.
**Figure S2.** Gene set enrichment analyses of AD retina mass spectrometry data.
**Figure S3.** P75NTR and photoreceptor degeneration pathway in the AD retina.
**Figure S4.** Full western blots of PSD95, Synaptophysin and p75NTR.
**Figure S5.** Regional and spatial retinal atrophy assessed by retinal thickness.
**Figure S6.** Extended data on retinal UCHL1.
**Figure S7.** UCHL1 and associated molecular pathways in the AD retina.

**Table S1. List of postmortem human retinas and brains used in this study.**

| Diagnosis | Age at death | Thal A | Braak B | CERAD C | CAA score | Co-morbid (LB/AS/VD) | Braak stage | CDR score | MMSE score | Study type |
|---|---|---|---|---|---|---|---|---|---|---|
| NC1 | 86 | n.a. | n.a. | n.a. | n.a. | n.a. | n.a. | n.a. | n.a. | Hist |
| NC2 | 75 | n.a. | n.a. | n.a. | n.a. | n.a. | n.a. | n.a. | n.a. | Hist |
| NC3 | 76 | n.a. | n.a. | n.a. | n.a. | n.a. | n.a. | n.a. | n.a. | Hist |
| NC4 | 73 | n.a. | n.a. | n.a. | n.a. | n.a. | n.a. | n.a. | n.a. | Hist |
| NC5 | 85 | 2 | 1 | 2 | 0 | −/+ | I-II | 0 | 30 | Hist |
| NC6 | 80 | n.a. | n.a. | n.a. | n.a. | n.a. | n.a. | n.a. | n.a. | Hist |
| NC7 | 84 | n.a. | n.a. | n.a. | n.a. | n.a. | n.a. | n.a. | 30 | Hist |
| NC8 | 81 | 3 | 1 | 2 | 0.25 | −/+ | I-II | 0 | 23 | Hist |
| NC9 | 95 | 3 | 3 | 2 | 1 | −/+ | V | 0 | 30 | Hist |
| NC10 | 69 | 0 | 0 | 1 | 0 | −/+ | 0 | 1 | 28 | Hist |
| NC11 | 88 | 2 | 1 | 2 | 0 | −/+ | I | 0 | 30 | Hist |
| NC12 | 77 | n.a. | n.a. | n.a. | n.a. | n.a. | n.a. | n.a. | 30 | Hist |
| NC13 | 70 | n.a. | n.a. | n.a. | n.a. | n.a. | n.a. | n.a. | 30 | Hist |
| NC14 | 87 | n.a. | n.a. | n.a. | n.a. | n.a. | n.a. | n.a. | 30 | Hist |
| NC15 | 58 | n.a. | n.a. | n.a. | n.a. | n.a. | n.a. | n.a. | 30 | Hist |
| NC16 | 92 | 1 | 1 | 0 | 0.5 | −/+ | I | n.a. | 25 | Hist |
| NC17 | 95 | 1 | 1 | 1 | 0 | n.a. | I | 0 | 30 | Hist |
| NC18 | 76 | 2 | 0 | 2 | 0 | n.a. | 0 | 0 | 29 | Hist |
| NC19 | 95 | 1 | 0 | 0 | 0.5 | −/+ | I-II | 0 | 30 | Hist |
| NC20 | 99 | 1 | 2 | 1 | 0 | n.a. | III | 0 | 29 | Hist |
| NC21 | 93 | 3 | 2 | 3 | 0.5 | −/+ | III-IV | 1 | 27 | Hist |
| MCI1 | 93 | 2 | 0 | 2 | 0 | −/+ | 0 | 3 | 19 | Hist |
| MCI2 | 86 | 3 | 1 | 3 | 0 | −/+ | I-II | 2 | 15 | Hist |
| MCI3 | 94 | 2 | 1 | 2 | 0 | n.a. | I-II | 0.5 | 29 | Hist |
| MCI4 | 88 | 1 | 2 | 2 | 0 | −/+ | III | 3 | n.a. | Hist |
| MCI5 | 93 | 3 | 2 | 2 | 2 | −/+ | IV | 3 | 11 | Hist |
| MCI6 | 98 | 2 | 3 | 2 | 2 | −/+ | V | 2 | 15 | Hist |
| MCI7 | 83 | 2 | 3 | 2 | 1.5 | −/+ | III-IV | 0 | 26 | Hist |
| MCI8 | 91 | 2 | 2 | 2 | 0 | −/+ | III | 3 | 29 | Hist |
| MCI9 | 87 | 3 | 3 | 3 | 1.5 | −/+ | V-VI | 3 | 13 | Hist |
| MCI10 | 89 | 1 | 2 | 2 | 1 | n.a. | III-IV | 0.5 | 24 | Hist |
| MCI11 | 80 | 3 | 3 | 2 | 1 | −/+ | V | 3 | 29 | Hist |
| AD1 | 76 | 3 | 3 | 3 | 2 | −/+ | V | 1 | 26 | Hist |
| AD2 | 90 | 1 | 3 | 3 | 1 | −/+ | V | 3 | 9 | Hist |
| AD3 | 93 | 3 | 2 | 2 | 0 | −/+ | III-IV | 3 | 20 | Hist |

| Diagnosis | Age at death | Thal A | Braak B | CERAD C | CAA score | Co-morbid (LB/AS/VD) | Braak stage | CDR score | MMSE score | Study type |
|---|---|---|---|---|---|---|---|---|---|---|
| AD4 | 86 | 3 | 3 | 2 | 1 | −/+ | V-VI | 3 | 18 | Hist |
| AD5 | 88 | 3 | 3 | 3 | 1.5 | −/+ | V-VI | 1 | 16 | Hist |
| AD6 | 93 | 2 | 2 | 2 | 1.5 | −/+ | III-IV | 3 | 17 | Hist |
| AD7 | 90 | 3 | 2 | 3 | 1 | −/+ | IV | 3 | n.a. | Hist |
| AD8 | 88 | 2 | 3 | 3 | 1.5 | −/+ | V | 3 | 4 | Hist |
| AD9 | 100 | 2 | 3 | 3 | 1 | −/+ | V-VI | 2 | 16 | Hist |
| AD10 | 88 | 2 | 3 | 2 | 1 | −/+ | V-VI | 1 | 4 | Hist |
| AD11 | 90 | 3 | 3 | 3 | 0 | −/+ | V-VI | n.a. | n.a | Hist |
| AD12 | 81 | 3 | 3 | 3 | 2 | −/+ | VI | n.a. | 4 | Hist |
| AD13 | 83 | 3 | 2 | 3 | 3 | −/+ | IV | 1 | 18 | Hist |
| AD14 | 89 | 3 | 3 | 3 | 2 | −/+ | V-VI | n.a. | 6 | Hist |
| AD15 | 75 | 3 | 3 | 3 | 0 | −/+ | III | n.a. | 4 | Hist |
| AD16 | 71 | 3 | 3 | 3 | 2 | −/+ | III | n.a. | 10 | Hist |
| AD17 | 64 | 3 | 3 | 3 | 0 | −/+ | III | 3 | 5 | Hist |
| AD18 | 90 | 3 | 3 | 2 | 2 | −/+ | III | 2 | 18 | Hist |
| AD19 | 97 | 3 | 3 | 3 | 0 | −/+ | III | n.a. | 4 | Hist |
| AD20 | 97 | 3 | 2 | 3 | 1 | −/+ | III | 1 | 26 | Hist |
| AD21 | 81 | 3 | 3 | 3 | 1.5 | −/+ | III | 3 | 12 | Hist |
| AD1 | 48 | 3 | 3 | 3 | 2 | −/− | V-VI | n.a. | n.a. | Proteins |
| AD2 | 93 | 2 | 3 | 3 | 1 | −/+ | V | 3 | n.a. | Proteins |
| AD3 | 88 | 2 | 3 | 2 | 1 | −/+ | V-VI | 1 | 4 | Proteins |
| AD4 | 88 | 1 | 2 | 2 | 0 | −/+ | III | 3 | n.a. | Proteins |
| AD5 | 94 | 3 | 3 | 3 | 0 | −/+ | V-VI | 3 | n.a. | Proteins |
| AD6 | 100 | 2 | 3 | 3 | 0.5 | −/+ | V-VI | 2 | 16 | Proteins |
| AD7 | 81 | 3 | 3 | 3 | 0 | −/+ | V-VI | n.a. | 12 | Proteins |
| AD8 | 78 | n.a. | n.a. | n.a. | n.a. | n.a. | n.a. | n.a. | n.a. | Proteins |
| AD9 | 74 | n.a. | n.a. | n.a. | n.a. | n.a. | n.a. | n.a. | n.a. | Proteins |
| NC1 | 75 | n.a. | n.a. | n.a. | n.a. | n.a. | n.a. | n.a. | n.a. | Proteins |
| NC2 | 72 | n.a. | n.a. | n.a. | n.a. | n.a. | n.a. | n.a. | n.a. | Proteins |
| NC3 | 81 | 3 | 1 | 2 | 0.25 | −/− | I-II | 0 | 23 | Proteins |
| NC4 | 69 | n.a. | n.a. | n.a. | n.a. | n.a. | n.a. | n.a. | n.a. | Proteins |
| NC5 | 79 | n.a. | n.a. | n.a. | n.a. | n.a. | n.a. | n.a. | n.a. | Proteins |
| NC6 | 85 | 0 | 1 | 3 | 1 | −/+ | I-II | 0.5 | n.a. | Proteins |
| NC7 | 76 | n.a. | n.a. | n.a. | n.a. | n.a. | n.a. | n.a. | n.a. | Proteins |

AD, Alzheimer's disease dementia; MCI, mild cognitive impairment; CN, normal cognition; F, female; M, male; A, Asian; B, Black; H, Hispanic; W, White; His, Histology; A, Aβ plaque score modified from Thal; B, NFT stage modified from Braak; C, Neuritic plaque score modified from CERAD; CAA, Cerebral amyloid angiopathy; LB, Lewy bodies; ASVD. Atherosclerosis; CDR, Clinical dementia rating; MMSE, Mini-Mental State Examination; n.a., not available; +: present; –: none; APOE, apolipoprotein alleles.

**Table S2**. Stratification of retinal synaptic markers by CDR score.

| Retinal | CDR=0-0.5 | | | CDR=1-2 | | | CDR=3 | | | CDR 0-0.5 vs 1-2 | | CDR 1-2 vs 3 | | CDR 0-0.5 vs 3 | |
|---|---|---|---|---|---|---|---|---|---|---|---|---|---|---|---|
| | Mean | SD | n | Mean | SD | n | Mean | SD | n | P | FC [95% CI] | P | FC [95% CI] | P | FC [95% CI] |
| VGLUT1 | 36.3 | 9.7 | 9 | 22.2 | 10.3 | 10 | 19.3 | 5.6 | 11 | <0.0001 | -1.6 [-2.18, -1.09] | 0.52 | -1.1 [-1.53, -0.77] | <0.0001 | -1.9 [-2.34, -1.42] |
| SYP | 19.6 | 7.2 | 5 | 9.1 | 4.0 | 5 | 6.4 | 3.0 | 8 | 0.019 | -2.2 [-3.24, -1.07] | 0.71 | -1.4 [-2.15, -0.71] | 0.0006 | -3.1 [-4.48, -1.68] |
| PSD95 | 14.0 | 4.9 | 9 | 9.5 | 5.7 | 12 | 7.3 | 1.8 | 13 | 0.21 | -1.5 [-2.08, -0.87] | 0.63 | -1.3 [-1.79, -0.83] | 0.0304 | -1.9 [-2.44, -1.42] |
| NMDAR2A | 17.6 | 8.1 | 7 | 7.5 | 2.7 | 6 | 7.0 | 2.2 | 11 | 0.0092 | -2.4 [-3.41, -1.29] | 0.99 | -1.1 [-1.44, 10.70] | 0.0013 | -2.5 [-3.50, -1.54] |

P < 0.05 value determines the statistical significance by two-way ANOVA and Tukey's post-hoc multiple comparison test. CDR, Clinical dementia rating; CI, Confidence interval; FC, Fold change; n, sample size; SD, Standard deviation

**Table S3**. Stratification of retinal synaptic markers by Braak stage.

| **Retinal** | **Braak=0-II** | | | **Braak=III-IV** | | | **Braak=V-VI** | | | **Braak 0-II vs III-IV** | | **Braak III-IV vs V-VI** | | **Braak 0-II vs V-VI** | |
|---|---|---|---|---|---|---|---|---|---|---|---|---|---|---|---|
| | Mean | SD | n | Mean | SD | n | Mean | SD | n | P | FC [95% CI] | P | FC [95% CI] | P | FC [95% CI] |
| VGLUT1 | 34.8 | 10.1 | 8 | 23.0 | 10.9 | 11 | 20.6 | 7.8 | 13 | 0.0006 | -1.5 [-2.04, -0.99] | 0.67 | -1.1 [-1.50, -0.73] | <0.0001 | -1.7 [-2.17, -1.20] |
| SYP | 17.2 | 5.8 | 5 | 6.3 | 3.1 | 4 | 8.6 | 6.8 | 11 | 0.042 | -2.7 [-4.26, -1.19] | 0.82 | -0.7 [-1.22, -0.24] | 0.049 | -2.0 [-3.09, -0.90] |
| PSD95 | 14.7 | 5.1 | 8 | 9.1 | 5.1 | 12 | 7.0 | 3.0 | 19 | 0.16 | -1.6 [-2.25, -0.97] | 0.66 | -1.3 [-1.78, -0.82] | 0.019 | -2.1 [-2.74, -1.45] |
| NMDAR2A | 15.8 | 9.4 | 7 | 8.0 | 2.4 | 7 | 7.2 | 4.0 | 12 | 0.074 | -2.0 [-2.96, -1.00] | 0.97 | -1.1 [-1.54, -0.68] | 0.02 | -2.2 [-3.39. -1.01] |

P < 0.05 value determines the statistical significance by two-way ANOVA and Tukey's post-hoc multiple comparison test. CI, Confidence interval; FC, Fold change; n, sample size; SD, Standard deviation.

**Table S4**. Stratification of retinal synaptic markers by MMSE score.

| **Retinal** | **MMSE>26** | | | **MMSE≤26** | | | **2way ANOVA Multiple comparisons** | |
|---|---|---|---|---|---|---|---|---|
| | Mean | SD | n | Mean | SD | n | P value | FC [95% CI] |
| VGLUT1 | 41.8 | 7.1 | 14 | 20.0 | 6.1 | 22 | <0.0001 | -2.1 [-2.42, -1.77] |
| SYP | 24.8 | 8.0 | 9 | 8.2 | 5.0 | 14 | <0.0001 | -3.0 [-4.16, -1.87] |
| PSD95 | 16.2 | 4.5 | 16 | 7.0 | 2.6 | 26 | <0.0001 | -2.3 [-2.76, -1.85] |
| NMDAR2A | 19.2 | 6.9 | 12 | 7.6 | 3.4 | 18 | <0.0001 | -2.5 [-3.27, -1.79] |

P < 0.05 value determines the statistical significance by two-way ANOVA with Šídák's post-hoc multiple comparison test. CI, Confidence interval; FC, Fold change; MMSE, Mini-Mental State Examination, n, sample size; SD, Standard deviation.

**Table S5**. Correlation between retinal synaptic markers and brain pathology.

| Brain / Retina | **Aβ plaques (severity score)** | **NFTs (severity score)** |
|---|---|---|
| ***VGLUT1*** | −0.071<br>32 | −0.38<br>32 |
| ***SYP*** | −0.47*<br>20 | −0.65**<br>20 |
| ***PSD95*** | −0.34*<br>39 | −0.45**<br>39 |
| ***NMDAR2A*** | −0.42*<br>26 | −0.48*<br>26 |

Pairwise Spearman's rank coefficient correlation analyses showing the strength of the association (*r*, upper value), sample size number (n, lower value), and adjusted P values with Holm-Šídák multiple-comparison correction method (adjusted for each retinal synaptic marker against 2 brain pathology scores): *P < 0.05, **P < 0.01.

**Table S6**. Synapse-associated DEPs upregulated in AD versus NC.

| Accession | Symbol | Description | NC | AD | FC | *P* | adj. *P* |
|---|---|---|---|---|---|---|---|
| O75487 | GPC4 | Glypican-4 | 20.9 | 39.1 | 1.87 | 0.0178 | 0.1898 |
| P09497 | CLTB | Clathrin light chain B | 249.1 | 462.2 | 1.86 | 0.0036 | 0.1294 |
| P62942 | FKBP1A | Peptidyl-prolyl cis-trans isomerase FKBP1A | 120.3 | 209.9 | 1.75 | 0.0047 | 0.1826 |
| Q01469 | FABP5 | Fatty acid-binding protein 5 | 1176.3 | 2053.5 | 1.75 | 0.0126 | 0.1736 |
| Q13542 | EIF4EBP2 | Eukaryotic translation initiation factor 4E-binding protein 2 | 11.3 | 18.9 | 1.67 | 0.0165 | 0.1857 |
| P08138 | NGFR | Tumor necrosis factor receptor superfamily member 16 | 24.7 | 40.4 | 1.64 | 0.0007 | 0.1203 |
| Q6PUV4 | CPLX2 | Complexin-2 | 131.2 | 214.0 | 1.63 | 0.0177 | 0.1894 |
| Q6UUV9 | CRTC1 | CREB-regulated transcription coactivator 1 | 54.5 | 83.5 | 1.53 | 0.0151 | 0.1826 |
| P42574 | CASP3 | Caspase-3 | 24.0 | 35.4 | 1.47 | 0.0073 | 0.1496 |
| Q13303 | KCNAB2 | Voltage-gated potassium channel subunit beta-2 | 83.9 | 122.1 | 1.46 | 0.0010 | 0.1203 |
| P62330 | ARF6 | ADP-ribosylation factor 6 | 298.8 | 431.8 | 1.45 | 0.0031 | 0.1285 |
| Q13526 | PIN1 | Peptidyl-prolyl cis-trans isomerase NIMA-interacting 1 | 719.4 | 1031.8 | 1.43 | 0.0106 | 0.1670 |
| I3L504 | EIF5A | Eukaryotic translation initiation factor 5A-1 | 956.2 | 1338.9 | 1.40 | 0.0211 | 0.1951 |
| P18085 | ARF4 | ADP-ribosylation factor 4 | 645.8 | 883.5 | 1.37 | 0.0007 | 0.1203 |
| P62166 | NCS1 | Neuronal calcium sensor 1 | 91.9 | 124.2 | 1.35 | 0.0090 | 0.1594 |
| P23763 | VAMP1 | Vesicle-associated membrane protein 1 | 117.0 | 157.1 | 1.34 | 0.0172 | 0.1880 |
| P46934 | NEDD4 | E3 ubiquitin-protein ligase NEDD4 | 14.4 | 19.3 | 1.34 | 0.0189 | 0.1939 |
| P68032 | ACTC1 | Actin, alpha cardiac muscle 1 | 3608.7 | 4806.4 | 1.33 | 0.0138 | 0.1775 |
| Q14643 | ITPR1 | Inositol 1,4,5-trisphosphate receptor type 1 | 10.7 | 14.2 | 1.33 | 0.0206 | 0.1944 |
| H7C394 | CAMK2B | Calcium/calmodulin-dependent protein kinase type II subunit beta | 41.1 | 54.4 | 1.32 | 0.0060 | 0.1415 |
| Q14012 | CAMK1 | Calcium/calmodulin-dependent protein kinase type 1 | 108.6 | 143.4 | 1.32 | 0.0003 | 0.1203 |
| A0A0U1RRM6 | ENAH | Protein enabled homolog | 235.1 | 310.3 | 1.32 | 0.0044 | 0.1349 |
| Q14515 | SPARCL1 | SPARC-like protein 1 | 420.5 | 552.9 | 1.31 | 0.0119 | 0.1702 |
| E5RJR5 | SKP1 | S-phase kinase-associated protein 1 | 1575.9 | 2046.6 | 1.30 | 0.0131 | 0.1744 |
| O00499 | BIN1 | Myc box-dependent-interacting protein 1 | 61.5 | 79.7 | 1.30 | 0.0056 | 0.1415 |
| Q8WUY3 | PRUNE2 | Protein prune homolog 2 | 153.4 | 194.3 | 1.27 | 0.0153 | 0.1826 |
| Q9UH65 | SWAP70 | Switch-associated protein 70 | 348.4 | 440.6 | 1.26 | 0.0015 | 0.1203 |
| O43765 | SGTA | Small glutamine-rich tetratricopeptide repeat-containing protein alpha | 1074.9 | 1357.5 | 1.26 | 0.0191 | 0.1939 |
| Q9UK22 | FBXO2 | F-box only protein 2 | 778.1 | 982.0 | 1.26 | 0.0030 | 0.1277 |
| Q99961 | SH3GL1 | Endophilin-A2 | 453.7 | 565.3 | 1.25 | 0.0223 | 0.1966 |

| | | | | | | | |
|---|---|---|---|---|---|---|---|
| Q96P47 | AGAP3 | Arf-GAP with GTPase, ANK repeat and PH domain-containing protein 3 | 144.6 | 179.4 | 1.24 | 0.0097 | 0.1653 |
| Q16515 | ASIC2 | Acid-sensing ion channel 2 | 30.3 | 37.6 | 1.24 | 0.0172 | 0.1880 |
| P45974 | USP5 | Ubiquitin carboxyl-terminal hydrolase 5 | 2076.7 | 2537.6 | 1.22 | 0.0093 | 0.1615 |
| P40123 | CAP2 | Adenylyl cyclase-associated protein 2 | 692.6 | 842.2 | 1.22 | 0.0115 | 0.1699 |
| Q9H1P3 | OSBPL2 | Oxysterol-binding protein-related protein 2 | 393.0 | 475.6 | 1.21 | 0.0016 | 0.1203 |
| O15145 | ARPC3 | Actin-related protein 2/3 complex subunit 3 | 276.8 | 334.1 | 1.21 | 0.0216 | 0.1963 |
| P31749 | AKT1 | RAC-alpha serine/threonine-protein kinase | 420.3 | 506.9 | 1.21 | 0.0045 | 0.1349 |

NC (n = 6) and AD (n = 6) are expressed as geometric means.
FC > 1.2; FDR-adjusted (adj.) *P* < 0.2.
The list of synapse-associated proteins was extracted from SynGo release 1.2 knowledge base for synapse research[1].

**Table S7**. Synapse-associated DEPs downregulated in AD versus NC.

| Accession | Symbol | Description | NC | AD | FC | *P* | adj. *P* |
|---|---|---|---|---|---|---|---|
| Q12791 | KCNMA1 | Calcium-activated potassium channel subunit alpha-1 | 125.6 | 69.8 | -1.80 | 0.0187 | 0.1933 |
| A0A286YEX2 | SYT6 | Synaptotagmin VI | 44.4 | 28.6 | -1.55 | 0.0088 | 0.1594 |
| Q13557 | CAMK2D | Calcium/calmodulin-dependent protein kinase type II subunit delta | 116.0 | 75.3 | -1.54 | 0.0090 | 0.1596 |
| O14775 | GNB5 | Guanine nucleotide-binding protein subunit beta-5 | 1751.3 | 1144.8 | -1.53 | 0.0012 | 0.1203 |
| Q9NQC3 | RTN4 | Reticulon-4 | 430.9 | 290.9 | -1.48 | 0.0076 | 0.1534 |
| Q08828 | ADCY1 | Adenylate cyclase type 1 | 582.3 | 394.8 | -1.47 | 0.0004 | 0.1203 |
| Q9UI40 | SLC24A2 | Sodium/potassium/calcium exchanger 2 | 376.1 | 255.8 | -1.47 | 0.0098 | 0.1654 |
| P49207 | RPL34 | 60S ribosomal protein L34 | 1669.6 | 1135.7 | -1.47 | 0.0001 | 0.1203 |
| P83731 | RPL24 | 60S ribosomal protein L24 | 2353.3 | 1601.0 | -1.47 | 0.0019 | 0.1209 |
| P62913 | RPL11 | 60S ribosomal protein L11 | 1643.0 | 1122.0 | -1.46 | 0.0032 | 0.1285 |
| P61353 | RPL27 | 60S ribosomal protein L27 | 1992.6 | 1376.6 | -1.45 | 0.0007 | 0.1203 |
| P62280 | RPS11 | 40S ribosomal protein S11 | 2800.1 | 1944.2 | -1.44 | 0.0063 | 0.1427 |
| P35268 | RPL22 | 60S ribosomal protein L22 | 614.0 | 429.6 | -1.43 | 0.0002 | 0.1203 |
| Q00839 | HNRNPU | Heterogeneous nuclear ribonucleoprotein U | 4906.0 | 3442.1 | -1.43 | 0.0057 | 0.1415 |
| F8VVT9 | AGAP2 | Arf-GAP with GTPase, ANK repeat and PH domain-containing protein 2 | 24.5 | 17.5 | -1.40 | 0.0116 | 0.1700 |
| P61764 | STXBP1 | Syntaxin-binding protein 1 | 1670.3 | 1209.7 | -1.38 | 0.0014 | 0.1203 |
| P52272 | HNRNPM | Heterogeneous nuclear ribonucleoprotein M | 17928.5 | 13023.5 | -1.38 | 0.0041 | 0.1323 |
| P36578 | RPL4 | 60S ribosomal protein L4 | 4413.8 | 3224.1 | -1.37 | 0.0038 | 0.1297 |
| P42766 | RPL35 | 60S ribosomal protein L35 | 2123.7 | 1558.9 | -1.36 | 0.0002 | 0.1203 |
| Q14839 | CHD4 | Chromodomain-helicase-DNA-binding protein 4 | 4536.8 | 3338.6 | -1.36 | 0.0029 | 0.1265 |
| P62244 | RPS15A | 40S ribosomal protein S15a | 1082.2 | 801.5 | -1.35 | 0.0071 | 0.1485 |
| Q14721 | KCNB1 | Potassium voltage-gated channel subfamily B member 1 | 763.1 | 568.7 | -1.34 | 0.0010 | 0.1203 |
| A0A0A0MRM8 | MYO6 | Unconventional myosin-VI | 1382.2 | 1038.1 | -1.33 | 0.0060 | 0.1415 |
| Q9BUJ2 | HNRNPUL1 | Heterogeneous nuclear ribonucleoprotein U-like protein 1 | 3273.8 | 2473.3 | -1.32 | 0.0022 | 0.1209 |
| P42677 | RPS27 | 40S ribosomal protein S27 | 683.1 | 516.6 | -1.32 | 0.0061 | 0.1415 |
| P27635 | RPL10 | 60S ribosomal protein L10 | 1574.7 | 1193.9 | -1.32 | 0.0049 | 0.1399 |
| P62277 | RPS13 | 40S ribosomal protein S13 | 2212.4 | 1696.7 | -1.30 | 0.0037 | 0.1294 |
| P62424 | RPL7A | 60S ribosomal protein L7a | 2942.2 | 2269.0 | -1.30 | 0.0050 | 0.1403 |
| P61026 | RAB10 | Ras-related protein Rab-10 | 2012.1 | 1555.6 | -1.29 | 0.0048 | 0.1385 |
| Q8TCG5 | CPT1C | Carnitine O-palmitoyltransferase 1, brain isoform | 488.7 | 378.6 | -1.29 | 0.0019 | 0.1209 |
| P50914 | RPL14 | 60S ribosomal protein L14 | 1194.2 | 925.5 | -1.29 | 0.0150 | 0.1826 |

| | | | | | | | |
|---|---|---|---|---|---|---|---|
| P40429 | RPL13A | 60S ribosomal protein L13a | 2466.8 | 1913.8 | -1.29 | 0.0032 | 0.1285 |
| Q4R9M9 | KIF1B | Kinesin family member 1Bbeta isoform II | 982.7 | 764.0 | -1.29 | 0.0021 | 0.1209 |
| Q13151 | HNRNPA0 | Heterogeneous nuclear ribonucleoprotein A0 | 3023.5 | 2351.1 | -1.29 | 0.0067 | 0.1448 |
| Q8NBU5 | ATAD1 | ATPase family AAA domain-containing protein 1 | 1262.0 | 986.9 | -1.28 | 0.0194 | 0.1939 |
| P39023 | RPL3 | 60S ribosomal protein L3 | 4503.3 | 3527.3 | -1.28 | 0.0220 | 0.1966 |
| Q15717 | ELAVL1 | ELAV-like protein 1 | 2854.3 | 2237.0 | -1.28 | 0.0025 | 0.1248 |
| P61247 | RPS3A | 40S ribosomal protein S3a | 4305.6 | 3383.3 | -1.27 | 0.0012 | 0.1203 |
| Q9NVJ2 | ARL8B | ADP-ribosylation factor-like protein 8B | 1105.3 | 870.2 | -1.27 | 0.0033 | 0.1287 |
| O60506 | SYNCRIP | Heterogeneous nuclear ribonucleoprotein Q | 33.5 | 26.4 | -1.27 | 0.0120 | 0.1702 |
| P61313 | RPL15 | 60S ribosomal protein L15 | 1127.9 | 890.0 | -1.27 | 0.0043 | 0.1346 |
| P62917 | RPL8 | 60S ribosomal protein L8 | 2556.6 | 2031.3 | -1.26 | 0.0174 | 0.1880 |
| P15880 | RPS2 | 40S ribosomal protein S2 | 3731.9 | 2971.8 | -1.26 | 0.0130 | 0.1742 |
| Q1KMD3 | HNRNPUL2 | Heterogeneous nuclear ribonucleoprotein U-like protein 2 | 5206.0 | 4147.4 | -1.26 | 0.0048 | 0.1384 |
| Q8TAC9 | SCAMP5 | Secretory carrier-associated membrane protein 5 | 2392.0 | 1915.4 | -1.25 | 0.0156 | 0.1829 |
| P29323 | EPHB2 | Ephrin type-B receptor 2 | 954.8 | 764.6 | -1.25 | 0.0127 | 0.1736 |
| Q06787 | FMR1 | Synaptic functional regulator FMR1 | 1013.6 | 811.9 | -1.25 | 0.0045 | 0.1349 |
| Q9BTU6 | PI4K2A | Phosphatidylinositol 4-kinase type 2-alpha | 510.9 | 410.7 | -1.24 | 0.0132 | 0.1744 |
| A0A087WWK8 | IQSEC1 | IQ motif and SEC7 domain-containing protein 1 | 2184.4 | 1761.3 | -1.24 | 0.0195 | 0.1939 |
| P46776 | RPL27A | 60S ribosomal protein L27a | 1539.6 | 1243.1 | -1.24 | 0.0225 | 0.1976 |
| P62241 | RPS8 | 40S ribosomal protein S8 | 2133.6 | 1725.3 | -1.24 | 0.0052 | 0.1403 |
| O75396 | SEC22B | Vesicle-trafficking protein SEC22b | 3207.5 | 2595.5 | -1.24 | 0.0062 | 0.1423 |
| Q9NP72 | RAB18 | Ras-related protein Rab-18 | 3017.4 | 2442.1 | -1.24 | 0.0088 | 0.1594 |
| P26373 | RPL13 | 60S ribosomal protein L13 | 2465.5 | 2000.2 | -1.23 | 0.0147 | 0.1805 |
| O00592 | PODXL | Podocalyxin | 464.2 | 376.8 | -1.23 | 0.0072 | 0.1485 |
| P62701 | RPS4X | 40S ribosomal protein S4, X isoform | 3977.3 | 3228.8 | -1.23 | 0.0102 | 0.1655 |
| Q9UQM7 | CAMK2A | Calcium/calmodulin-dependent protein kinase type II subunit alpha | 925.4 | 752.0 | -1.23 | 0.0109 | 0.1672 |
| P18124 | RPL7 | 60S ribosomal protein L7 | 4105.9 | 3336.5 | -1.23 | 0.0223 | 0.1966 |
| P62888 | RPL30 | 60S ribosomal protein L30 | 810.1 | 662.2 | -1.22 | 0.0165 | 0.1857 |
| Q02878 | RPL6 | 60S ribosomal protein L6 | 3584.9 | 2934.1 | -1.22 | 0.0103 | 0.1656 |
| J3QQ67 | RPL18 | 60S ribosomal protein L18 (Fragment) | 1882.2 | 1543.3 | -1.22 | 0.0224 | 0.1969 |
| Q8NC96 | NECAP1 | Adaptin ear-binding coat-associated protein 1 | 1072.7 | 879.9 | -1.22 | 0.0019 | 0.1209 |
| P23246 | SFPQ | Splicing factor, proline- and glutamine-rich | 9991.3 | 8196.4 | -1.22 | 0.0034 | 0.1290 |
| Q96CW1 | AP2M1 | AP-2 complex subunit mu | 4427.6 | 3634.0 | -1.22 | 0.0227 | 0.1984 |
| F8W059 | UNC13A | Protein unc-13 homolog A | 48.6 | 39.9 | -1.22 | 0.0189 | 0.1939 |
| M0R0P8 | MYO9B | Unconventional myosin-IXb | 42.7 | 35.1 | -1.22 | 0.0013 | 0.1203 |

| | | | | | | | |
|---|---|---|---|---|---|---|---|
| P48454 | PPP3CC | Serine/threonine-protein phosphatase 2B catalytic subunit gamma | 431.4 | 356.4 | -1.21 | 0.0111 | 0.1678 |
| P84098 | RPL19 | 60S ribosomal protein L19 | 751.9 | 622.9 | -1.21 | 0.0206 | 0.1944 |
| P07948 | LYN | Tyrosine-protein kinase Lyn | 2270.9 | 1882.2 | -1.21 | 0.0012 | 0.1203 |
| P62829 | RPL23 | 60S ribosomal protein L23 | 2161.3 | 1798.7 | -1.20 | 0.0081 | 0.1547 |
| P26232 | CTNNA2 | Catenin alpha-2 | 7712.4 | 6434.0 | -1.20 | 0.0066 | 0.1447 |

NC (n = 6) and AD (n = 6) are expressed as geometric means.
|FC| > 1.2; FDR-adjusted (adj.) $P < 0.2$.
The list of synapse-associated proteins was extracted from SynGo release 1.2 knowledge base for synapse research[1].

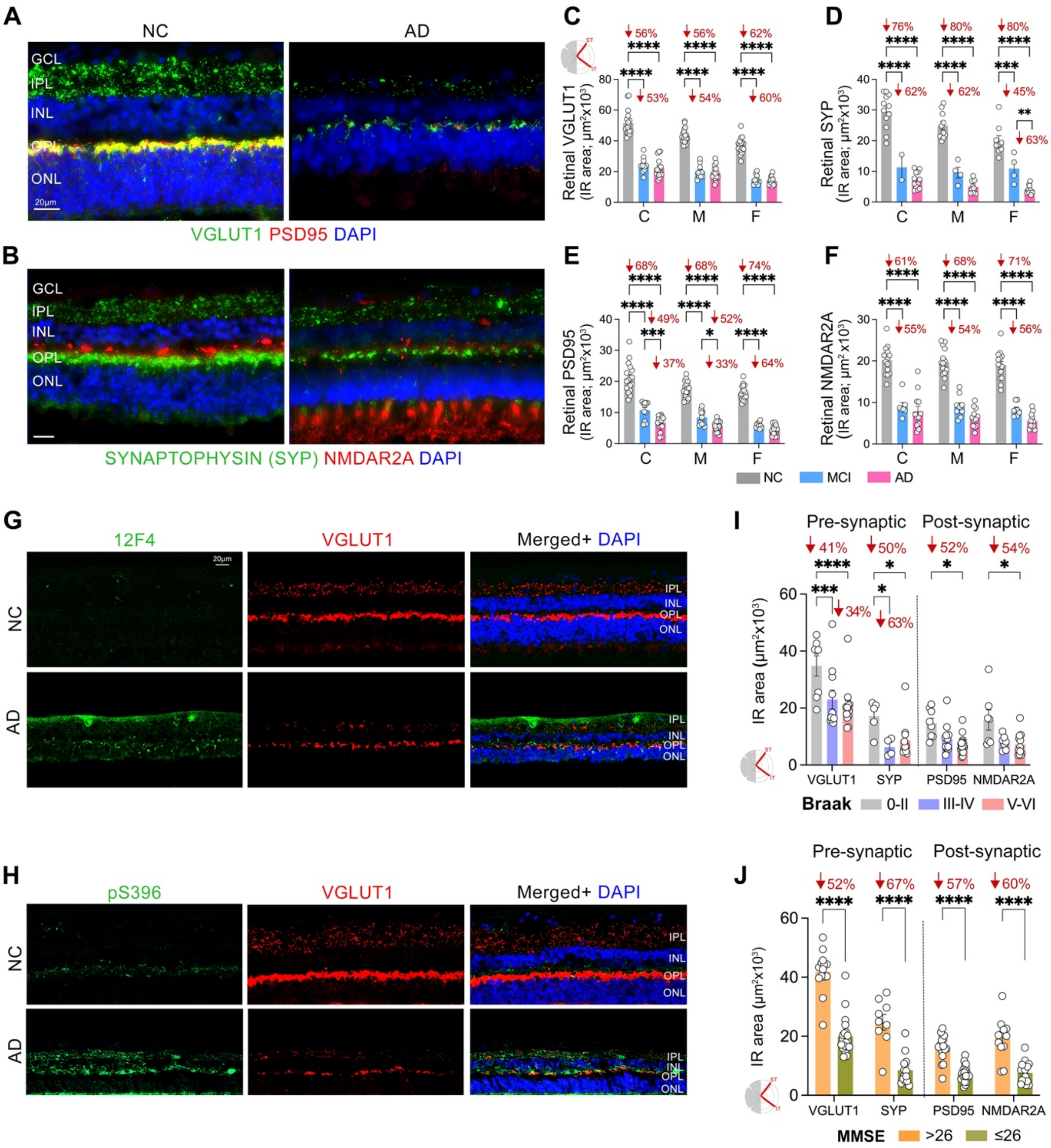
A
NC
AD
GCL
IPL
INL
OPL
ONL
VGLUT1 PSD95 DAPI
B
SYNAPTOPHYSIN (SYP) NMDAR2A DAPI
C
Retinal VGLUT1 (IR area; µm²x10³)
D
Retinal SYP (IR area; µm²x10³)
E
Retinal PSD95 (IR area; µm²x10³)
F
Retinal NMDAR2A (IR area; µm²x10³)
NC MCI AD
G
12F4
VGLUT1
Merged+ DAPI
H
pS396
I
Pre-synaptic
Post-synaptic
IR area (µm²x10³)
VGLUT1 SYP PSD95 NMDAR2A
Braak 0-II III-IV V-VI
J
MMSE >26 ≤26

**Figure S1. Pre- and post-synaptic density in retinal sub-regions from MCI and AD patients and relations to retinal amyloidosis and tauopathy.**

(**A**) Representative high-magnification fluorescence micrographs (x63 objective) of retinal cross-sections immunolabeled for pre-synaptic VGLUT1 (green) and post-synaptic PSD95 (red) markers with nuclei DAPI (blue) in a patient with AD versus NC control. Scale bars: 20 µm.

(**B**) Representative high-magnification fluorescence micrographs (x63 objective) of retinal cross-sections immunolabeled for pre-synaptic Synaptophysin (green) and post-synaptic NMDAR2A (red) in a patient with AD versus NC control. Scale bars: 20 µm.

(**C-F**) Quantitative analyses of VGLUT1 (**C**), SYP (**D**), PSD95 (**E**), and NMDAR2A (**F**) immunoreactive area (IR) in predefined retinal subregions, central (C), mid- (M), and far-periphery (F), in patients with MCI (due to AD; n = 2-11) and AD-dementia (n = 11-20) as compared with age and sex-matched NC controls (n = 9-18).

(**G**) Representative fluorescence micrographs of retinal cross-sections immunolabeled for $12F4^{+}$-$A\beta_{42}$ (green) and VGLUT1 (red) in a patient with AD and in NC control. Nuclei are stained with DAPI (blue). Scale bars: 20 µm.

(**H**) Representative fluorescence micrographs of retinal cross-sections immunolabeled for pS396-tau (green) and VGLUT1 (red) in a patient with AD versus NC control. Scale bars: 20 µm.

(**I, J**). Quantitative analysis of retinal VGLUT1, SYP, PSD95 and NMDAR2A, when donors are stratified according to Braak stage [0-II = 5-8; III-IV = 4-12; V-VI = 11-19] (**I**) and 26-score cutoff on the mini-mental state examination [MMSE; >26 = 9-16; ≤26 = 14-66] (**J**).

Data from individual subjects (circles) as well as group means ± SEMs are shown. *P < 0.05, **P < 0.01, ***P < 0.001, ****P < 0.0001, by two-way ANOVA followed by Tukey's or Šídák's post-hoc multiple comparison tests. Percentage (%) changes are shown in red. Nerve fiber layer (NFL), ganglion cell layer (GCL), inner plexiform layer (IPL), inner nuclear layer (INL), outer plexiform layer (OPL), and outer nuclear layer (ONL).

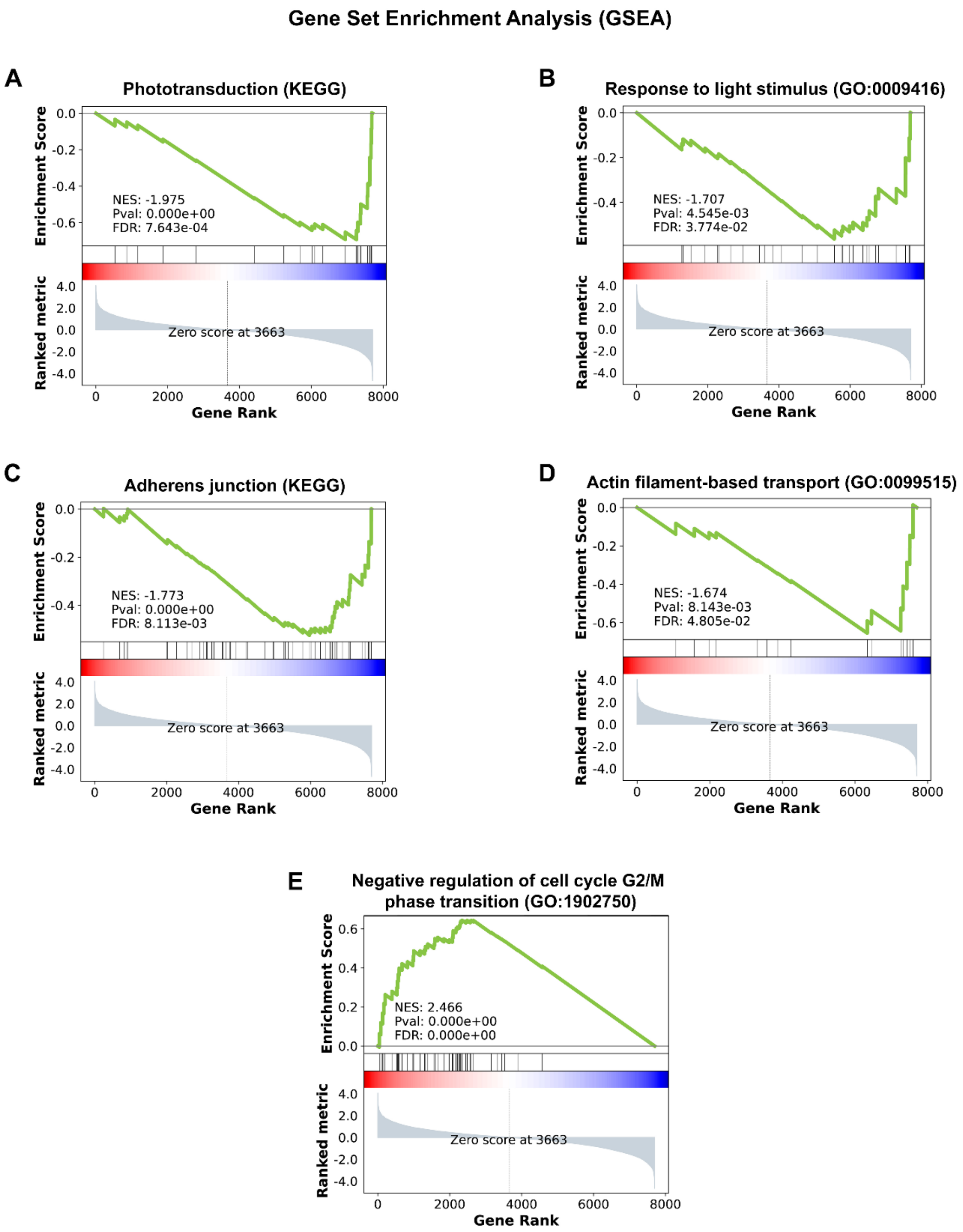


**Figure S2. Gene set enrichment analyses (GSEA) of AD versus NC retina mass spectrometry data**

(**A-B**) GSEA enrichment plots showing enrichment in the bottom-ranked dataset (downregulated in AD) for KEGG phototransduction (NES=-1.975, FDR<0.001) (**A**) and GO response to light stimulus (NES=-1.707, FDR<0.05) (**B**).

(**C-D**) GSEA enrichment plots showing enrichment in the bottom-ranked dataset (downregulated in AD) for KEGG adherens junction (NES=-1.773, FDR<0.01) (**C**) and GO actin filament-based transport (NES=-1.674, FDR<0.05) (**D**).

(**E**) GSEA enrichment plot for GO negative regulation of cell cycle G2/M phase transition enriched in our top-ranked dataset (upregulated in AD: NES=2=466, FDR<0.0001).

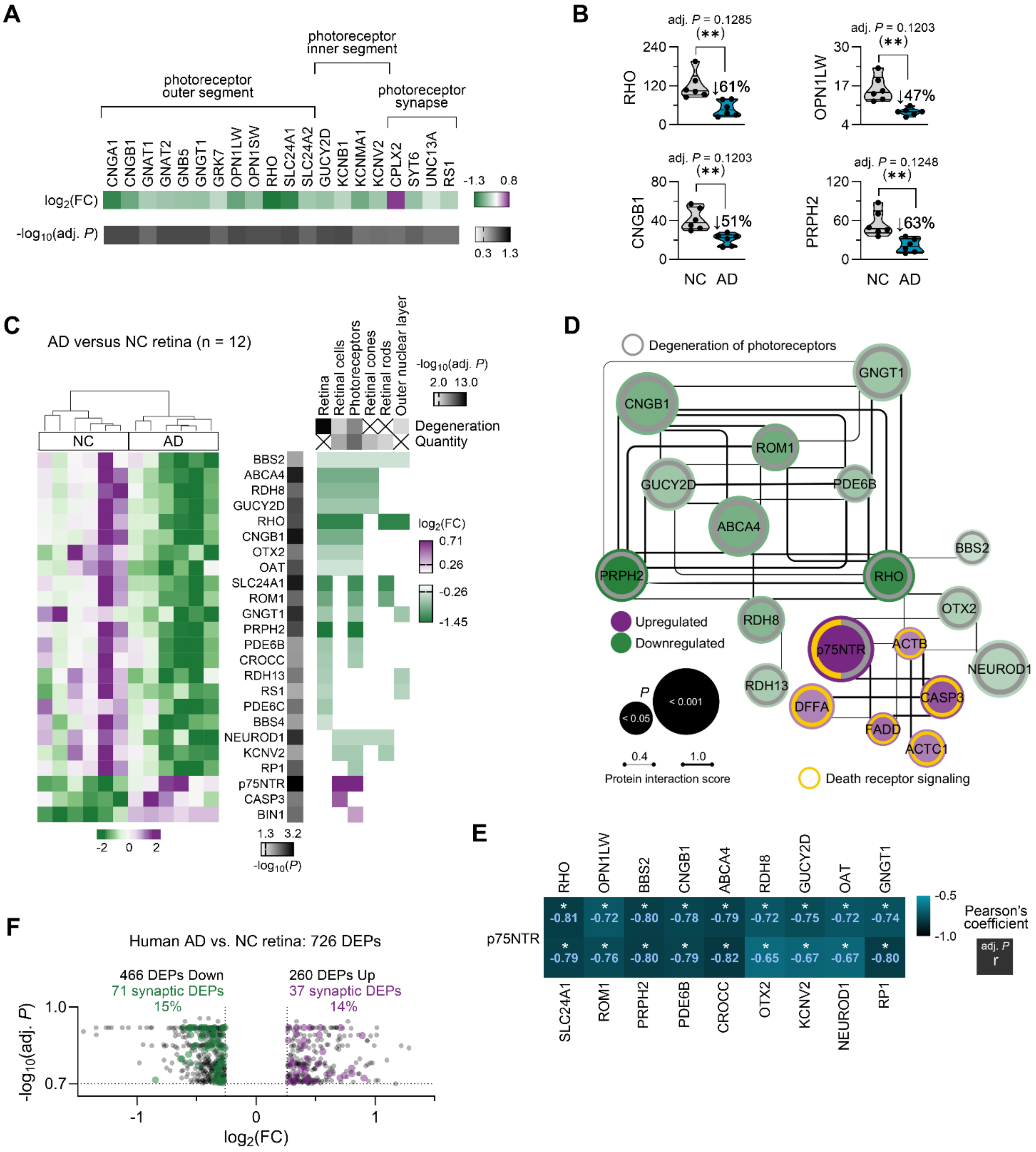
A
photoreceptor outer segment
photoreceptor inner segment
photoreceptor synapse
CNGA1 CNGB1 GNAT1 GNAT2 GNB5 GNGT1 GRK7 OPN1LW OPN1SW RHO SLC24A1 SLC24A2 GUCY2D KCNB1 KCNMA1 KCNV2 CPLX2 SYT6 UNC13A RS1
log2(FC)
-log10(adj. P)
-1.3 0.8
0.3 1.3
B
adj. P = 0.1285
adj. P = 0.1203
adj. P = 0.1203
adj. P = 0.1248
RHO
OPN1LW
CNGB1
PRPH2
↓61%
↓47%
↓51%
↓63%
NC AD
C
AD versus NC retina (n = 12)
NC
AD
Retina
Retinal cells
Photoreceptors
Retinal cones
Retinal rods
Outer nuclear layer
-log10(adj. P)
2.0 13.0
Degeneration
Quantity
BBS2 ABCA4 RDH8 GUCY2D RHO CNGB1 OTX2 OAT SLC24A1 ROM1 GNGT1 PRPH2 PDE6B CROCC RDH13 RS1 PDE6C BBS4 NEUROD1 KCNV2 RP1 p75NTR CASP3 BIN1
log2(FC)
0.71
0.26
-0.26
-1.45
-2 0 2
1.3 3.2
-log10(P)
D
Degeneration of photoreceptors
GNGT1 CNGB1 ROM1 GUCY2D PDE6B ABCA4 BBS2 PRPH2 RHO OTX2 RDH8 p75NTR ACTB NEUROD1 RDH13 DFFA CASP3 FADD ACTC1
Upregulated
Downregulated
P
< 0.05
< 0.001
0.4
1.0
Protein interaction score
Death receptor signaling
E
RHO OPN1LW BBS2 CNGB1 ABCA4 RDH8 GUCY2D OAT GNGT1
p75NTR
-0.81 -0.72 -0.80 -0.78 -0.79 -0.72 -0.75 -0.72 -0.74
-0.79 -0.76 -0.80 -0.79 -0.82 -0.65 -0.67 -0.67 -0.80
SLC24A1 ROM1 PRPH2 PDE6B CROCC OTX2 KCNV2 NEUROD1 RP1
-0.5
-1.0
Pearson's coefficient
adj. P
r
F
Human AD vs. NC retina: 726 DEPs
466 DEPs Down
71 synaptic DEPs
15%
260 DEPs Up
37 synaptic DEPs
14%
-log10(adj. P)
log2(FC)

**Figure S3. P75NTR and photoreceptor degeneration pathway in the AD retina.**
(**A**) Heatmaps of photoreceptor-relevant DEPs in the AD retina, organized by cellular compartment.

(**B**) Violin plots of selected photoreceptor-specific proteins quantified by MS. Individual patient values are presented.

(**C**) Heatmaps of DEPs [$\log_2$(FC) and $-\log_{10}(P)$] in AD retina for select pathways linked to retinal degeneration. The heatmap on the left corresponds to the protein expression level in the 6 NC individuals and the 6 AD patients, standardized by unit variance scaling and generated in ClustVis. Clustering of DEPs was carried out manually based on their involvement in select pathways for visual clarity.

(**D**) Interaction network analysis (String v12.0) of DEPs in AD retina related to photoreceptor degeneration (yellow inner ring) and death receptor signaling (grey inner ring). Blue and red node fills indicate downregulated and upregulated DEPs in AD retina, respectively. The node size increases with the significance level of the difference in protein expression between AD patients and NC individuals.

(**E**) Heatmap of Pearson's correlation coefficients between p75NTR and DEPs in AD retina related photoreceptor degeneration. *adj. $P<0.5$.

(**F**) Volcano plot showing the proportion of dysregulated synapse-associated proteins in AD versus NC control retinas. Synapse-associated proteins were extracted from SynGo v1.2 database.
For this figure, DEPs were defined by $|FC| > 1.2$ and adj. $P < 0.2$.

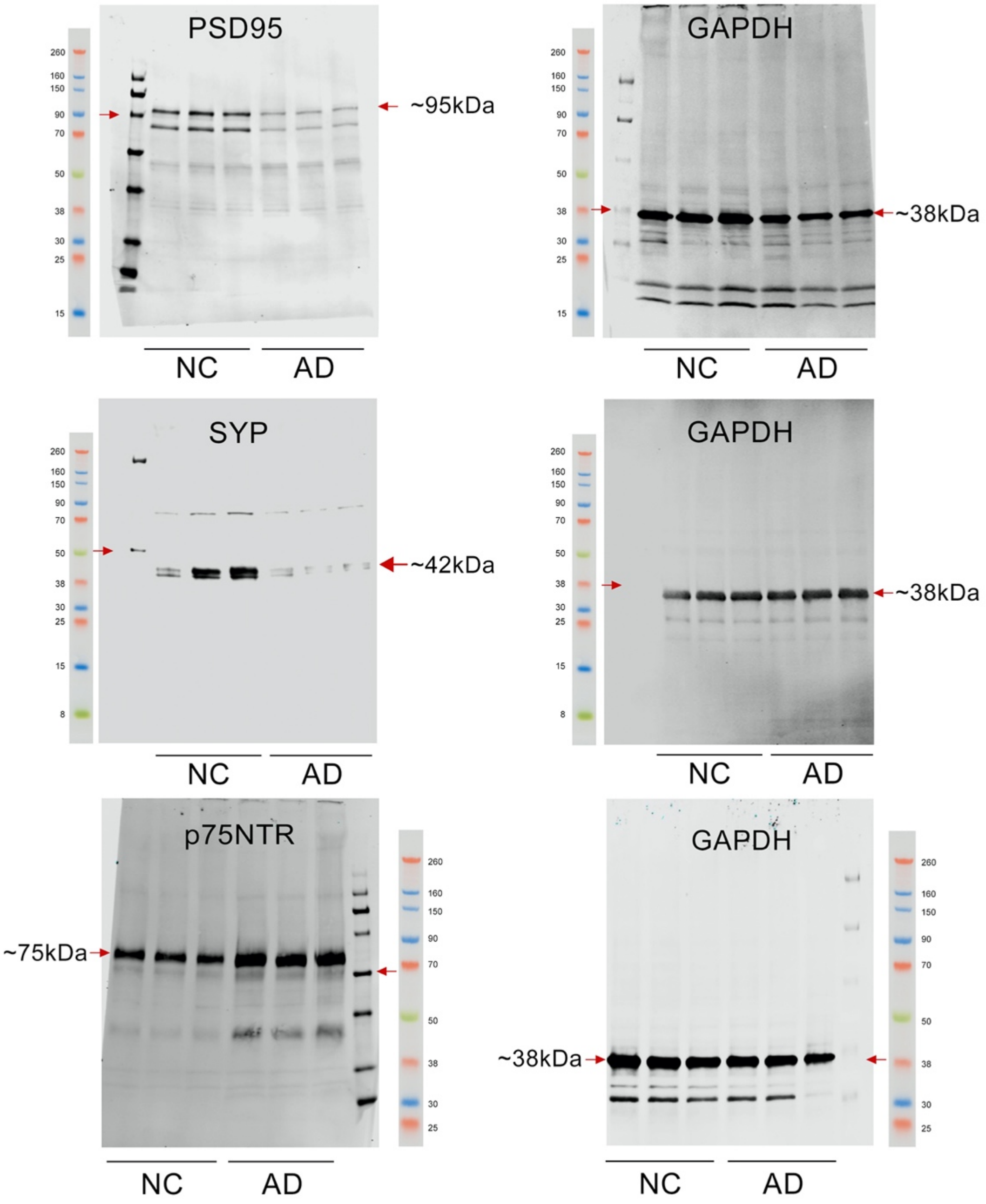


**Figure S4. Full western blots for retinal PSD95, Synaptophysin and p75NTR.**

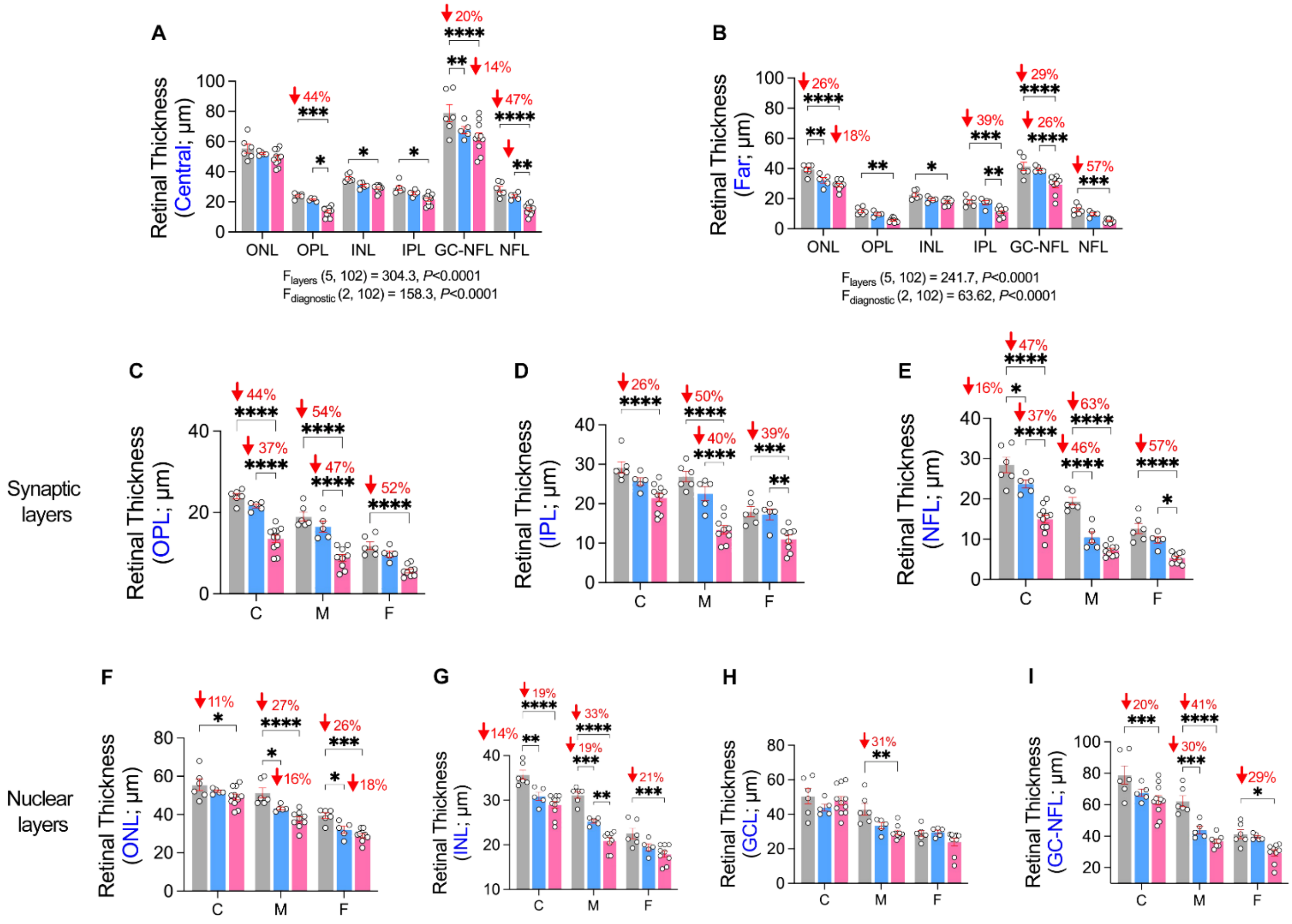


**Figure S5. Regional and spatial retinal atrophy assessed by retinal thickness.**

**(A–I)** Retinal thickness measurements across different regions and layers from a subset of individuals with NC (n =6), MCI (n = 5) and AD (n =9-11). Central (**A**) and far peripheral (**B**) total retinal thickness are shown. Thickness of specific retinal layers is shown for: OPL (**C**), IPL (**D**), NFL (**E**), ONL (**F**), INL (**G**), GCL (**H**), and GCL-NFL (**I**).

Data from individual subjects (circles) as well as group means ± SEMs are shown. *P < 0.05, **P < 0.01, ***P < 0.001, ****P < 0.0001, by two-way ANOVA followed by Tukey's post-hoc multiple comparison test. Percentage (%) changes are shown in red. NFL, Nerve fiber layer; GCL, ganglion cell layer; IPL, inner plexiform layer; INL, inner nuclear layer; OPL, outer plexiform layer; ONL, outer nuclear layer.

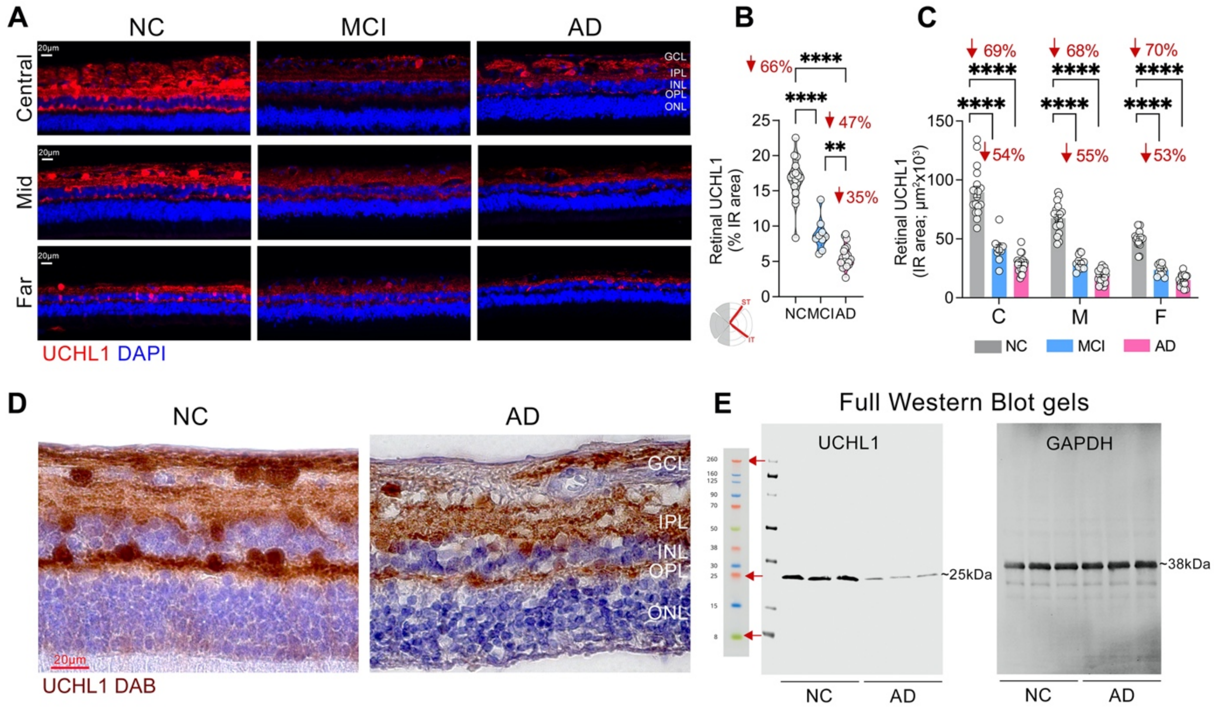


**Figure S6. Extended data on retinal UCHL1.**

(**A**) Representative fluorescence micrographs of retinal cross-sections immunolabeled for UCHL1 (red) and DAPI (blue) in patients with MCI and AD versus NC control. Scale bars: 20 µm.

(**B**) Quantitative analyses of retinal UCHL1 % IR area in MCI (n = 9) and AD-dementia (n = 18) subjects compared with age and sex-matched NC controls (n = 16).

(**C**) Quantitative analyses of retinal UCHL1 IR in predefined retinal subregions, central (C), mid- (M), and far-periphery (F), in patients with MCI (due to AD; n = 11) and AD-dementia (n = 21) as compared with age and sex-matched NC controls (n = 18).

(**D**) Representative micrographs of retinal UCHL1 using peroxidase-based 3,3′diaminobenzidine (DAB) and hematoxylin counterstaining. Scale bars: 20 µm.

(**E**) Full blots of retinal UCHL1 and GAPDH.

Violin plots show individual subjects (circles) with lower, median, and upper quartiles. Bar graph displays group means ± SEMs. *P < 0.05, ** P < 0.01, *** P < 0.001, **** P < 0.0001, by one- or two-way ANOVA followed by Tukey's post-hoc multiple comparison test. Percentage (%) changes are shown in red. GCL, ganglion cell layer; IPL, inner plexiform layer; INL, inner nuclear layer; OPL, outer plexiform layer; ONL, outer nuclear layer.

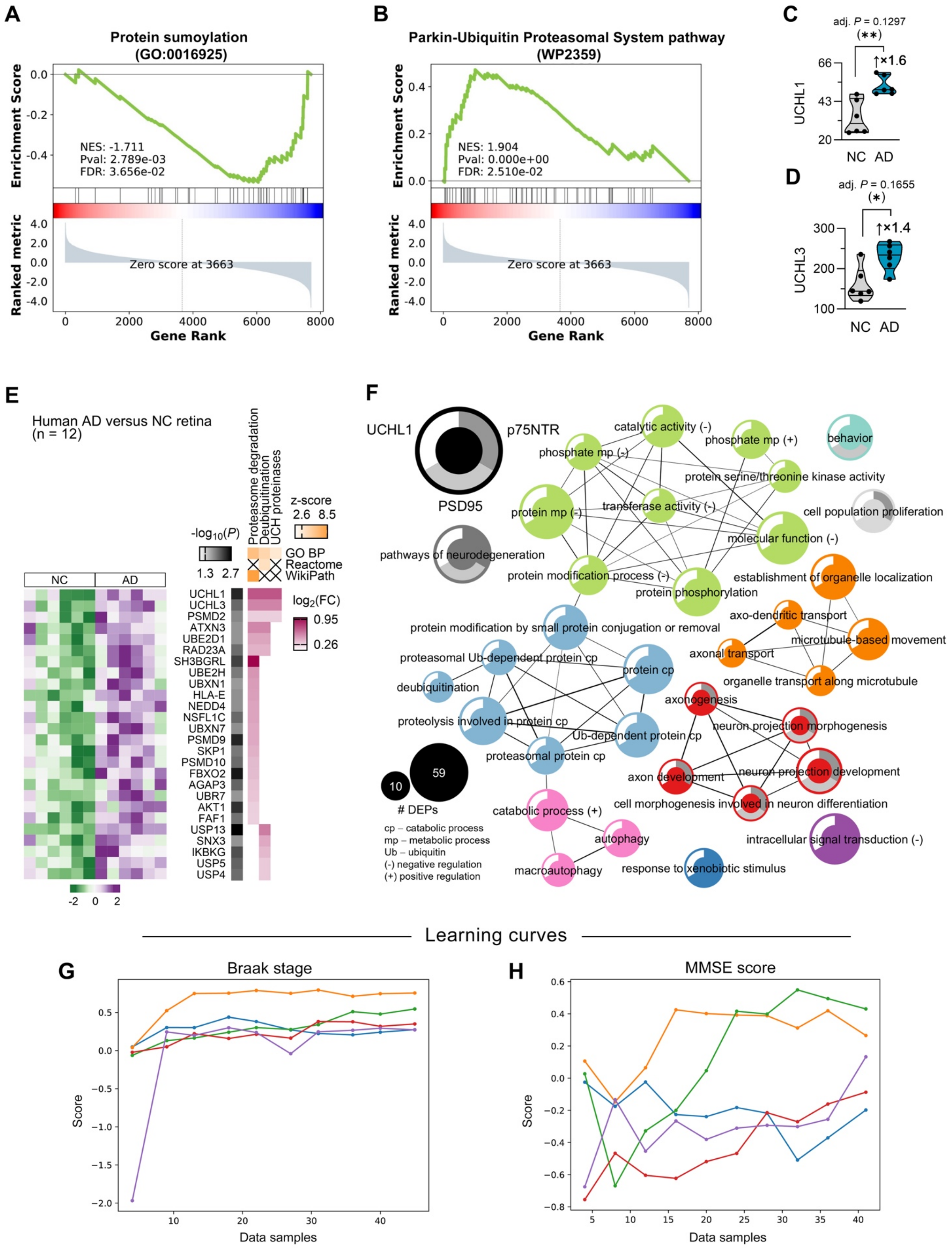

A
Protein sumoylation
(GO:0016925)
Enrichment Score
NES: -1.711
Pval: 2.789e-03
FDR: 3.656e-02
Ranked metric
Zero score at 3663
Gene Rank
B
Parkin-Ubiquitin Proteasomal System pathway
(WP2359)
NES: 1.904
Pval: 0.000e+00
FDR: 2.510e-02
C
adj. P = 0.1297
(**)
↑×1.6
UCHL1
NC AD
D
adj. P = 0.1655
(*)
↑×1.4
UCHL3
E
Human AD versus NC retina
(n = 12)
Proteasome degradation
Deubiquitination
UCH proteinases
z-score
-log10(P)
GO BP
Reactome
WikiPath
log2(FC)
UCHL1
UCHL3
PSMD2
ATXN3
UBE2D1
RAD23A
SH3BGRL
UBE2H
UBXN1
HLA-E
NEDD4
NSFL1C
UBXN7
PSMD9
SKP1
PSMD10
FBXO2
AGAP3
UBR7
AKT1
FAF1
USP13
SNX3
IKBKG
USP5
USP4
F
UCHL1
p75NTR
PSD95
pathways of neurodegeneration
catalytic activity (-)
phosphate mp (-)
phosphate mp (+)
behavior
protein serine/threonine kinase activity
protein mp (-)
transferase activity (-)
cell population proliferation
molecular function (-)
protein modification process (-)
protein phosphorylation
establishment of organelle localization
axo-dendritic transport
protein modification by small protein conjugation or removal
microtubule-based movement
axonal transport
proteasomal Ub-dependent protein cp
protein cp
organelle transport along microtubule
deubiquitination
axonogenesis
proteolysis involved in protein cp
neuron projection morphogenesis
Ub-dependent protein cp
proteasomal protein cp
axon development
neuron projection development
cell morphogenesis involved in neuron differentiation
catabolic process (+)
### DEPs
cp – catabolic process
mp – metabolic process
Ub – ubiquitin
(-) negative regulation
(+) positive regulation
autophagy
intracellular signal transduction (-)
macroautophagy
response to xenobiotic stimulus
Learning curves
G
Braak stage
H
MMSE score
Score
Data samples

**Figure S7. UCHL1 and associated molecular pathways in the AD retina.**

(**A-B**) GSEA enrichment plots for GO protein sumoylation enriched in our bottom-ranked dataset (downregulated in AD: NES=-1.711, FDR<0.05) (**A**) and for WikiPathway Parkin-ubiquitin proteasomal system enriched in our top-ranked dataset (upregulated in AD: NES=1.904, FDR<0.05) (**B**).

(**C-D**) Violin plots of the ubiquitin hydrolases, UCHL1 (**C**) and UCHL3 (**D**), quantified by MS in AD versus NC control retinas.

(**E**) Heatmaps of DEPs [log2(FC) and -log10(P)] in AD retina for select pathways linked to deubiquitination. The heatmap on the left corresponds to the protein expression level in the 6 NC individuals and the 6 AD patients, standardized by unit variance scaling and generated in ClustVis. Clustering of DEPs was carried out manually based on their involvement in select pathways for visual clarity.

(**F**) Metascape network of dysregulated UCHL1-associated pathways in the AD retina. The size of the nodes represents the number of DEPs, while the thickness of edges represents the overlapping DEPs (association score) between GO terms. The inner ring indicates the co-involvement of PSD95 and p75NTR with UCHL1 in the molecular pathways. DEPs: |FC|>1.2 and P<0.05.

(**G-H**) Learning curves for Braak stage (**G**) and MMSE (**H**) produced from Random Forest machine learning prediction.

Reference

1. Koopmans, F., van Nierop, P., Andres-Alonso, M., Byrnes, A., Cijsouw, T., Coba, M.P., Cornelisse, L.N., Farrell, R.J., Goldschmidt, H.L., Howrigan, D.P., et al. (2019). SynGO: An Evidence-Based, Expert-Curated Knowledge Base for the Synapse. Neuron *103*, 217-234 e214. 10.1016/j.neuron.2019.05.002.
